# Super Recombinator (SuRe): An *in vivo* recombination system for scalable and efficient transgene assembly at a single genomic locus

**DOI:** 10.1101/2025.04.15.646138

**Authors:** Junjie Luo, Cheng Huang, Caitlin Ann Taylor, Jane Li, Seung Je Woo, Chenghao Yu, Kang Shen, Mark J. Schnitzer

**Affiliations:** James Clark Center, Stanford University, Stanford, CA, 94305, USA; Howard Hughes Medical Institute, Stanford University, Stanford, CA, 94305, USA; Dept. of Biology, Stanford University, Stanford, CA, 94305, USA; CNC Program, Stanford University, Stanford, CA, 94305, USA; Dept. of Applied Physics, Stanford University, Stanford, CA, 94305, USA; Dept. of Neurosurgery, Stanford University, Stanford, CA, 94305, USA

## Abstract

The capacity to engineer organisms with multiple transgenic components is crucial to synthetic biology and basic biology research. For the former field, transgenic organisms allow the creation of novel biological functions; for the latter, such organisms provide potent means of dissecting complex biological pathways. However, the size limitations of a single transgenesis event and challenges associated with the assembly of multiple DNA fragments hinder the efficient integration of multiple transgenes. To overcome these hurdles, here we introduce a building block for synthetic design termed an integrated genetic array (IGA), which incorporates all genetic components into a single locus to prevent their separation during genetic manipulations. Since the natural recombination rate for genes located in the same locus is near zero, to construct IGAs we developed the Super Recombinator (SuRe) system, which uses CRISPR/Cas9, alone or in combination with site-specific serine recombinases, for *in vivo* transgene recombination at a single genomic locus. SuRe effectively doubles the number of elements assembled in each recombination round, exponentially accelerating IGA construction. By preventing the separation of transgenic elements, SuRe greatly reduces screening burdens, as validated here through studies of *Drosophila melanogaster* and *Caenorhabditis elegans*. To optimize SuRe, we compared CRISPR/Cas9-induced homology-directed recombination to site-specific recombination using various serine recombinases. Optimized versions of SuRe achieved efficiency and fidelity values near their theoretical maxima and allowed the generation of recombinant products up to 4.2 Mbp in size in *Drosophila*. Using SuRe, we created fruit flies with 12 transgenic elements for fluorescence voltage imaging of neural activity in precisely defined cell-types. Mathematical modeling of the scalability of SuRe to large transgene assemblies showed that integration times and gene assembly workloads respectively scale logarithmically and linearly with the number of transgenes, both major improvements over conventional approaches. Overall, SuRe enables the efficient integration of multiple genes at individual loci, up to the chromosomal scale.

## Introduction

Synthetic biologists seek to precisely engineer biological systems for the purpose of achieving novel biological functions^1,2^. This pursuit often requires modifying organisms via the introduction of multiple interacting multiple genes, regulatory elements, or other functional modules^3^. The subfield of synthetic genomics focuses on genome construction, which requires the assembly of a large number of DNA components into chromosome-scale structures^4–6^. More broadly, many biomedical experiments require the integration of multiple driver and effector components into a host organism for the purpose of dissecting complex biological functions.

Although transgenesis allows the introduction of foreign DNA into an organism’s genome, the size of a DNA fragment that can be reliably integrated within a single recombination step is limited, typically ranging from a few kilobases (kb) to 20–100 kb, depending on the transgenesis system^7–11^. For instance, in *Drosophila melanogaster*, the CRISPR/Cas9 system generally allows transgene knock-ins that are 1-10 kb long^12^, whereas the ΦC31 integrase can handle insertions up to 100 kb^13^. These size limitations imply that larger genetic fabrications require the combined assembly of multiple, smaller transgenic components.

A common way to combine multiple transgenic elements involves integrating them at separate chromosomal locations and then combining them through meiotic recombination. However, this strategy is constrained by the basic principles of inheritance. Based on Mendel’s law of independent assortment and Morgan’s law of linkage and crossing, the probability that two genes at different loci will recombine equals the probability that they will dissociate^14–20^. Hence, attempts to add genetic elements with high efficiency by integrating a distal component also raises the risk of the newly added element dissociating from the assembly. As the number of transgenes in an assembly increases, the number of potential recombination products grows exponentially. This combinatorial complexity makes identifying and isolating the correct combination exceedingly difficult, especially in polyploid organisms. Moreover, the availability of suitable genomic sites for precise transgene insertion is limited, as these sites must be carefully chosen to avoid disrupting endogenous gene function.

Another common strategy for integrating multiple transgenic components into the genome involves sequential integration via genome editing or manipulation, iteratively introducing each component, usually into the same genomic site^4,21–23^. This approach is constrained by its dependence on the successful completion of each successive integration step, making it a time-consuming process. The time required scales linearly with the number of genetic components that must be combined, which is especially problematic for organisms with long life cycles.

To overcome the challenges of identifying correctly recombined products in meiotic recombination-based approaches, we propose the ’Integrated Genetic Array’ (IGA) design principle. An IGA consolidates all transgenic components within a single genomic locus and functions as an indivisible unit. This drastically simplifies screening, as all desired components are either present together or absent entirely. Although sequential integration offers a route to build IGAs, it is time-consuming. Thus, we developed the Super Recombinator (SuRe) system to speed IGA construction by facilitating efficient *in vivo* recombination of transgenes at a single genomic locus.

SuRe constitutes a versatile toolkit that leverages CRISPR/Cas9 and site-specific recombination technologies. It operates by inserting reciprocally designed adaptor cassettes both upstream and downstream of target genes located at the same locus. These cassettes then mediate recombination between the target genes. SuRe offers several advantages over conventional methods. First, SuRe allows iterative recombination at a single genomic locus; this in turn substantially reduces screening efforts, since the assembled components do not segregate, and also maximizes the use of valuable transgene docking sites. Second, the system uses fluorescent markers to streamline the screening process; moreover, each recombination cycle yields a marker-free product, ensuring a clean integration and enabling iterative cycles of assembly. Third, SuRe exhibits broad compatibility, as it requires only a dedicated pair of recombinators for any set of transgenes sharing a common genomic locus and flanking sequences, where each recombinator adds a corresponding adaptor cassette to the target transgenes. Finally, and perhaps most importantly, because SuRe enables parallel *in vivo* recombination at a single locus, assembly times scale logarithmically with the number of transgenes. This is a major improvement over the linear scaling of sequential integration, and the efficiency gain becomes increasingly significant as more transgenes are integrated together.

Here, we document the capability of the SuRe system to recombine transgenes at a single locus in both *Drosophila melanogaster* and *Caenorhabditis elegans*. We evaluated CRISPR/Cas9-mediated homology-directed repair and site-specific recombination using a panel of serine recombinases within the SuRe system framework. This led to optimized versions of SuRe that exhibit high efficiency and fidelity in recombining transgenes and can generate megabase-scale recombinant products in *Drosophila*. We also showcase the practical utility of SuRe by using it to enhance neural imaging studies, including multi-color voltage imaging and Ca^2+^ imaging in multiple neuron-types. Finally, mathematical modeling of SuRe and conventional methods for large-scale transgene assembly highlights the substantial advantages of SuRe over existing approaches.

## Results

### Design of the SuRe system for recombination of transgenes at a single genomic locus

Constructing an IGA requires combining multiple transgenic components at a single genomic locus. Natural meiotic recombination at a single locus is extremely rare, making this a challenging task. The Super Recombinator (SuRe) system overcomes this limitation and uses two strains, Recombinator 1 (*R1*) and Recombinator 2 (*R2*), to integrate transgenes. The integration process has two steps: 1) Adaptor insertion: *R1* and *R2* strains facilitate insertion of reciprocally designed adaptor cassettes, *AD1* and *AD2*, upstream and downstream, respectively, of two target transgenes located at a single locus; and 2) Adaptor-mediated recombination: the adaptors mediate recombination between the transgenes, generating an IGA (**Figure 1A**). Iterative cycles of adaptor insertion and recombination exponentially increase the number of components in the IGA.

**Figure 1.**
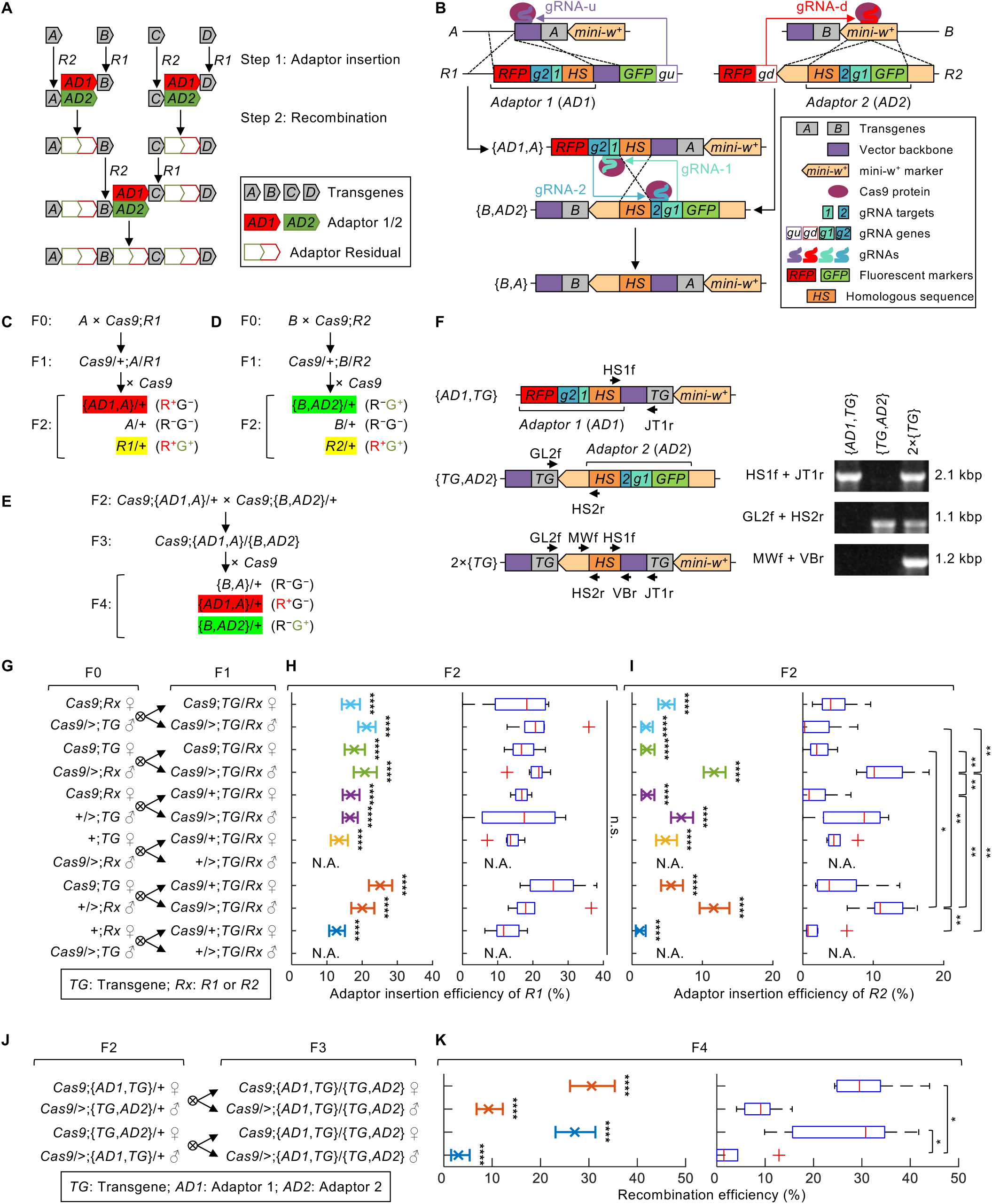
The SuRe system mediates recombination of transgenes at a single genomic locus. **(A)** Schematic of SuRe’s operational principles. Two strains, Recombinator 1 (*R1*) and Recombinator 2 (*R2*), recombine transgenes at the same genomic locus in two steps: 1) *R1* facilitates insertion of the adaptor 1 (*AD1*) cassette upstream of one transgene, and *R2* integrates the adaptor 2 (*AD2*) cassette downstream of the other transgene, generating transgene-adaptor strains; 2) The two reciprocally designed adaptors, *AD1* and *AD2*, mediate the recombination to generate an IGA. Repeated cycles of adaptor insertion and recombination exponentially expand the number of transgenes in the IGA. Elements in the diagram are not drawn to scale. **(B)** Schematic of the molecular design and working mechanism for SuRe using CRISPR/Cas9 for both adaptor insertion and recombination. Transgenes *A* and *B* are located at the same chromosomal locus and share identical flanking sequences, a portion of the plasmid vector backbone sequence upstream and a *mini-w^+^* marker downstream. The recombinator pair *R1* and *R2* strains are engineered reciprocally. Each carries a Cas9 protein expressed under a pan cell promoter (shown as Cas9 protein expression in the figure) and a genetic segment comprising three major elements: a *U6:3-gRNA-u* or *U6:3-gRNA-d* expressing a guide RNA that targets upstream transgene *A* or downstream transgene *B*, an *AD1* or *AD2* adaptor cassette, and a green or red fluorescent protein (GFP or RFP) marker that is distinct from the fluorescent marker in the adaptor cassette. The *AD1* and *AD2* adaptors are designed reciprocally, each with flanking homologous sequences for insertion upstream or downstream of the transgene, and each carries a gRNA that targets the other adaptor. *R1* facilitates the formation of a Cas9/gRNA-u complex, which creates a double-strand break upstream of transgene *A*, enabling *AD1* insertion via homology-directed repair. Likewise, *R2* enables *AD2* insertion downstream of transgene *B*. Crossing the resulting transgene-adaptor strains {*AD1*,*A*} and {*B*,*AD2*} brings *AD1* and *AD2* together, enabling their reciprocal gRNAs to target each other. This induces double-stranded breaks near the homologous sequence (HS) in the adaptors, leading to recombination. The recombination product {*B*,*A*} contains the original flanking sequences of transgenes *A* and *B*. This allows the same *R1* and *R2* pairs to be used for further rounds of recombination and expansion of the IGA. Reciprocal fluorescent markers in *R1*, *R2*, *AD1*, and *AD2* allow easy screening for the desired genotypes at each step. The vector backbone is a remnant of the transgenic generation process. Diagram elements are not drawn to scale. This working mechanism for SuRe applies to the experimental details and results presented in Figures 1C**–K****, 2, S1–3**. The legend also applies to panel **F.** Colored rectangles and pentagons represent distinct genetic elements. Magenta ovals denote the Cas9 protein, and thick curves depict gRNAs, of which the colors match those of their corresponding targets and expression genes. **(C–E)** Genetic crosses for SuRe-mediated recombination. **(C)** Transgene *A* is crossed with the recombinator strain *Cas9*;*R1*. The F1 transheterozygote *Cas9*/*+*;*A*/R1 is then crossed with a *Cas9* strain to produce F2 flies. Among F2 offspring, the desired {*AD1*,*A*} strain exhibit RFP but no GFP, whereas unreacted *R1* contains both RFP and GFP, and unreacted transgene *A* contains neither RFP nor GFP. **(D)** Transgene *B* is crossed with the recombinator strain *Cas9*;*R2*. The F1 transheterozygote *Cas9*/*+*;*B*/*R2* is then crossed with a *Cas9* strain to produce F2 flies. Among the F2 offspring, the desired {*B*,*AD2*} strains have GFP but no RFP, whereas unreacted *R2* contains both RFP and GFP, and unreacted transgene *B* contains neither RFP nor GFP. **(E)** *Cas9*;{*AD1*,*A*}/+ and *Cas9*;{*B*,*AD2*}/+ from F2 in **(C)** and **(D)** are crossed together to generate the F3 transheterozygote *Cas9*;{*AD1*,*A*}/{*B*,*AD2*}. The transheterozygote is then crossed with a *Cas9* strain for F4. Among F4 offspring, the desired {*B*,*A*} strain lacks both RFP and GFP, while unreacted {*AD1*,*A*} or {*B*,*AD2*} contain RFP or GFP. **(F)** PCR Confirmation of the adaptor insertion and the transgene recombination for duplicating an example transgene (*TG*: *82C10-LexA*). Primers HS1f (on *AD1*) and JT1r (on the transgene) were used to confirm *AD1* insertion. Primer HS2r (on *AD2*) and GL2f (on the transgene) were used to confirm *AD2* insertion. Primers VBr (on the *AD1*) and MWf on the *AD2* were used to confirm the recombination that generates a duplication of *TG*, *2×*{*TG*}. **(G–I)** Experimental design and results for studies of adaptor insertion efficiency for Recombinators *R1* and *R2*. **(G)** Cross scheme for assaying adaptor insertion efficiency. We assayed the insertion of *AD1* or *AD2* into an example transgene (*TG*: *82C10-LexA*). To account for potential maternal effects on Cas9 and gRNA expression that may influence the efficiency of *R1* and *R2*, we conducted all possible reciprocal crosses between F0 parents to generate transheterozygotes *TG/R1* or *TG/R2*. 10 of the 12 F1 transheterozygous genotypes were crossed with the Cas9 strain for F2, whereas 2 male genotypes without Cas9 expression were omitted (marked as N.A.). **(H, I)** We calculated the adaptor insertion efficiency for *R1* or *R2* as the percentage of desired transgene-adaptor animals {*AD1*,*TG*} or {*TG*,*AD2*} among all F2 progeny, respectively. *Left panels*, Mean ± 95% C.I. weighted average adaptor insertion efficiencies, calculated here and in all subsequent figures using the Clopper-Pearson method. There were at least five replicates per cross (n = 5–13, each with a parent of indicated F1 genotype crossed to a Cas9 strain). The mean efficiency of *R1* was ∼13–25% and that of *R2* was ∼1–12%, values that were significantly higher than the near-zero efficiency of natural recombination (**Methods**) for generating the transgene-adaptor strain (binomial test; ****: *P* < 10^-6^). *Right panels*, Box-and-whisker plots for efficiency values measured from individual F1 animals (parents of F2). We compared adaptor insertion efficiencies across different cross designs wit a Kruskal-Wallis ANOVA followed by Tukey’s honestly significant difference (HSD) test (*: *P* < 0.05; **: *P* < 0.01, n = 5–13). Here, and in all subsequent box-and-whisker-plots in the paper, red central lines denote median values, boxes span the 25th–75th percentiles, whiskers extend to 1.5 times the interquartile range, and red ‘+’ symbols mark outlier data points. **(J, K)** Experimental design and results for studies of adaptor-mediated recombination efficiency. **(J)** Cross scheme for assaying adaptor-mediated recombination efficiency. To account for potential maternal effects on Cas9 and gRNA expression, we performed reciprocal crosses between F2 flies carrying the transgene-adaptor {*AD1*,*TG*} and {*TG*,*AD2*} (*TG*: *82C10-LexA*). We crossed the resulting F3 transheterozygote {*AD1*,*TG*}/{*TG*,*AD2*} males and females with the Cas9 strain for F4. **(K)** Efficiency of adaptor-mediated recombination using each cross strategy of **(J)**. We determined recombination efficiency as the percentage of the desired animals carrying the IGA *2×*{*TG*} among all F4 progeny. *Left panels,* Mean ± 95% C.I. weighted average recombination efficiency (Each cross had n = 5 replicates, each with a parent of indicated F3 genotype crossed to a Cas9 strain). The average recombination efficiency was ∼27–31% for females and ∼3–9% for males, both higher than the near-zero rates from natural recombination (binomial test; ****: *P* < 10^-6^). *Right panels,* Recombination efficiencies measured from individual F3 animals (parents of F4). Some recombination efficiencies among different cross designs were significantly different (Kruskal-Wallis one-way ANOVA followed by Tukey’s HSD test; *: *P* < 0.05); non-significant comparisons between crosses (n.s.: *P* > 0.05) are not explicitly marked on the figure.

We engineered SuRe in *Drosophila melanogaster* by using CRISPR/Cas9-mediated genome editing to precisely insert adaptor cassettes adjacent to target transgenes and induce recombination between them (**Figure 1B**). Our design targets the Janelia GAL4 and LexA stock collections^24^, in which each transgene is flanked upstream by a transgenic vector backbone sequence and downstream by a mini-white marker, which enables an eye pigmentation phenotype (**Figure S1A**). Because the Janelia split GAL4 collection^25^, the CEP lines^26^, the Vienna Tile GAL4 (VT-GAL4) library^27,28^, and UAS and LexAop lines that use vectors from the Janelia Research Campus^29^ all share the same backbone vector, SuRe is compatible with the use of lines from any of these libraries.

We designed the Recombinators *R1* and *R2* strains to have a reciprocal relationship. Both *R1* and *R2* carry a Cas9 expressed via a pan-cell promoter and a genetic segment with three major elements. Specifically, this segment includes: 1) A gRNA (*gu*) targeting the transgene’s upstream flanking sequence, which is a residual sequence from the transgenic vector common to all lines in a given library; 2) A *3×P3-GFP* eye-specific green fluorescent protein (GFP) marker; and 3) An adaptor cassette, *AD1*, which contains a *3×P3-RFP* eye-specific red fluorescent protein (RFP) marker along with other functional elements and which is flanked by homology arms that allow integration adjacent to the transgene (**Figures 1B** and **S1B**). The Recombinator *R2* strain includes fluorescent markers, distinct from those in *R1*, and carries the corresponding adaptor *AD2* allowing its integration downstream of the transgene **(Figures 1B** and **S1C**). Further, *AD1* (from *R1*) and *AD2* (from *R2*) are reciprocally designed and contain gRNA components that target each other, along with a shared homologous sequence (*HS*) to mediate recombination in the subsequent step (**Figure 1B**).

Specifically, *AD1* carries a gRNA (*g2*) targeting a unique sequence in *AD2*, whereas *AD2* contains a gRNA (*g1*) targeting a unique sequence in *AD1* (**Figure 1B**).

The CRISPR/Cas9-mediated insertion efficiency of a DNA fragment is reportedly enhanced when the donor DNA is in close proximity to the target site^30^. Thus, to maximize the efficiency of SuRe, we positioned the *R1* and *R2* strain elements, including the donor DNA components, at the same genomic locus as the target library transgenes. To recombine transgenes *A* and *B*, which share identical flanking sequences and are located at a single locus, we first performed adaptor insertion adjacent to each transgene. We crossed *R1* with the stain bearing transgene *A* to generate transheterozygote F1 progeny *A*/*R1* containing transgene *A* (RFP–, GFP–) and all *R1* components (RFP+, GFP+) (**Figure 1C**). In the germline of such F1 progeny, the Cas9/gRNA complex, expressed from the R1 components, creates a double-strand break (DSB) upstream of *A*. This DSB facilitates homology-directed repair, leading to insertion of the *AD1* cassette at the break site. Subsequently, we outcrossed *A*/*R1* to either a wildtype or *Cas9* strain and selected the F2 progeny {*AD1*,*A*} that carried transgene *A* with *AD1* inserted upstream and exhibited the RFP+, GFP– phenotype (**Figures 1B,C** and **S1D**). The RFP marker in the adaptor confirms the presence of *AD1*, and the external GFP marker enables selection of the {*AD1*,*A*} (RFP+, GFP–) by distinguishing it from the *R1* components (RFP+, GFP+). Similarly, the *R2* strain was crossed with transgene *B* to yield transheterozygous F1 *B*/*R2*. We then crossed these F1 with Cas9. Among the resulting F2, we selected {*B*,*AD2*} (GFP+, RFP–), in which *AD2* was downstream of transgene *B* (**Figures 1B,D** and **S1E**).

In the adaptor-mediated recombination step, we crossed {*AD1*,*A*} with {*B*,*AD2*} to obtain transheterozygote F3 progeny {*AD1*,*A*}/{*B*,*AD2*} with a RFP+, GFP+ phenotype (**Figure 1E**). In the germline of these F3, Cas9 expression activates the gRNAs encoded from the adaptors *AD1* and *AD2* to reciprocally induce DSBs in both adaptors. These DSBs, together with the shared homologous sequence present in both adaptors, facilitate homology-directed repair leading to the recombination between transgenes *A* and *B*, generating the desired {*B*,*A*} array product (**Figure 1B,E**). We crossed F3 with wild type to obtain F4 progeny. The desired genotype {*B*,*A*} exhibits a w+, RFP-, and GFP-phenotype, indicating successful recombination, the presence of both transgenes *A* and *B*, and the absence of *AD1* and *AD2* adaptors (**Figures 1B,E** and **S1F**).

The successful recombination produced a {*B*,*A*} array. To mitigate potential crosstalk between *A* and *B*, an insulator sequence was incorporated in the shared homologous sequence (*HS*) used for recombination. This insulator is strategically positioned between the two transgenes in the resulting {*B*,*A*}. Importantly, {*B*,*A*} is free of any residual gRNA sequences, gRNA target sites, or fluorescent markers (**Figure 1B**), and its flanking upstream and downstream sequences are identical to those of the original single transgene *A* and *B*. This allows {*B*,*A*} to be treated as a single unit in subsequent rounds of recombination using the same *R1* and *R2* strains to further expand the IGA. Notably, The IGA remains intact during backcrossing, greatly simplifying genetic background cleaning.

As an example, we performed SuRe-mediated recombination using *82C10-LexA* (denoted *TG* for simplicity, located in the *attP40* site) for both transgenes *A* and *B*. The resulting {*B*,*A*} array is thus a duplication of *TG*, denoted *2×*{*TG*}. We verified the desired strains by PCR genotyping (**Figure 1F**).

### Efficiency of adaptor insertion

To assess the SuRe system’s efficiency, we measured the efficiency of adaptor insertion adjacent to the target transgene. We define the percentage of {*AD1*,*A*} or {*B*,*AD2*} in F2 progeny as the adaptor insertion efficiency for recombinator strains *R1* or *R2*, respectively. To simplify the experiment, we examined the efficiency of inserting *AD1* or *AD2* to the same transgene *82C10-LexA* (*TG*) using recombinator *R1* (*Act5C-Cas9*;*R1*) or *R2* (*Act5C-Cas9*;*R2*) strains, respectively.

Adaptor insertion efficiency can potentially be influenced by maternal effects. Cas9 and gRNA expression (driven by pan-cell *Act5C* and *U6* promoters, respectively) may vary depending on whether these genes are inherited maternally or paternally. Maternal inheritance typically results in higher expression levels of Cas9 and gRNA during early embryo development, compared to paternal inheritance, for which expression is delayed^31^. To account for this potential source of variation, we conducted all possible reciprocal crosses between F0 parents, generating heterozygotes *TG*/*R1* and *TG*/*R2* F1 progeny with each parental combination (**Figure 1G**). We crossed these F1 progeny with a *Cas9* strain, yielding F2 animals for assessments of adaptor insertion efficiency (**Figure 1G–I**).

The value of adaptor insertion efficiency is important, because it determines the number of animals that must be screened to obtain the desired transgene with an integrated adaptor. For *Drosophila*, an efficiency >1% is usually considered acceptable, given the use of fluorescent markers to facilitate screening and feasibility of collecting hundreds of F2 progeny. Several key factors influence the insertion efficiency, including the length of the homology arms, the size of the adaptor to be inserted, and the cleavage efficiency of the Cas9/gRNA complex. In our initial design of the *R1* and *R2* recombinator strains, the homology arms for adaptor insertion were 1.0–1.8 kb in length, and the adaptors were ∼2.7 kb. *R1* achieved an adaptor insertion efficiency of ∼13–25% (**Figure 1G,H**), whereas the first version of *R2* had an efficiency of <1% (**Figure S1G–I**). To improve the performance of *R2*, we created a new version using a different gRNA and a correspondingly modified homology arm to match the new gRNA’s cleavage site. These changes to *R2* boosted adaptor insertion efficiency to ∼1–12% (**Figure 1G,I**). Although we observed substantial variations in insertion efficiency among different F1 progeny (the parents of F2) for the improved *R2* (**Figure 1I**), efficiencies for both versions of *R2* were much higher than the near-zero rate obtained by using natural integration to generate the transgene-adaptor strain (**Figures 1I** and **S1H**; *Statistical analysis* in **Methods**).

Since close proximity enhances recombination efficiency^30^, we positioned the recombinator components (excluding *Cas9*) at the same genomic locus as the target transgene. To facilitate transgene recombination at commonly used docking loci, we engineered three pairs of recombinator strains, with components respectively located at the *attP40*, *attP2*, or *VK27* sites. Although the experiments of **Figure 1G,I** used Recombinator components and transgenes integrated in *attP40*, our analysis revealed no significant differences in adaptor insertion efficiency across all three docking sites (**Figure S1M–O**)

### Efficiency of adaptor-mediated recombination

Following the generation of F2 transgene-adaptor strains {*AD1*,*TG*} and {*TG*,*AD2*}, to take maternal effects into account we performed reciprocal crosses between them to obtain the transheterozygote F3 {*AD1*,*TG*}/{*TG*,*AD2*} (**Figure 1J**). We crossed these F3 with a *Cas9* strain and evaluated the efficiency of adaptor-mediated recombination in the resulting F4 generation (**Figure 1J,K**). Recombination efficiency was defined as the percentage of F4 progeny harboring the desired *2×*{*TG*} array. *2×*{*TG*} lacked both *AD1* and *AD2*, resulting in a GFP–, RFP– phenotype that was distinguishable from undesired siblings. As with adaptor insertion, a recombination efficiency >1% is generally sufficient for practical applications. Our results met this criterion, with male parents having recombination efficiencies of ∼3–9% and female parents from ∼27–31% (**Figure 1K**). Importantly, all measured recombination efficiencies were significantly higher than the near-zero rate of natural recombination rate (**Figure 1K**; **Methods**). Further, when transgenes were integrated at *attP2* and recombined using their corresponding SuRe Recombinators, also located at *attP2*, the recombination efficiency remained above 1% (**Figure S1P–R**).

In our design for adaptor-mediated recombination, each adaptor (*AD1* and *AD2*) contains a gRNA that reciprocally targets its partner, leading to two Cas9-mediated DSBs, one in each adaptor, during the recombination process. This dual-DSB strategy allows efficient and accurate repair using the flanking sequences containing the transgene as templates, maximizing generation of the desired recombinant product {*B*,*A*} (**Figure 1B**). We investigated whether a single DSB sufficed to trigger the recombination to produce desired products. We generated transgenes integrated with mutated adaptors ({*AD1\**,*TG*} and {*TG*,*AD2\**}) whose gRNA target sites were disrupted (**Figure S2A**, see **Methods** for details on mutant adaptor strain generation). These mutations prevented Cas9 cleavage at the respective adaptors during recombination.

By crossing strains carrying these mutated adaptors with the original transgene-adaptor strains ({*AD1*,*TG*} and {*TG*,*AD2*}), we created scenarios with zero, one, or two DSBs (**Figure S2B**). Analysis of recombination efficiency under these different conditions revealed that recombination occurred with at least one functional DSB, although efficiency was sometimes reduced (**Figure S2C–F**). Therefore, to ensure optimal performance and minimize potential variability associated with individual gRNA cutting efficiency, we retained the two-DSB design in the SuRe system.

### Fidelity of the SuRe system

Adaptor insertion and the subsequent adaptor-mediated recombination generate the desired IGA while retaining the flanking sequences of the original transgene (**Figure 1B**), thereby enabling iterative use of the same *R1* and *R2* for further rounds of adaptor insertion and recombination and hence exponential expansion of the transgene assembly (**Figure 2A**).

**Figure 2.**
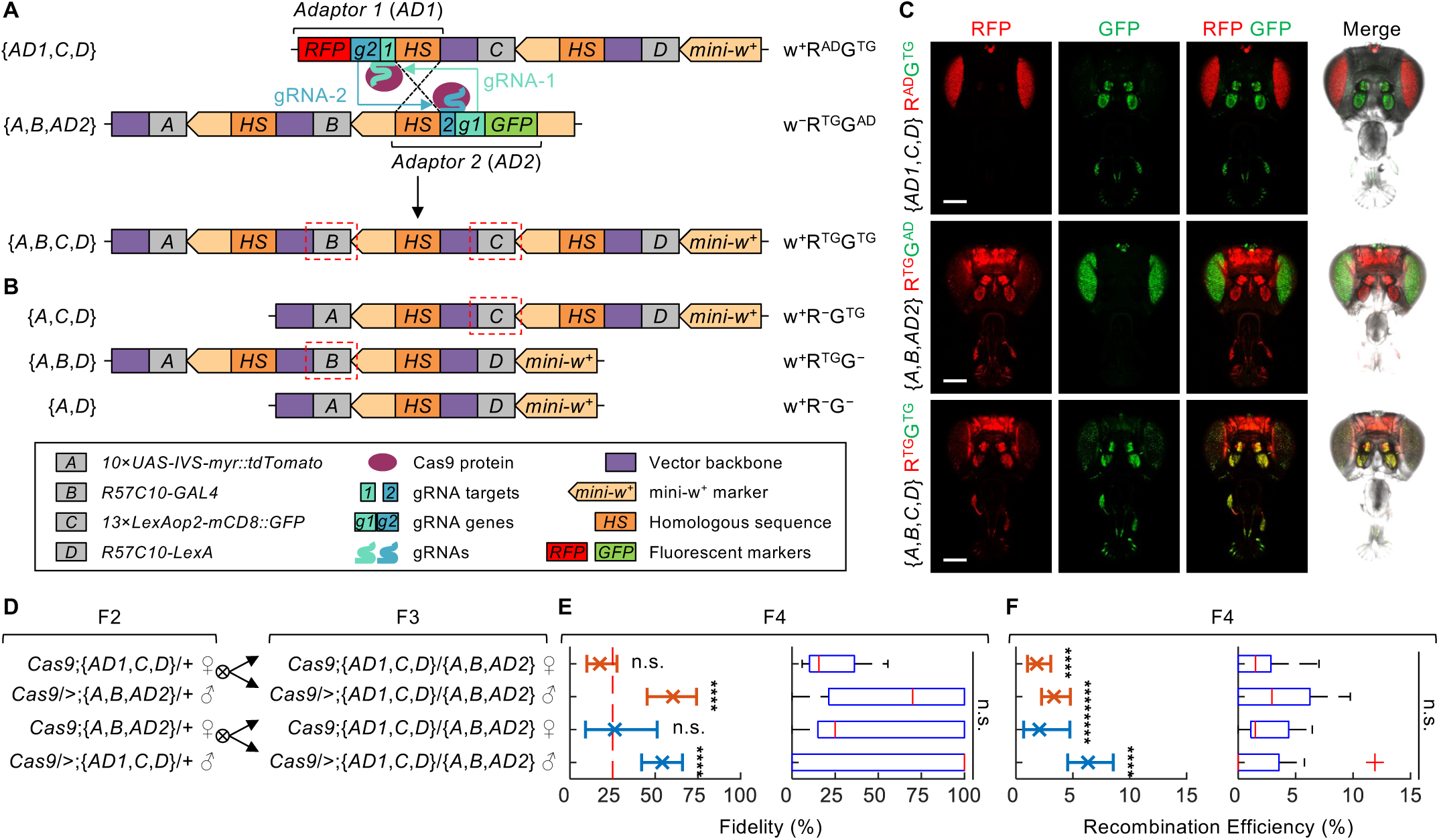
Recombination fidelity of the SuRe system. **(A, B)** Schematics depict the potential products resulting from subsequent rounds of recombination with Integrated Gene Arrays (IGAs), illustrating the desired outcomes and possible byproducts that arise from alternative recombination events. Individual transgenes *A*, *B*, *C*, and *D* are at the same chromosomal locus and share identical flanking sequences. {*AD1*,*C*,*D*} and {*A*,*B*,*AD2*} are results from insertion of adaptors to the previous integrated {*C,D*} and {*A,B*} arrays, respectively. {*AD1*,*C*,*D*} and {*A*,*B*,*AD2*} both contain two identical homologous sequences (*HS*). Products from the recombination between {*AD1*,*C*,*D*} and {*A*,*B*,*AD2*} were identified after an initial screening for the absence of eye-specific RFP (R^AD^) and GFP (G^AD^) markers (from the adaptors) and the presence of mini-w^+^ (from the IGA) was applied. The desired product {*A*,*B*,*C*,*D*} is generated from the desired pairing of *HS*s, **(A)**, while three possible undesired products result from alternative pairings of *HS*s, **(B)**. **Figure S3A–C** has further details about the undesired pairing scenarios. Colored rectangles and pentagons represent distinct genetic elements. Magenta ovals denote the Cas9 protein, and thick curves depict gRNAs, of which the colors match those of their corresponding targets and expression genes. Red dashed boxes highlight a key criterion for genotyping: only the desired product retains both transgenes flanking the recombination site (transgene *B* and *C*), whereas undesired products lose at least one of the two genes. After an initial screening using visible markers (selecting for w^+^R^AD–^G^AD–^ flies), PCR genotyping for the presence of transgenes *B* and *C* identifies the desired recombination product. The use of example transgenes *A*–*D* with readily observable expression patterns allows the detection of the desired product by fluorescence microscopy, rather than relying solely on PCR genotyping. This approach streamlines the measurements of recombination fidelity. Example transgenes used for fidelity measurements are: *A*: *10×UAS-IVS-myr::tdTomato*, *B*: *R57C10-GAL4*, *C*: *13×LexAop2-mCD8::GFP*, and *D*: *R57C10-LexA.* Screening markers: w^+^: eye-pigmentation marker; R^AD^/G^AD^: eye-specific RFP/GFP markers from adaptors; R^TG^/G^TG^: pan-neuronal RFP/GFP markers from transgenes; R^−^: no RFP marker; G^−^: no GFP marker. **(C)** Expression patterns of fluorescent proteins in the indicated genotypes. The *R57C10* promoter drives expression in the brain and antenna; the *3×P3* promoter (found in the markers in *AD1* and *AD2*) drives expression in the eyes. These distinct expression patterns allow visual differentiation of desired {*A*,*B*,*C*,*D*} and other recombination products and unreacted transgenes with adaptors. Transgene strains A, B, C and D are detailed in **(A, B)**. R^AD^/G^AD^: eye-specific RFP/GFP markers from adaptors. R^TG^/G^TG^: pan-neuronal RFP/GFP markers from transgenes. Scale bars: 200 µm. **(D–F)** Experimental design and results for studies of adaptor-mediated recombination efficiency and fidelity for two IGAs. **(D)** Cross scheme for assaying recombination between two IGAs. Note that F0 and F1 crosses for adaptor insertion are omitted. To account for possible maternal effects on Cas9 and gRNA expression, we performed reciprocal cross between the F2 flies carrying the {*AD1*,*C*,*D*} and {*A*,*B*,*AD2*} adaptors. F3 transheterozygote *Cas9*;{*AD1*,*C*,*D*}/{*A*,*B*,*AD2*} males and females were then crossed with Cas9 for F4. Transgene strains *A*, *B*, *C* and *D* are as described in **(A, B)**. **(E)** Fidelity of recombination between two IGAs in the corresponding cross scheme shown in **(D)**. We determined recombination fidelity as the percentage of desired animals {*A*,*B*,*C*,*D*} (phenotype w^+^R^TG^G^TG^) among the F4 progeny resulting from the initial screening for the absence of both adaptor markers R^AD^ and G^AD^ and the presence of the *mini-w*^+^ (phenotypes w^+^R^TG^G^TG^, w^+^R^−^G^TG^, w^+^R^TG^G^−^, and w^+^R^−^G^−^). *Left panels,* Mean ± 95% C.I. weighted average recombination fidelity (n = 5–14 replicates, each with a parent of the indicated F3 genotype crossed to a Cas9 strain). If the four recombination products occur with equal probability, the expected recombination fidelity is 25% (red dashed line). The recombination fidelity of females was not significantly different from 25%, whereas male F3 progeny had a significantly higher fidelity (binomial test; n.s.: *P* > 0.05, ****: *P* < 10^-6^), suggesting a bias towards the desired product in males. *Right panels,* Distributions of recombination fidelity measured for individual F3 animals (parents of F4). Recombination fidelities among different cross designs were not significantly different (Kruskal-Wallis ANOVA and Tukey’s HSD test; n.s.: *P* > 0.05). **(F)** Efficiency of recombination between two IGAs using the corresponding cross schemes of **(D)**, determined as the percentage of desired animals carrying {*A*,*B*,*C*,*D*} among all F4 progeny. *Left panels,* Mean ± 95% C.I. weighted average recombination efficiency (n = 5–14 replicates, each with a parent of the indicated F3 genotype crossed to a Cas9 strain). The average recombination efficiency was ∼2% for females and ∼3–6% for males, both significantly higher than that via natural recombination (binomial test; ****: *P* < 10^−6^). *Right panels,* Distributions of recombination efficiencies for individual F3 animals (parents of F4). Recombination efficiencies among different cross designs are not significantly different (Kruskal-Wallis ANOVA followed by Tukey’s HSD test; n.s.: *P* > 0.05).

When recombining four transgenes *A*, *B*, *C* and *D* using SuRe, we first generate two IGAs, {*A*,*B*} and {*C*,*D*}, through a round of adaptor insertion and recombination as discussed above. Although markers and gRNA target sequences are removed during the process, each round leaves a homologous sequence (*HS*) in the adaptor residual between the combined transgenes (**Figure 1B**). Consequently, {*A*,*B*} contains an *HS* between *A* and *B*, and {*C*,*D*} contains an *HS* between *C* and *D*. In the second round, inserting *AD1* and *AD2* to generate {*AD1*,*C*,*D*} and {*A*,*B*,*AD2*} introduces an additional *HS*. {*AD1*,*C*,*D*} and {*A*,*B*,*AD2*} now each contain two *HS*s (**Figure 2A**). These multiple *HS*s can potentially lead to alternative pairing scenarios during recombination, generating undesired products ({*A*,*C*,*D*}, {*A*,*B*,*D*}, and {*A*,*D*}) that are indistinguishable from the desired {*A*,*B*,*C*,*D*} based on adaptor markers alone (**Figures 2A, 2B** and **S3A–C**).

We define the recombination fidelity of the SuRe system as the proportion of desired products that are obtained after screening for the absence of adaptor markers. To directly measure the fidelity without laborious PCR genotyping, we performed two rounds of recombination using components of the UAS/GAL4 and LexA/LexAop binary expression systems. Specifically, we used the following four transgenes as *A*, *B*, *C*, and *D* above, respectively: *20×UAS-myr::TdTomato*, *R57C10-GAL4*, *13×LexAop2-mCD8::GFP* and *R57C10-LexA*. We first recombined *20×UAS-myr::Tdtomato* with *R57C10-GAL4* and *13×LexAop2-mCD8::GFP* with *R57C10-LexA*. Next, we recombined the resulting arrays to generate an assembly with all four transgenes (**Figure 2A**). The *R57C10* pan-neuronal driver, derived from *Drosophila nSyb*, was chosen for its ability to drive strong fluorescence expression that is readily detectable through the cuticle of the fly head. This allowed us to easily identify the desired product, as only the combination of *R57C10-GAL4* with *20×UAS-myr::TdTomato* produces pan-neuronal TdTomato expression, and the combination of *R57C10-LexA* and *13×LexAop2-mCD8::GFP* results in pan-neuronal GFP expression (**Figure 2C**).

When recombining the two IGA-adaptors {*20×UAS-myr::TdTomato,R57C10-GAL4*,*AD2*} and {*AD1*,*13×LexAop2-mCD8::GFP*,*R57C10-LexA*}, after selection for lack of adaptor markers, four potential products remain: {*20×UAS-myr::TdTomato*,*13×LexAop2-mCD8::GFP*,*R57C10-LexA*}, {*20×UAS-myr::TdTomato,R57C10-GAL4*,*R57C10-LexA*}, {*20×UAS-myr::TdTomato,R57C10-LexA*}, and the desired {*20×UAS-myr::TdTomato,R57C10-GAL4*,*13×LexAop2-mCD8::GFP*,*R57C10-LexA*} (**Figure 2A,B**). Assuming equal probability for each product, the expected fidelity of this recombination step would be 25%. However, we observed a fidelity of ∼18–26% for F3 females (F4 parents) and up to ∼54–60% for F3 males (F4 parents), indicating a sex-specific bias towards the desired product from F3 male parents (**Figure 2D,E**). We also measured the recombination efficiency, defined as the percentage of desired {*A*,*B*,*C*,*D*} progeny among all F4 offspring, which was ∼2% and ∼3–6% for F3 female and male parents, respectively (**Figure 2D,F**). All measured recombination efficiencies were significantly higher than the near-zero natural recombination rate. Since recombination fidelity is not 100%, PCR genotyping remains necessary to identify desired products, especially when recombination does not yield easily observable phenotypes. Efficient PCR screening can be achieved by verifying the presence of the two genes flanking the recombination site (Transgenes *B* and *C* in **Figure 2A,B**), rather than genotyping every gene in the assembly.

It is important to note that adaptor insertion into IGAs can also produce undesired outcomes due to alternative pairings of homology arms (**Figure S3D–G**). In these cases, after screening for the presence of the adaptor marker, PCR genotyping of the gene adjacent to the inserted adaptor (Transgene *D* in **Figure S3D**) is sufficient to identify the desired IGA-adaptor product, especially when no distinguishing markers in the IGA are available. We tested inserting *AD2* to {*A*,*B*,*C*,*D*} and observed a very high adaptor insertion fidelity of ∼95–96% and an insertion efficiency of ∼6–11% (**Figure S3H,J**). The high fidelity for adaptor insertion suggests that minimal PCR genotyping is required to verify successful insertion.

The key difference in fidelity between adaptor insertion and recombination concerns their requirements for DNA break repair. Adaptor insertion demands precise alignment of both ends of the Cas9/gRNA-induced double-strand break with the template (**Figure S3D**), whereas recombination can be initiated by a single double-strand break (**Figure S2**). This mechanism allows the released homologous sequence (HS) in the adaptor to recombine with residual HS between genes, resulting in undesired products (**Figure S3E–G**).

The fidelity of both adaptor insertion and recombination is crucial for estimating the number of strains requiring PCR genotyping. These estimations allow for a direct comparison of the genotyping workload between the SuRe system and traditional genetic approaches, as detailed below and in the **Supplemental Appendix**.

### Validating the SuRe system in *C. elegans*

To assess the generalizability of the SuRe system, we adapted it for use in *C. elegans* (**Figure 3A**). As a proof of concept, we targeted a single-copy transgene, *pdes-2::myr-mScarlet::let-858 3’UTR* (referred to as *TG*), that expresses myristoylated mScarlet in the nociceptive neurons PVD and FLP (**Figure 3A–C**). This transgene was inserted at a defined locus (**Figure 3B**) using CRISPR-Cas9 and confirmed by visual inspection and PCR (**Methods**).

**Figure 3.**
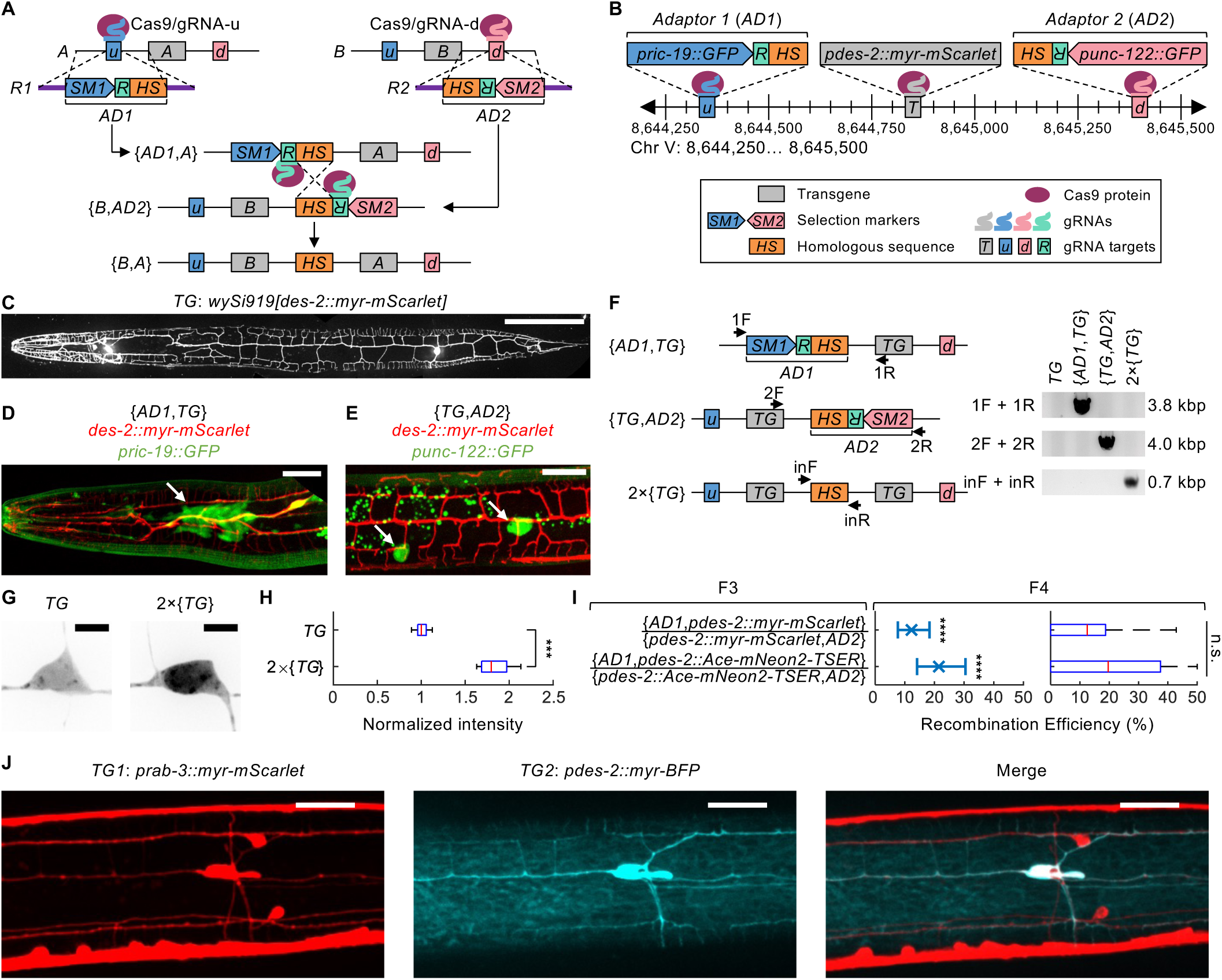
SuRe facilitates efficient transgene recombination in *C. elegans*. **(A)** Molecular design and mechanism of the SuRe system in *C. elegans*. SuRe delivers recombinator components Cas9 protein, gRNAs, and DNA templates containing adaptors through injection. The DNA templates are linearized double-strand DNA with homology arms flanking adaptors *AD1* or *AD2*. *AD1* and *AD2* are designed reciprocally, each with a distinct selection marker (*SM1* in *AD1* and *SM2* in *AD2*), the same gRNA targeting sequence and the same homologous sequences that mediate recombination. Transgenes *A* and *B* are located in the same chromosomal locus and share identical flanking sequences. In the adaptor insertion step, *AD1* is inserted upstream of transgene A, and *AD2* is inserted downstream of transgene B. This creates two transgene-adaptor strains, {*AD1*,*A*} and {*B*,*AD2*}. In the recombination step, {*AD1*,*A*} and {*B*,*AD2*} are crossed for the transheterozygote {*AD1*,*A*}/{*B*,*AD2*}. The Cas9/gRNA-R is injected into {*AD1*,*A*}/{*B*,*AD2*} to induce recombination. The desired recombination product {*B*,*A*} retains the original flanking sequences of individual transgenes A and B, and lacks the adaptor markers, allowing for repeated cycles of adaptor insertion and transgene recombination to construct larger IGAs. Elements in the diagram are not drawn to scale. This working mechanism for SuRe in *C. elegans* applies to the experimental details and results presented in subsequent panels of Figures 3. The legend also applies to panels **B** and **F**. Colored rectangles and pentagons denote distinct genetic elements. Magenta ovals denote the Cas9 protein, and thick curves depict gRNAs, of which the colors match those of their corresponding targets and expression genes. **(B)** Schematic representation of genetic compositions and insertion loci of the transgene *pdes-2::myr-mScarlet* (*TG*) and adaptor cassettes *AD1* or *AD2*. *wySi919[pdes-2::myr-mScarlet]* has a red fluorescent marker Scarlet inserted at the *oxTi365* safe harbor site on Chromosome V. *AD1* and *AD2* both contain the same 0.5 kbp homologous sequence (HS) and the same gRNA target sequence (R), enabling recombination at the target site when the Cas9/gRNA is present. However, they have distinct fluorescent markers: *pric-19::GFP* in *AD1* and *punc-122::GFP* in *AD2*, which have different expression patterns. *AD1* is inserted approximately 0.5 kbp upstream of the transgene *wySi919[pdes-2::myr-mScarlet]*, and *AD2* is inserted approximately 0.5 kbp downstream. The DNA sequence between the adaptor insertion site and the transgene insertion site serves as a homology arm in the *AD1* or *AD2* insertion process. **(C–E)** Expression patterns of fluorescent proteins in the indicated genotypes. **(C)** Fluorescence image showing the expression pattern of the transgene *wySi919[pdes-2::myr-mScarlet]* (*TG*). *TG* expresses a red fluorescent protein, Scarlet, in two pairs of highly-branched neurons, Posterior Ventral Process D (PVD) and FLaP-like Dendritic Ending (FLP). Scale bar: 100 µm. **(D)** The expression pattern of fluorescent proteins in {*AD1*,*TG*}. *TG* expresses Scarlet in multidendritic FLP neurons. *AD1* expresses a marker *pric-19::GFP* in the nerve ring (arrow). **(E)**The expression pattern of fluorescent proteins in {*TG*,*AD2*}.*TG* expresses Scarlet in multidendritic PVD neurons. *AD2* expresses a marker *punc-122::GFP* in coelomocytes (arrows). Scale bars in **D, E**: 20 µm. **(F)** PCR genotyping for indicated genotypes. *TG* is *wySi919[pdes-2::myr-mScarlet]*. {*AD1*,*TG*} carries a *TG* and an adaptor *AD1* inserted upstream of *TG*. {*TG*,*AD2*} carries the same *TG* and an *AD2* inserted downstream of *TG*. *2×*{*TG*} is a product resulting from recombination between {*AD1*,*TG*} and {*TG*,*AD2*}, bringing together two copies of the *TG*. PCR primers targeted specific regions flanking the *AD1* and *AD2* insertion sites and adaptor residuals. **(G, H)** Comparison of Scarlet expression in the single-copy transgene {*pdes-2::myr-mScarlet*} (*TG*) and the double-copy array *2×*{*pdes-2::myr-mScarlet*} (*2×*{*TG*}) in the PVD neuron soma. **(G)** Example confocal microscopy images showing Scarlet expression in the PVD neuron soma of *TG* and *2×*{*TG*} worms. Scale bar: 5 µm. **(H)** Scarlet fluorescence intensity values in the PVD neuron soma of *TG* and *2×*{*TG*} worms. Fluorescence intensity values were normalized to the median intensity for worms with the single-copy transgene (*TG*). Worms with the double-copy transgene (*2×*{*TG*}) exhibited significantly greater fluorescence intensities compared to those with the single-copy (Wilcoxon rank sum test; ***: *P* < 0.001; n = 12, 11 PVD neurons examined per genotype). **(I)** Efficiency of SuRe-mediated recombination in *C. elegans*, measured as the percentage of desired recombination products among all F1 offspring (progeny of injected transheterozygous). *Left panels,* Mean ± 95% C.I. weighted average adaptor insertion efficiency (n = 14, 21 replicates, each with an indicated F3 genotype worm that self-fertilized). The average efficiencies were ∼12–21%, significantly higher than that of the natural recombination (binomial test; ****: *P* < 10^-6^). *Right panels,* Recombination efficiencies measured from individual F3 animals (parents of F4) were not significantly different between different cross designs (Kruskal-Wallis one-way ANOVA followed by Tukey’s HSD test; n.s.: *P* > 0.05). **(J)** Confocal fluorescence images showing co-expression of *prab-3::myr-mScarlet* (red) and *pdes-2::myr-BFP* (blue) in the IGA strain *prab-3::myr-mScarlet*, *pdes-2::myr-BFP*, generated with SuRe by recombining *prab-3::myr-mScarlet* and *pdes-2::myr-BFP* at the same locus (see **B** for transgene insertion site). Scale bars: 20 µm.

In this adaptation, rather than relying on transgenic expression as in *Drosophila*, we delivered Cas9 and gRNAs via direct injection into *C. elegans*. We designed a pair of adaptor cassettes, *AD1* and *AD2*, each containing a distinct fluorescent marker for easy identification: *AD1* with *pric-19::GFP* expressed in the nerve ring and *AD2* with *punc-122::GFP* expressed in coelomocytes (**Figure 3A**). Both adaptors contained the same gRNA target sequence (**Figure 3A**). Using CRISPR/Cas9 for adaptor insertion, we generated two strains: {*AD1*,*TG*} with *AD1* inserted upstream of *TG*, and {*TG*,*AD2*}, with *AD2* inserted downstream (**Figure 3A,B,D,E**).

To test the recombination efficacy of SuRe, we crossed P0 {*AD1*,*TG*} with {*TG*,*AD2*} to generate trans-heterozygotes F1 {*AD1*,*TG*}/{*TG*,*AD2*}. We injected Cas9 and gRNAs into these F1 worms to promote recombination between the two *TGs*. Following injection, individual F1 animals were isolated and allowed to self-fertilize. Their F2 were screened visually for the absence of both *pric-19::GFP* and *punc-122::GFP* markers, indicating successful recombination. PCR genotyping was performed to confirm the presence of the desired *2×*{*TG*} recombinant product (**Figure 3F**).

Successful recombination produced a tandem duplication of the transgene, *2×*{*TG*}, with upstream and downstream flanking sequences identical to those of the original transgene and lacking any residual gRNA target sites. To assess expression levels, we measured myr-mScarlet fluorescence intensity and found that the *2×*{*pdes-2::myr-mScarlet::let-858 3’UTR*} array exhibited 1.8-fold greater brightness than the single-copy transgene (**Figure 3G,H**). This shows that SuRe can reliably generate IGAs of single-copy transgenes with predictable expression levels at a single defined locus in *C. elegans*. SuRe offers a substantial advantage over both traditional multi-copy extrachromosomal arrays, which often exhibit instability and high levels of expression variability^32–34^, and typical single-copy insertions, which can be limited by low expression.

We assessed the recombination efficiency of SuRe in *C. elegans* by generating duplication strains, specifically *2×*{*pdes-2::myr-mScarlet*} and *2×*{*pdes-2::Ace-mNeon2-TSER*} (**Figure 3I**). Approximately 70% of injected F1 trans-heterozygotes produced at least one offspring carrying the desired recombinant product. The overall recombination efficiency, measured as the percentage of desired recombinant offspring among all F2 progeny from injected transheterozygote F1s, ranged from ∼12–22% in these two examples (**Figure 3I**).

Next, we tested the SuRe system in recombining two distinct transgenes with expression in different tissues: *prab-3::myr-mScarlet* (with pan-neuronal expression of myr-mScarlet) and *pdes-2::myr-BFP* (with PVD-specific expression of myr-BFP). We generated transgene-adaptor strains {*AD1*,*prab-3::myr-mScarlet*} and {*pdes-2::myr-BFP*,*AD2*} and crossed them to generate trans-heterozygote F1 {*AD1*,*prab-3::myr-mScarlet*}/{*pdes-2::myr-BFP*,*AD2*}. Following Cas9/gRNA injection and marker-based screening, as described for the *2×*{*TG*} experiment, we isolated recombinant F2 {*prab-3::myr-mScarlet*, *pdes-2::myr-BFP*} carrying both transgenes. As expected, these recombinants exhibited pan-neuronal myr-mScarlet expression and PVD-specific myr-BFP expression (**Figure 3J**). This demonstrates SuRe’s utility in generating complex single-copy transgenes with controlled expression in a single locus.

### Near optimal recombination efficiency and fidelity with serine recombinases in SuRe

As discussed above, undesired recombination products can arise in the SuRe system due to unwanted homology pairings, particularly when integrating multiple transgenes. To enhance recombination fidelity, we developed SuRe-CR, a refined version of the SuRe system. To distinguish it from the prior version, we termed the original system SuRe-CC. Both SuRe-CC and SuRe-CR use CRISPR/Cas9 for adaptor insertion (denoted by the shared “C”), but they employ different recombination mechanisms. SuRe-CC relies on CRISPR/Cas9-mediated homology-directed repair; SuRe-CR uses serine-recombinase-mediated site-specific recombination (indicated by the “R”). In SuRe-CR, we replaced the gRNA target sites and homologous sequences in the adaptor cassette with *attP* or *attB* sites (**Figure 4A**). Serine recombinases catalyze unidirectional recombination between *attP* and *attB* sites, generating *attL* and *attR* sites^35–37^. Consequently, only an *attR* site remains in the IGA following recombination (**Figure 4A**). This *attR* site does not interact with *attP* or *attB* sites in newly added adaptors, effectively preventing the fidelity loss observed with SuRe-CC. To prevent crosstalk between the enhancers of transgenes *A* and *B*, *AD1* includes an insulator element. After recombination, this insulator is positioned between the two transgenes (**Figure 4A**).

**Figure 4.**
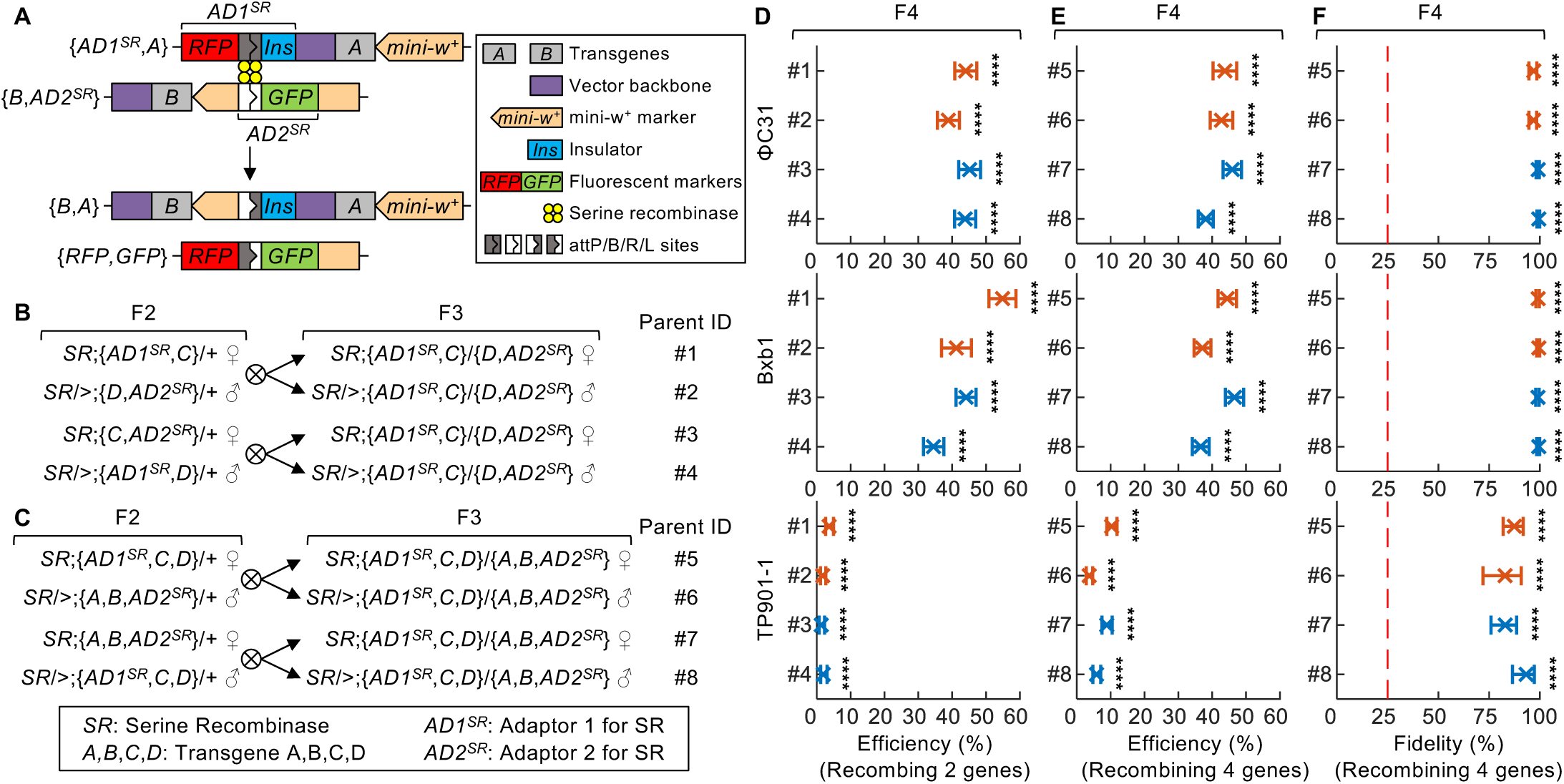
SuRe-CR recombines transgenes at the same locus with high efficiency and fidelity. **(A)** Molecular mechanism of recombinase-mediated recombination in the SuRe-CR system. Like SuRe-CC, SuRe-CR uses CRISPR/Cas9 to insert adaptors (*AD1* and *AD2*) upstream or downstream of the transgene (the adaptor insertion step is omitted from this illustration). However, instead of using CRISPR/Cas9 to induce recombination, SuRe-CR employs a serine recombinase. The adaptors *AD1^SR^* and *AD2^SR^* in SuRe-CR contain reciprocal *attP* and *attB* sites, which are recognized by the serine recombinase. Upon recombination, these sites are converted to *attR* and *attL* sites. The *attR* site remains between the recombined transgenes, while the *attL* site is lost in the byproduct {*RFP*,*GFP*}. Since the recombination process generates a byproduct, the theoretical maximum efficiency of the SuRe-CR system is 50%. To prevent crosstalk between the enhancers of transgenes A and B, an insulator carried by *AD1* is positioned between the transgenes. This working mechanism for SuRe-CR with serine recombinase applies to the experimental details and results presented in Figures 4 and **S4B,E–I**. Colored rectangles and pentagons represent distinct genetic elements. Yellow discs denote the serine recombinase protein. The four different types of rectangles with internal white and/or dark gray patterns respectively denote attP, attB, attR, and attL sites. **(B)** Cross schemes for recombining two transgenes *C* and *D* to create the IGA {*C*,*D*}. The adaptor insertion step is omitted from the illustration. To account for potential maternal effects on recombinase expression, reciprocal crosses between the F2 flies carrying the transgene-adaptor were performed to generate F3 transheterozygote {*AD1^SR^*,*C*}/{*D*,*AD2^SR^*}, which were then crossed with the recombinase strain for F4. Transgenes were the same as in Figure 2: *C*: *13×LexAop2-mCD8::GFP*, and *D*: *R57C10-LexA*. **(C)** Cross schemes for recombining two IGAs {*A*,*B*} and {*C*,*D*} to create an IGA of four transgenes {*A*,*B*,*C*,*D*}. {*A*,*B*} and {*C*,*D*} were created using SuRe-CR as illustrated in **(A)**. Adaptor insertion into {*A*,*B*} or {*C*,*D*} is omitted from the illustration. To account for potential maternal effects on recombinase expression, reciprocal crosses between the F2 flies carrying the IGA-adaptor {*AD1^SR^*,*C*,*D*} and {*A*,*B*,*AD2^SR^*} were performed to generate F3 transheterozygote {*AD1^SR^*,*C*,*D*}/{*A*,*B*,*AD2^SR^*}, which was then crossed with the recombinase strain for F4. Transgenes were the same as in Figure 2: *A*: *10×UAS-IVS-myr::tdTomato*, *B*: *R57C10-GAL4*, *C*: *13×LexAop2-mCD8::GFP*, and *D*: *R57C10-LexA*. **(D–F)** Mean ± 95% C.I. weighted average recombination efficiency and fidelity values (defined as in Figure 2C**,D****,F**) for SuRe-CR systems using the cross scheme in **(B)** and **(C)** (n = 8–19 replicates for each cross, each with a parent of the indicated F3 genotype crossed to the corresponding serine recombinase strain). We assayed three SuRe-CR system variants, each using a different serine recombinase: ΦC31, Bxb1, or TP901-1. The y-axis index corresponds to the F3 parent from the cross designs in **(B)** or **(C)**. The average recombination efficiencies for recombining transgenes *C* and *D* to generate the {*C*,*D*} array were: ΦC31, ∼39–45%; Bxb1, ∼34–55%; TP901-1, ∼1–4%. Average recombination efficiencies for recombining two two-transgene arrays {*A*,*B*} and {*C*,*D*} to create the four-transgene array {*A*,*B*,*C*,*D*} were: ΦC31, ∼38–46%; Bxb1, ∼37–47%; TP901-1, ∼4–10%. All these efficiencies were significantly higher than that of the natural recombination (binomial test; ****: *P* < 10^-6^). Notably, the efficiencies of the ΦC31 and Bxb1 versions of SuRe-CR system were close to the theoretical maximum (50%). Mean fidelity values for recombining two-transgene arrays {*A*,*B*} and {*C*,*D*} to create the four-transgene array {*A*,*B*,*C*,*D*} were all above 83% or even close to 100%: ΦC31, ∼97–99%; Bxb1, ∼99%; TP901-1, ∼83–93%, significantly higher than the theoretical fidelity of 25% for the SuRe-CC system (red dashed line, binomial test; ****: *P* < 10^-6^; n = 8–19 replicates per cross; see Figure 2E for further details). This indicates a dramatic improvement in accuracy achieved by the SuRe-CR system as compared to SuRe-CC.

We chose serine recombinases for SuRe-CR due to their unidirectional recombination mechanism, which enhances fidelity and simplifies the system compared to tyrosine recombinases (**Figures 4A** and **S4A–F**). Tyrosine recombinases catalyze a reversible reaction where the original recombination sites are retained in the product. For example, Cre catalyzes recombination between two *loxP* sites, but the resulting product retains *loxP* sites^38^ (**Figure S4A**). This can lead to unintended recombination and instability, particularly as *loxP* sites accumulate over multiple recombination cycles (**Figure S4C**). Although mutated *loxP* variants can mitigate these issues, it requires *N*–1 distinct mutated *loxP* sites for recombining *N* genes into the desired IGA product (**Figure S4D**), increasing complexity and limiting the reusability of the SuRe Recombinator strains. By contrast, serine recombinases mediate an unidirectional reaction between *attP* and *attB* sites^35–37^, leaving only *attR* sites in the IGA product (**Figures 4A** and **S4B**). These *attR* sites do not interact with *attP* or *attB* sites in subsequently added adaptors, for greater fidelity and simplicity with (**Figure S4E,F**).

We developed three versions of SuRe-CR, each using a different serine recombinase (ΦC31^35^, Bxb1^36^ and TP901-1^37^) and a specific *attP* and *attB* sequence^35–37^. We evaluated the efficiency and fidelity of all three SuRe-CR versions in recombining two or four transgenes, and all showed significantly higher recombination efficiency than natural recombination (**Figure 4B–F**). When combining two transgenes, *A* and *B*, the desired recombination between transgene-adaptor strains {*AD1*,*A*} and {*B*,*AD2*} generates the desired {*B*,*A*} array accompanying by an undesired {*RFP*,*GFP*} array (**Figure 4A**). Consequently, even if all {*AD1*,*A*} and {*B*,*AD2*} undergo the desired recombination, only half of the resulting progeny will possess the desired {*B*,*A*} configuration. This establishes a theoretical maximum recombination efficiency of 50% for the SuRe-CR system.

We found that both ΦC31 and Bxb1 versions of SuRe-CR achieved very high or even near-theoretical maximum recombination efficiency for generating both two-transgene and four-transgene assemblies (**Figure 4D,E**). Specifically, the ΦC31 version reaches ∼39–45% when recombining two transgenes and ∼38–46% efficiency when combining two-transgene arrays to generate a four-transgene assembly. Similarly, the Bxb1 version achieved ∼34–55% efficiency for two-transgene recombination and ∼37–47% for four-transgene assembly. The TP901-1 version had lower efficiency (∼1–4% for two-transgene and ∼4–10% for four-transgene), but is still sufficient for practical use, especially with the aid of adaptor fluorescent markers to facilitate screening. Importantly, all three versions of SuRe-CR maintained very high fidelity, with ΦC31 and Bxb1 approaching 100% and TP901-1 reaching ∼90% (**Figure 4F**). This further validates the advantages of serine recombinases for precise and reliable transgene recombination.

The availability of multiple SuRe-CR versions, each using a different serine recombinase and corresponding recombination sites, enhances the system’s versatility and compatibility with diverse genetic elements. For example, some transgenes, such as those in SPARC^39^ and intMEMOIR^40^ systems, have pre-existing *attP* or *attB* sites recognizable by specific serine recombinases. The intMEMOIR tool, for instance, utilizes *attP* and *attB* sites flanking Bxb1-inducible cassettes^40^. In such cases, the Bxb1 version of SuRe-CR would disrupt these cassettes. However, the ΦC31 version, which relies on a different recombinase and recognition sites, can be safely employed, highlighting the importance of having multiple recombinase options within the SuRe-CR toolkit.

### Generating recombinant products on the mega-base scale

We define the capacity of the SuRe system as the maximum size of a recombinant product, or IGA, that can be successfully generated (**Figure 5A**). As the IGA grows longer with each round of recombination, the increasing distance between the two adaptors may hinder their proximity during subsequent recombination events, potentially reducing efficiency. Eventually, the efficiency becomes too low to generate the desired products. To assess SuR’s capacity, we designed an experiment to simulate recombination of large DNA segments at a single locus (**Figure 5B**). We recombined a transgene carrying *AD1* ({*TG1*,*AD1*}) in the *attP40* site (*25C6* locus) with a series of transgenes carrying *AD2* ({*AD2*,*TG2*}) located at progressively distant genomic loci (**Methods**). By treating the intervening genomic region between the *TG1* and *TG2* locations as a single large segment, successful recombination would result in a duplication of this region. The maximum size of the observed duplication product thus establishes a lower bound for the SuRe’s capacity (**Figure 5B**).

**Figure 5.**
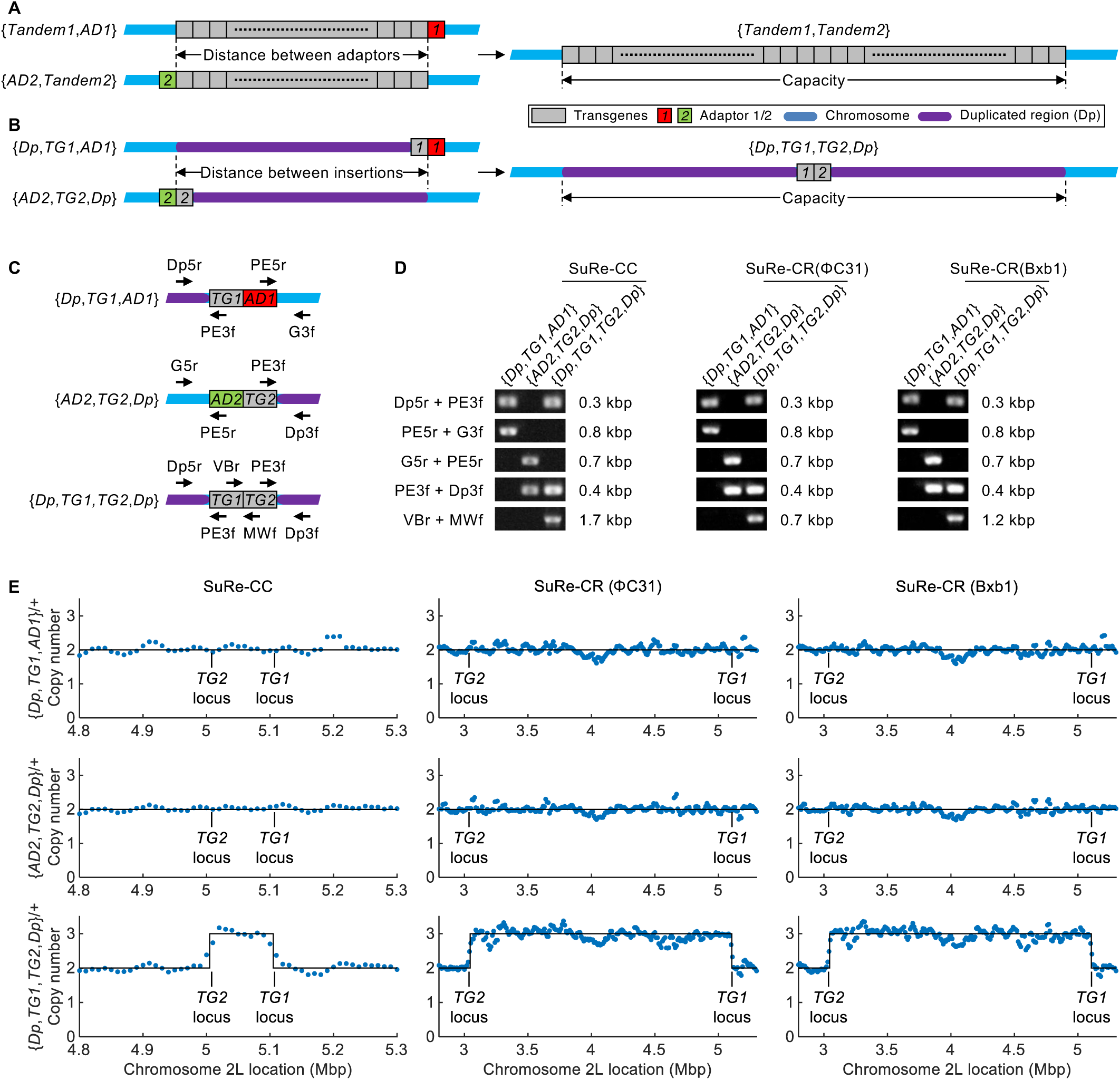
SuRe-CR can generate recombinant products on the mega-base scale. **(A)** Schematic showing the assembly of IGAs with a large number of individual transgenes using SuRe. The system’s capacity is defined as the maximum size of a combined DNA segment that can be successfully generated. For a library of transgenes located in the same genomic locus, as more transgenes are combined with the SuRe system, the distance between the adaptor pair *AD1* and *AD2* increases, making recombination progressively less efficient. Eventually, the efficiency becomes too low for practical use, indicating that the combined transgene segment has reached the limit of the SuRe system’s capacity. Gray, red and green rectangles respectively represent the transgenes, *AD1,* and *AD2*. Blue and purple bars respectively represent the chromosome and the duplicated region. The legend also applies to panels **(B)** and **(C)**. **(B)** Schematic of a strategy to assay the SuRe system’s capacity via duplication of a large DNA segment. Transgenes *TG1* and *TG2* are located at different genomic loci. We inserted adaptor *AD1* upstream of *TG1* and adaptor *AD2* downstream of *TG2*. The DNA segment between the two transgene loci can be viewed as a large IGA (similar to the one in **A**) to be further recombined. Recombining *TG1* and *TG2* creates a duplication of the DNA segment (*Dp*). By systematically increasing the distance between transgenes *TG1* and *TG2* and assaying whether they can be recombined to form a duplication of the DNA segment between them, we can evaluate SuRe’s maximum capacity for constructing recombinant products. **(C,D)** Primer design, **(C)**, and results, **(D**), for PCR genotyping of transgenic strains carrying an adaptor, {*Dp*,*TG1*,*AD1*} and {*AD2*,*TG2*,*Dp*}, as well as the duplication strains {*Dp*,*TG1*,*TG2*,*Dp*}. Specific primers were designed to target the regions flanking the edges of the original transgene *TG1* or *TG2*, the adaptor insertion sites, and the adaptor residual. The *TG1* was *R57C10-LexA* in *attP40* (25C6) locus. The *TG2* recombined by SuRe-CC is *P{lacW}Tfb5^k10127^*, which is 0.1 Mbp away from *attP40*. The *TG2* recombined by SuRe-CR (both ΦC31 and Bxb1 versions) was *P{lacW}FASN2^k05816^*, 2.1 Mbp away from *attP40*. PCR results confirmed the expected amplicons at the edges of {*Dp*,*TG1*,*AD1*} and {*AD2*,*TG2*,*Dp*}, and also at the upstream edge of *TG1*, the downstream edge of *TG2*, and the adaptor residual in {*Dp*,*TG1*,*TG2*,*Dp*}. **(E)** Whole genome sequencing analysis of copy number variances (CNVs) of the duplication region in indicated strains genotyped in **(D)**. CNVs are shown as the normalized sequencing read densities (blue dots) at the corresponding genomic coordinates along the *x*-axis. Black solid lines represent the estimated copy number inferred by the analysis. In heterozygous transgene-adaptor strains {*Dp*,*TG1*,*AD1*}/+ and {*AD2*,*TG2*,*Dp*}/+, the CNVs within the DNA segment between *TG1* and *TG2* are the same as those in its surrounding DNA. However, the CNVs in the DNA segment increased to 3 in the heterozygous duplication strain {*Dp*,*TG1*,*TG2*,*Dp*}/+, confirming successful duplication of the targeted DNA segment between *TG1* and *TG2*. Based on these results, the capacity of the SuRe-CC system was determined to be at least 0.2 Mbp (2 × 0.1 Mbp), while the SuRe-CR(ΦC31) and SuRe-CR(Bxb1) systems exhibited a capacity in creating a recombinant product exceeding 4.2 Mbp (2 × 2.1 Mbp). *TG1* and *TG2s* are the same as those in **(D)**.

To validate the successful generation of duplications, we designed five primer pairs for PCR verification (**Figure 5C**). Four pairs targeted the regions flanking the edges of the transgene-adaptors (two for {*TG1*,*AD1*} and two for {*AD2*,*TG2*}), while one pair targeted the adaptor residual sequence of the desired recombination product (**Figure 5C**). PCR results confirmed successful duplication by both SuRe-CC and SuRe-CR (**Figure 5C,D**). SuRe-CC generated duplication products up to 0.2 Mbp in size (duplicating 0.1 Mb segment), whereas SuRe-CR (using either ΦC31 or Bxb1) produced duplication products >4.2 Mbp (duplicating 2.1 Mb segment). Further, we performed whole-genome sequencing and copy number variation (CNV) analysis to verify the accuracy of the duplications (**Figure 5E**). CNV data revealed a copy number increase of 1 in the duplicated region compared to other genomic regions, validating successful generation of the desired duplications (**Figures 5E** and **S6**).

This experiment showcases the SuRe system’s potential for constructing large IGAs and generating large genome duplications. SuRe can also generate large deletions based on the relative position of *AD1* and *AD2* (**Figure S5A,B**). The duplication and deletion capabilities further expand the applicability of SuRe in genetic engineering and synthetic biology.

### Polytransgenic animals engineered with SuRe for imaging of neural activity

Creating transgenic animals with multiple reporter and effector genes is crucial for many complex imaging and manipulation experiments. Traditional recombination approaches are often inefficient and time-consuming when assembling multiple transgenes. In contrast, the SuRe system offers an efficient solution for generating sophisticated animals harboring a large number of transgenes.

To show its utility for advanced neural imaging, we used SuRe to create a tandem duplication of the Ace::mNeon voltage indicator^41^ (*2×*{*Ace::mNeon*}) by recombining two individual *Ace::mNeon* transgenes. We confirmed the transgene-adaptor strains {*AD1*,*Ace::mNeon*} and {*Ace::mNeon*,*AD2*}, as well as the *2×*{*Ace::mNeon*} duplication product with PCR genotyping (**Figure 6A**). When expressed in dopamine neurons (DANs) via the *R82C10-LexA* driver, *2×*{*Ace::mNeon*} exhibited approximately doubled fluorescence intensity in PPL1-α’2α2 and PPL1-γ2α’1 DAN subtypes, compared to the single *Ace::mNeon* transgene (**Figure 6B**). Quantification of this improvement revealed a significant enhancement in both the mean intensity and signal-to-noise ratio (SNR) for *2×*{*Ace::mNeon*} in both DAN populations (**Figure 6C**). Here, the SNR in voltage imaging is defined as the ratio of the average amplitude of the fluorescence signal during neural spikes to the s.d. of the fluorescence intensity during the baseline (non-spiking) period. This example highlights SuRe’s potential as a generalizable approach for enhancing gene expression and improving functionality by increasing the copy number of a desired gene.

**Figure 6.**
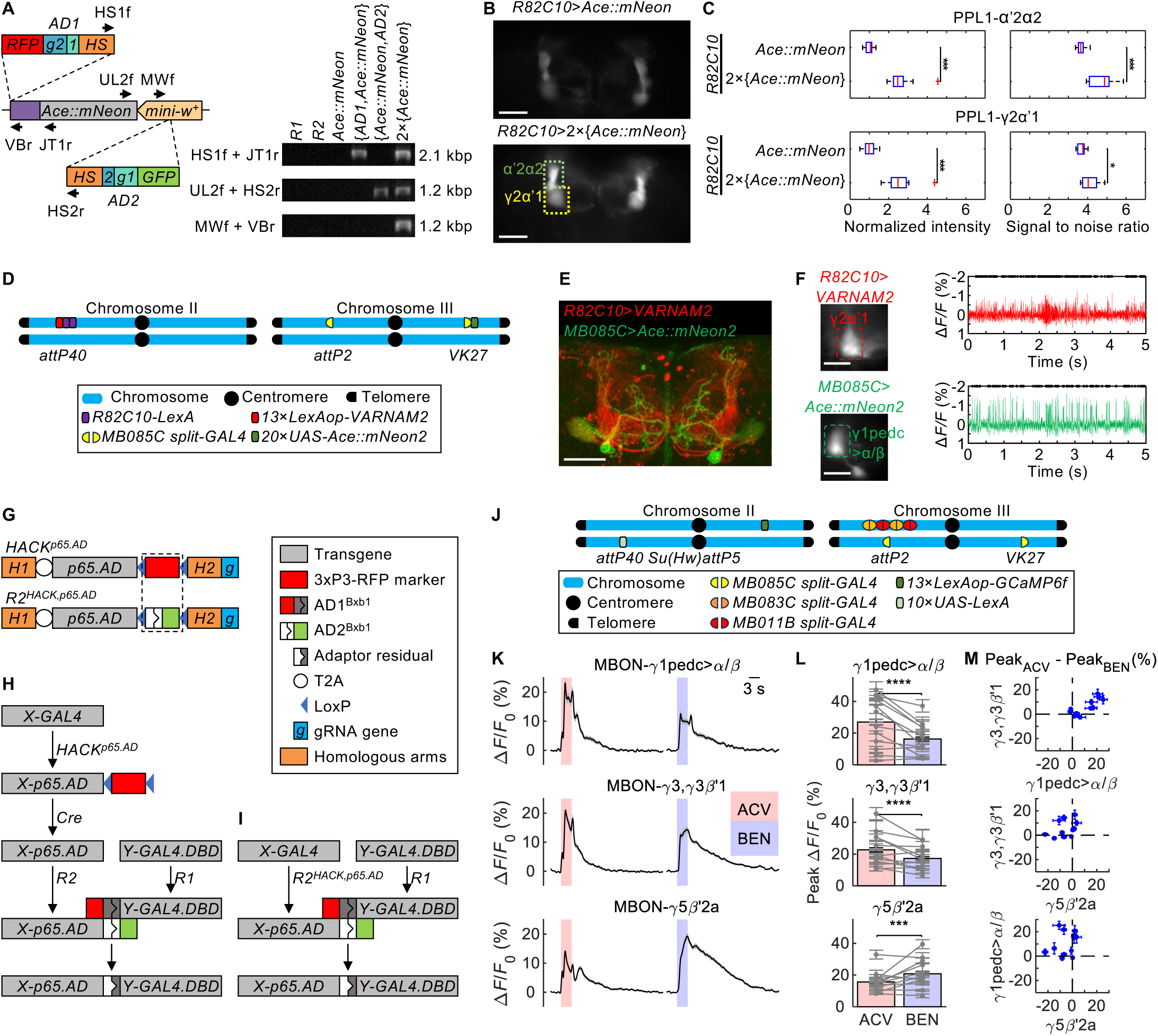
Engineering polytransgenic flies for targeted imaging of neuronal voltage activity. **(A)** PCR genotyping with indicated primers to confirm the generation of a dual-copy version of the *Ace::mNeon* fluorescent voltage indicator using SuRe-CC. *R1* and *R2* are the Recombinator strains of the SuRe-CC system. *Ace::mNeon* is *26×LexAop-Ace::mNeon*. {*AD1*,*Ace::mNeon*} is the strain with the adaptor *AD1* inserted to the upstream of *Ace::mNeon*, and {*Ace::mNeon*,*AD2*} is the strain with the adaptor *AD2* inserted to the downstream of *Ace::mNeon*. *2×*{*Ace::mNeon*} is the desired product of recombination between {*AD1*,*Ace::mNeon*} and {*Ace::mNeon*,*AD2*}. **(B)** Representative images showing expression of the *Ace::mNeon* (top) and *2×*{*Ace::mNeon*} (bottom) voltage indicators under the control of a LexA driver *R82C10* that labels two dopamine neuron (DAN) subtypes, PPL1-α’2α2 and PPL1-γ2α’1. The *2×*{*Ace::mNeon*} strain had significantly higher fluorescence intensity than the *Ace::mNeon* strain under identical imaging conditions, indicating greater Ace::mNeon expression. Dashed boxes enclose the PPL1-α’2α2 and PPL1-γ2α’1 neurons. Scale bars: 50 μm. **(C)** Quantitative comparisons of the mean intensity and signal-to-noise ratio (SNR, the ratio of the mean peak fluorescence intensity during neural action potentials to the s.d. of the baseline fluorescence intensity) of *Ace::mNeon* and *2×*{*Ace::mNeon*} in PPL1-α’2α2 and PPL1-γ2α’1. There were significant increases in both average intensity and SNR for *2×*{*Ace::mNeon*} compared to *Ace::mNeon* in both neuron-types (Wilcoxon rank-sum test; *: *P* < 0.05, ***: *P* < 0.001), highlighting the improved performance of the IGA for voltage imaging. Multiple flies (n = 12) were used for each genotype, with three imaging sessions per fly. **(D)** Schematic of transgene locations in SuRe-engineered multi-transgene strain for advanced voltage imaging. The strain carries six transgenes: a copy of the red fluorescent voltage indicator *13×LexAop-VARNAM2* and two copies of *R82C10-LexA driver* integrated in the *attP40* site by SuRe-CC, a copy of *R52H01-p65.AD* (the *AD* hemidriver of *MB085C split-GAL4*) and a copy of *20×UAS-Ace::mNeon2* integrated in *VK27* site by SuRe-CC, and a copy of *R52B07-GAL4.DBD* (the *DBD* hemidriver of *MB085C split-GAL4*) alone in *attP2* site. In this strain, two copies of *R82C10-LexA* drive the expression of the red fluorescent voltage indicator *VARNAM2* in the DANs, and the combined presence of the two split portions of *MB085C-GAL4* drives expression of the green fluorescent voltage indicator Ace::mNeon2 in mushroom body output neurons (MBONs). **(E)** Confocal microscopy image showing expression of VARNAM2 in PPL1-α’2α2 and PPL1-γ2α’1, and expression of Ace::mNeon2 in MBON-γ1pedc>α/β neurons from the strain of **(D)**. The distinct expression patterns of these indicators in their respective neural populations enables concurrent imaging of voltage dynamics in these cells in individual flies. Scale bar: 50 μm. **(F)** Dual-color voltage imaging of neural spiking in PPL1-γ2α’1 and MBON-γ1pedc>α/β neurons using the strain in **(D)**. Scale bars: 50 μm. Red and green traces are representative recordings from PPL1-γ2α′1 and MBON-γ1pedc>α/β using the two indicators. **(G–I)** The SuRe-HACK system integrates SuRe-CR and HACK (homology assisted CRISPR knock-in) systems to allow conversion of a *GAL4* into a *Split-Gal4* hemidriver and then integration with its hemidriver counterpart at the same locus. In existing libraries, the two hemidrivers of a *split-GAL4* are usually at separate genomic locations. Combining specific hemidrivers into a single locus generally helps to simplify genetic crosses and animal husbandry for experimental applications. This can be achieved by either integrating the existing HACK and SuRe-CR systems **(H)** or using the streamlined SuRe-HACK System **(I)**. The legend applies to panels **(G–I).** Gray, red, blue and orange rectangles represent distinct genetic elements. The red rectangle joined with a dark gray rectangle represents the *AD1* of SuRe-CR(Bxb1). The green rectangle joined with a white rectangle represents the *AD2* of SuRe-CR(Bxb1). The rectangle colored white and dark gray denotes the adaptor residual. The white circle denotes the sequence encoding the T2A peptide. The blue triangle denotes the LoxP site. **(G)** Comparison of the designs of the HACK system (*HACK^p^*^65^*^.AD^*, top) and the Recombinator *R2* of the SuRe-HACK system (*R2^HACK.p^*^65^*^.AD^*, bottom). *Cas9* transgenes are omitted. As highlighted by the dash line box, *R2* of the SuRe-HACK system was made by replacing the fluorescent marker *3×P3-DsRed* in the HACK system with the *AD2* cassette from the SuRe-CR(Bxb1) system. **(H)** Schematic of the genetic manipulation process using the existing HACK and the SuRe-CR systems to generate a recombinant with two hemidrivers of a *split-GAL4* locating in a single locus. The *split-GAL4* consists of *X-p65.AD* and *Y-GAL4.DBD*, where *X* and *Y* represent different promoters. Usually, an available *X-p65.AD* hemidriver is not at the same location as the *Y-GAL4.DBD* hemidriver, but an *X-GAL4* is. Combining the HACK and the SuRe-CR systems can generate a recombinant with *X-p65.AD* and *Y-GAL4.DBD* hemidrivers at a single locus in 4 steps: 1) converting the *X-GAL4* into *X-p65.AD* by *HACK^p^*^65^*^.AD^*; 2) removing the fluorescent marker *3×P3-DsRed* in *X-p65.AD* by recombinase Cre; 3) inserting *AD2* into *X-p65.AD* and inserting *AD1* into *Y-GAL4.DBD* by *R1* and *R2* of the SuRe-CR system; 4) rrecombining {*X-p65.AD*,*AD2*} and {*Y-GAL4.DBD*,*AD1*} to generate the desired recombinant {*X-p65.AD*,*Y-GAL4.DBD*}. **(I)** Schematic representation of the streamlined genetic manipulation process using the SuRe-HACK system to generate a recombinant with two hemidrivers of a *split-GAL4* (*X-p65.AD* and *Y-GAL4.DBD*). The recombinator strain *R2^HACK.p^*^65^*^.AD^* simultaneously converts a *GAL4* transgene into *p65.AD* and inserts the *AD2* adaptor, simplifying the first three steps in **(H)** into one. The *AD1* insertion in *Y-GAL4.DBD* and the rest of the recombination step are the same as those in **(H)**. **(J)** Application of the SuRe-HACK system to create a transgenic animal for simultaneous Ca^2+^ imaging of multiple neuron types. The strain has two copies of *MB083C split-GAL4* (each with two hemidrivers) and two copies of *MB011B split-GAL4* (each with two hemidrivers). These eight transgenic components were integrated at the *attP2* site using the SuRe-HACK and SuRe-CR systems. The strain also contains a set of *MB085C split-GAL4* (one hemidriver in *attP2* and the other in *VK27* site), a single copy of the *10×UAS-LexA* and a single copy of the Ca^2+^ indicator *13×LexAop-GCaMP6f*. *Split-GAL4s* drive *10×UAS-LexA* expression, which yields *13×LexAop-GCaMP6f* expression in three distinct MBONs. **(K)** Traces of the relative (*ΔF*/*F*) changes in fluorescence obtained during concurrent imaging of Ca^2+^ activity in the MBON-γ1pedc>α/β, MBON-γ3,γ3β’1, and MBON-γ5β’2a neurons in response to apple cider vinegar (ACV) and 3% benzaldehyde (BEN), using the strain in **(J)**. Black curves: mean activity traces. Gray shaded regions: S.E.M. (n = 12 flies). Color shaded regions: time intervals (3 s) during which the indicated chemicals ACV or BEN were presented. These traces reveal the distinct response patterns of different MBON populations to specific odor stimuli. Twelve flies (n = 12) were used, with three imaging sessions per fly. In each experiment, ACV and BEN were presented three times. **(L)** Quantitative analysis of the responses of different MBON populations to ACV and BEN. MBON-γ1pedc>α/β, MBON-γ3,γ3β’1 neurons had significantly higher responses to ACV than to BEN; MBON-γ5β’2a neurons showed the opposite pattern, responding more strongly to BEN (Wilcoxon signed-rank test; ***: *P* < 0.001, ****: *P* < 10^-6^). Gray data points and error bars: mean ± S.E.M. of the nine technical replicates (3 imaging sessions, 3 odor presentation per session) from each fly. **(M)** Joint distributions of the difference in the responses to the ACV and BEN odors across pairs of MBONs. Blue data points and error bars: mean ± S.E.M. of the nine technical replicates (3 imaging sessions, 3 odor presentation per session) from each fly.

To investigate how activity is coordinated across various neural types, it is often desirable to use multiple cell-specific drivers to express distinct indicators in the different cell types. However, traditional methods for assembling transgenic animals typically limit neural imaging to a single driver per individual, hindering this type of research.

Building on our lab’s prior development of multicolor fluorescent voltage indicators with diverse response kinetics^42^, we used SuRe to engineer a fly strain for dual-color voltage imaging. This strain carries a total of 6 transgenes: a copy of the red fluorescent voltage indicator *13×LexAop-VARNAM2* and two copies of *R82C10-LexA* driver integrated in the locus *attP40*, a copy of *R52H01-p65.AD* (the activation domain hemidriver of *MB085C split-GAL4*) and a copy of *20×UAS-Ace::mNeon2* integrated in the locus *VK27*, and a copy of *R52B07-GAL4.DBD* (the DNA binding domain hemidriver of *MB085C split-GAL4*) alone in the locus *attP2* (**Figure 6D**). In this strain, *R82C10-LexA* drives expression of the red fluorescent voltage indicator, VARNAM2, in DANs; we used two copies of *R82C10-LexA* to enhance the expression level. The joint presence of the two hemidrivers of *MB085C-GAL4* drives expression of the green fluorescent Ace::mNeon2^42^, an molecular optimized version of the Ace::mNeon voltage indicator, in mushroom body output neurons (MBONs) (**Figure 6E,F**). With this expression strategy, we recorded neural spiking from DANs and MBONs (**Figure 6F**), demonstrating SuRe’s ability to generate sophisticated animals for multi-color neural imaging studies.

### SuRe-HACK system for combining split drivers in the same locus

Binary systems like GAL4/UAS^43,44^, LexA/LexAop^45^, and QF/QUAS^46,47^ are potent tools for cell-type-specific control of gene expression in *Drosophila* and other organisms. Split versions of these systems, including split-GAL4^29,48^, split*-*LexA^49^, and split-QF^49,50^, offer even greater precision (as illustrated above in the prior section). By requiring co-expression of activation (AD) and DNA binding domain (DBD) hemidrivers for activity, split systems restrict gene expression to the intersection of two cell populations for highly specific genetic targeting^29,48–50^.

In the Janelia *split-GAL4* collection, the *AD* and *DBD* hemidrivers are often located at distal loci to enable easy combination by traditional genetic approaches^51^. However, this arrangement can complicate further genetic manipulations and strain maintenance, especially when integrating additional components or preventing hemidriver separation during crosses. Integrating both hemidrivers at the same locus avoids these complications. To achieve this, we combined SuRe with the Homology-Assisted CRISPR Knock-in (HACK) system^30^ (**Figure 6G–I**). HACK allows *in vivo* conversion of existing *GAL4* drivers into *QF2*, *GAL80*, or *split-GAL4*^30,52^. As an example, we focused on converting a full-length *GAL4* driver to the *AD* hemidriver.

In the Janelia *split-GAL4* collection, a *X-p65.AD* hemidriver (an *AD* hemidriver where *X* represents a promoter) is located in *attP40* or *VK27*; its corresponding *Y-GAL4.DBD* hemidriver (a *DBD* hemidriver where *Y* represents a promoter different from *X*) is located in an *attP2* site^51^. Janelia *GAL4* insertions are in the *attP2* site^25,29^, the same locus as *Y-GAL4.DBD*. We used HACK to convert an *X*-*GAL4* into the corresponding *X-p65.AD* hemidriver, allowing subsequent integration of both hemidrivers at a single locus with SuRe.

This conversion and integration can be done by combining the existing HACK and SuRe-CR systems in four steps (**Figure 6H**): (1) converting the *X*-*GAL4* (in *attP2*) into the *X-p65.AD* hemidriver using *HACK^p^*^65^*^.AD^*, with the resulting hemidriver located in the same locus as its *Y-GAL4.DBD* counterpart in *attP2*; (2) using Cre recombinase to remove the residual fluorescent marker from the converted *X-p65.AD* hemidriver; (3) inserting *AD2* into the *X-p65.AD* hemidriver and *AD1* into the *Y-GAL4.DBD* hemidriver using SuRe-CR; (4) recombining the two adaptor-hemidrivers to generate the combined *split-GAL4* driver {*X-p65.AD*,*Y-GAL4.DBD*}.

To streamline this process, we developed SuRe-HACK, which integrates HACK and SuRe-CR into a single system. This uses a HACK-compatible *R2* strain (*R2^HACK.p^*^65^*^.AD^*) that replaces the HACK system’s fluorescent marker with the *AD2* adaptor cassette from SuRe-CR(Bxb1) (**Figure 6G**). This change allows *R2^HACK.p^*^65^*^.AD^* to both convert a *GAL4* transgene into a *p65.AD* hemidriver and insert the *AD2* adaptor, simplifying the workflow (**Figure 6I**). The *AD1* insertion and subsequent recombination steps remain the same as in the standard SuRe-CR system.

This advance greatly enhances the efficiency of generating animals harboring components for multiple binary or/and split-binary systems, allowing previously infeasible genetic configurations. To showcase this versatility, we used SuRe-HACK and SuRe-CR to create a fly line expressing the Ca^2+^ indicator GCaMP6f^53^ in three distinct MBON populations via their respective *split-GAL4* drivers (*MB011B*, *MB083C*, *MB085C*). This fly strain comprises 12 transgenes across three loci (**Figure 6J**).

Specifically, the strain has two copies of *MB083C split-GAL4* (each with two hemidrivers, where the *p65.AD* hemidriver with *AD2* was made by SuRe-HACK) and two copies of *MB011B split-GAL4* (each with two hemidrivers, where the *p65.AD* hemidriver with *AD2* was made by SuRe-HACK), totaling eight hemidrivers integrated to the *attP2* site using SuRe-CR (**Figure 6J**). The strain also has a set of *MB085C split-GAL4* (one hemidriver in *attP2*, the other in *VK27* site), one copy of the *10×UAS-LexA* and one copy of the Ca^2+^ indicator (*13×LexAop-GCaMP6f* ; **Figure 6J**). In this configuration, *split-GAL4* drivers activate *10×UAS-LexA* expression, which in turn drives *13×LexAop-GCaMP6f* expression in the three distinct MBON populations. We chose *10×UAS-LexA* and *13×LexAop-GCaMP6f* to amplify GCaMP6f expression, which allowed us to perform concurrent imaging of baseline and odor-evoked Ca^2+^ activity in these MBONs (**Figure 6K–M**).

Our results show that MBON-γ1pedc>α/β and MBON-γ3,γ3β’1 neurons exhibit responses to apple cider vinegar (ACV) of greater amplitude than those to 3% benzaldehyde (BEN). MBON-γ5β’2a neurons displayed the opposite pattern and responded more strongly to BEN (**Figure 6L**). These observations fit with prior reports showing that MBONs innervating the γ1 and γ3 mushroom body compartments encode the innate valences of attractive odor, whereas those innervating the γ5 compartment encode the innate valence of repulsive odors^54–57^. Our optical study reveals multineuronal responses and the cells’ covariations in their responses to ACV and BEN (**Figure 6M**), which cannot be measured with recordings of only one MBON at a time. Further multineuronal imaging studies of the type shown here seem poised to provide valuable insights into the different neurons’ distinctive odor sensitivities and complementary functional roles in the olfactory system.

### Mathematical modeling of large-scale transgene assembly

Many synthetic biology applications, such as the construction of complex metabolic pathways, require the integration of many transgenes into one organism^58–61^. The construction of synthetic genomes requires even larger-scale assemblies of transgenic components^4–6^. The SuRe system shows substantial potential for simplifying and accelerating large-scale transgene assembly as compared to current available techniques.

Based on the mechanisms of the different SuRe versions and fidelity measurements described above (**Figures 2E, 4F** and **S3I**), we compared mathematically the workloads and time requirements for transgene assembly using SuRe against those incurred with conventional genetic approaches (**Supplemental Appendix**). We determined the time cost as the number of recombination or integration rounds needed to created the desired organism. The workload includes efforts to create the transgenes and those to screen for the correct product during transgene assembly.

Our mathematical analyses evaluated two assembly processes: sequential and binary (**Figure 7A–D**). Sequential assembly involves recombination or integration of genes linearly from upstream to downstream, adding one transgene per step (**Figure 7A,B**). In binary assembly, two sets of transgenes are recombined at each step, doubling transgene number (**Figure 7C,D**). Traditional natural recombination supports both sequential and binary assembly (**Figure 7A,C**), while sequential integration with genome editing supports only sequential assembly (**Figure 7B**). SuRe supports both sequential and binary assembly (**Figure 7B,D**). Although we considered both strategies in our mathematical modeling, binary assembly proved to be substantially more efficient, and we thus used this approach in our experiments.

**Figure 7.**
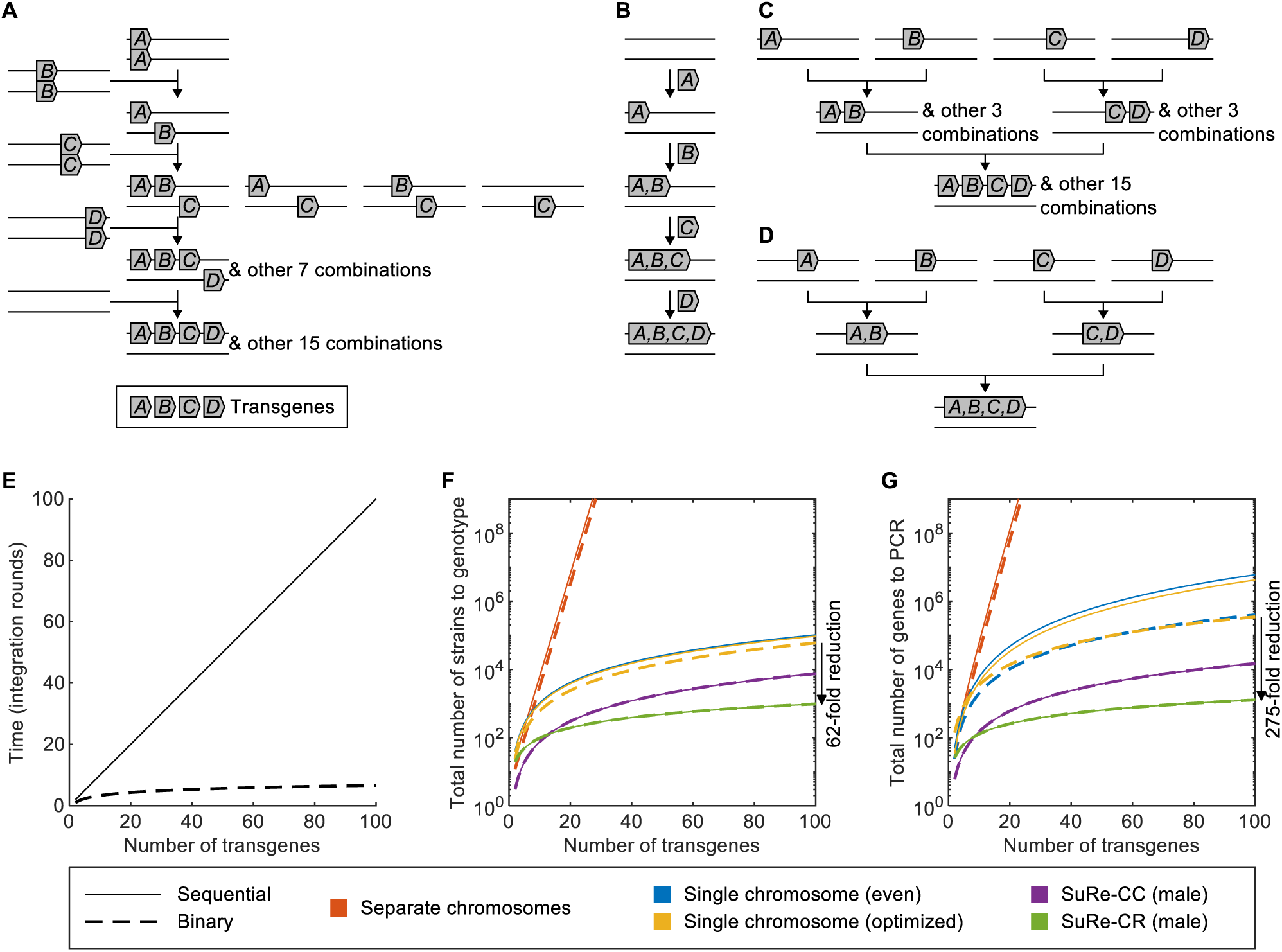
Mathematical modeling predicts the superior efficiency of SuRe for large-scale transgene assembly. **(A)** Illustration of the conventional sequential assembly process via natural recombination, in which transgenes are added sequentially one-by-one via recombination at separate locations. This leads to a rapid increase in undesired recombination products, due to transgene segregation. **(B)** Illustration of the conventional sequential assembly process via genome editing, in which transgenes are added linearly at a single locus. Integrated transgenes do not segregate in subsequent steps. SuRe can also perform sequential integration, preventing transgene segregation. **(C)** Illustration of the conventional binary assembly process via natural recombination, in which transgenes at distinct loci are recombined in parallel, doubling the gene number per step. This leads to a rapid increase in undesired recombination products due to transgene segregation like the conventional sequential assembly process via natural recombination. **(D)** Illustration of the binary assembly process with the SuRe system, in which transgenes at a single locus are recombined in parallel, doubling the gene number per step. Unlike the binary assembly process shown in **(C)**, transgenes recombined by SuRe do not segregate in subsequent steps. Note that the SuRe system can also perform sequential integration. **(E)** The time required for sequential and binary assembly processes. The former rises linearly with the total number of transgenes, whereas the latter rises logarithmically. Note that a natural recombination strategy in principle allows both sequential and binary assembly, but with practical constraints. SuRe efficiently supports both strategies, whereas genome editing is limited to sequential integration. **(F, G)** Total number of strains requiring genotyping, **(F)**, or genes requiring PCR typing, **(G)**, during transgene assembly using sequential or binary recombination with natural recombination and the SuRe system. Strain genotyping can be achieved by genotyping every recombined transgene, or by a single whole-genome sequencing (WGS) test (**Supplemental Appendix** has details). Red curves show results for scenarios in which transgenes are placed on distinct chromosomes and assembled via natural meiotic independent assortment. Blue and yellow curves show results for scenarios in which transgenes are placed at separate loci on a single chromosome and assembled via natural meiotic recombination. Blue curves show results for the case in which the transgenes are evenly placed; yellow curves show results for the case in which transgene placement is optimized to minimize the genotyping workload (**Supplemental Appendix**). Purple and green curves show results for scenarios in which transgenes are placed at a single locus and assembled with SuRe. Both sequential recombination (solid lines) and binary recombination (dashed lines) strategies are modeled. Binary recombination requires equal or fewer genotyping in generating desired products than sequential recombination. The binary recombination using the SuRe system, especially SuRe-CR (green dashed line), is the most efficient. It reduces the number of strains to be genotyped by over 62-fold (black arrow in **(F)**), and the number of genes to be PCR-typed by over 275-fold (black arrow in **(G)**), for recombining 100 transgenes, compared to traditional genetic methods.

As researchers seldom recombine more than 4 transgenes in one chromosome with traditional genetic approaches, it is hard to find published comparisons of sequential and binary recombination. We systematically compared the two assembly processes, in both traditional genetic approaches and with SuRe. Binary assembly greatly outperforms sequential assembly regarding time costs and requires only log_2_*N* rounds to assemble *N* transgenes, whereas sequential assembly requires *N* rounds (**Figure 7E**). Importantly, this reduction in time is not offset by an increase in workload; our analysis showed that binary assembly also results in reduced genetic screening under all conditions (**Figure 7F,G**; **Table 1** in **Supplemental Appendix**).

We systematically estimated the workload for recombining transgenes with traditional natural recombination. If all the transgenes are on separate chromosomes, the screening workload rises exponentially with the number of genes (**Figure 7F,G**; **§3, §4** and **Table 1** in **Supplemental Appendix**). Moreover, since the number of chromosomes is finite, this limits the number of genes to not more than the number of chromosomes. If the transgenes are all located on a single chromosome, the screening workload is highly dependent on their specific locations. A random distribution of genes results in an infinite workload (**§3, §4** and **Table 1** in **Supplemental Appendix**). In the optimal scenario, where we can precisely control the docking sites of these transgenes to minimize the screening workload, this workload can be much lower than that of the case where they are located on separate chromosomes. Nevertheless, the screening workload remains significantly higher than that of the SuRe system (**Figure 7F,G; §5, §6** and **Table 1** in **Supplemental Appendix**). Achieving this optimal configuration (**Figure S7A–C**) necessitates accurate genome editing and careful verification to avoid disrupting endogenous genes, creating additional workloads. Since transgene locations are specifically tailored for a particular trait, these transgenes may not easily be reused for other trait engineering. In contrast, with SuRe, the integration of all transgenes at the same docking site permits their reuse in other IGAs, enhancing modularity and efficiency.

We also estimated the workload for recombining IGAs with SuRe. The number of strains in an adaptor insertion step or a recombination step is inversely proportional to the fidelity of this step. The fidelity of the recombination step using the SuRe-CC system can be estimated based on the number of HS pairs. When there are two HSs in the transgene-adaptors undergoing recombination (**Figures 2A,B** and **S3A–C**), our results suggest that all possible recombination products will appear with equal probability in the progeny of F3 females (**Figure 2D,E**). However, we observed a bias toward the desired product in males, and we roughly estimate the fidelity in males to be about two fold of that in females (**Figure 2D,E**). In general, when recombining two IGAs containing *n_U_* and *n_D_*transgenes, respectively, the recombination fidelity is estimated to be 1/(*n_U_n_D_*) for females and 2/(*n_U_n_D_*) for males (**§5** in **Supplemental Appendix**). Because the serine recombinase only catalyzes the recombination of the *attP*-*attB* pair, the fidelity of the recombination step using the SuRe-CR system is not related to the number of genes in the IGAs (**Figures 4F** and **S4E,F**).

Both the SuRe-CC and SuRe-CR systems use the CRISPR/Cas9 system to insert the adaptors. It is also possible to create undesired adaptor insertion products (**Figure S3D–G**). If all possible adaptor insertion products occur with equal probability, the fidelity is inversely proportional to the number of homology arms in the transgenic tandem. That is, for an IGA with *n* transgenes, the expected fidelity of adaptor insertion is estimated to be 1/*n* (**Figure S3D–G**). However, unlike the recombination step of SuRe-CC, our empirical data showed that adaptor insertion has a strong bias to create the desired product (**Figures S3I** and **S4H**), and the fidelities in males and females are both about ∼90% (**Figures S3I** and **S4H**; **§6** in **Supplemental Appendix**).

The total workload for screening is the sum of screen workload at each step. We developed a geometric approach to estimate the sum (**Figure S7D,E**; **§5** in **Supplemental Appendix**). Among the different versions of SuRe, the SuRe-CR system coupled with a binary recombination strategy shows the fastest recombination time and least screening workload. This dramatically reduces the screening workload compared to optimal traditional genetic approaches when *e.g.*, recombining 100 transgenes. Specifically, there is a >62-fold reduction in the number of strains and a >275-fold reduction in the number of genes that must be genotyped (**Figure 7F,G**). Further, SuRe only requires a single genome docking site, eliminating the need to test additional sites for transgene insertion.

## Discussion

This paper presents and validates the SuRe system for *in vivo* integration of multiple transgenes at a single genomic locus—which we term the generation of IGAs—in *Drosophila* **(Figures 1, 2, 4** and **5)** and *C. elegans* **(Figure 3)**. We developed and tested two versions of SuRe: SuRe-CC (based on CRISPR/Cas9, **Figures 1–3**) and SuRe-CR (based on CRISPR/Cas9 and site-specific recombination, **Figure 4**). Notably, SuRe-CR exhibited exceptionally high recombination efficiency and fidelity, approaching the theoretical maximum (**Figure 4**), and allowed the generation of megabase-scale recombinant products (up to 4.2 Mb, **Figure 5**). In *Drosophila*, we showed the utility of SuRe for enhancing transgene expression via duplication for advanced neural imaging studies **(Figure 6)**. We successfully used SuRe to create fly strains with 6-12 transgenic components, which had been previously difficult to achieve by other means, for dual- and multi-neuron population imaging experiments (**Figure 6**). Our mathematical modeling showed that SuRe’ capacity for exponential expansion of transgene assemblies, coupled with its low screening workload, offers a significant efficiency advantage over traditional approaches for large-scale transgene assembly (**Figure 7**), thereby paving the way for chromosome-scale gene assembly construction.

### Engineering polytransgenic animals for advanced experiments

The SuRe system’s adaptability extends beyond individual transgenes to entire libraries with similar genetic backbones. For example, the SuRe version of **Figures 1**, **2** and **4** can be applied to the Janelia GAL4/LexA collection^25^, CEP lines^26^, and the Vienna Tile GAL4 (VT-GAL4) library^27,28^, plus the UAS and LexAop lines using vectors from the Janelia Research Campus^29^. This allows researchers to leverage these extensive resources to construct complex transgenic flies with SuRe. By replacing the gRNA and the homologous arms for adaptor insertion, the SuRe system can also be easily adapted to other widely-used transgenic libraries, including the Transgenic RNAi Project (TRiP) collections^62^ and the Vienna Drosophila Resource Center (VDRC) RNAi library^63^.

Further, the hybrid SuRe-HACK system facilitates the integration of split-GAL4 hemidrivers, which are often located at distant genomic loci. By combining the HACK and SuRe-CR systems, one can efficiently relocate a hemidriver and integrate it with its counterpart hemidriver at a single locus, streamlining the generation of complex split-GAL4 drivers. We generated a collection of strains with integrated split-GAL4 hemidrivers and deposited them at the Bloomington Drosophila Stock Center (BDSC) (**Table S4** lists these strains).

SuRe offers a versatile approach to control gene expression levels by precisely duplicating transgenes. In our *Drosophila* studies, increasing the copy number of the Ace::mNeon indicator improved voltage imaging by boosting the fluorescence intensity and signal-to-noise ratio (**Figure 6A–C**). In *C. elegans*, duplication of the *pdes-2::myr-mScarlet::let-858 3’UTR* transgene led to a predictable rise in fluorescence intensity (**Figure 3F–H**), offering a substantial advantage over the unpredictable expression levels often observed with multi-copy extrachromosomal arrays. This ability to fine-tune gene expression via controlled duplication expands SuRe’s set of potential applications to a variety of research areas. Additionally, SuRe is often more efficient than conventional methods, particularly for engineering polytransgenic animals harboring a range of different copy numbers. Its unique ability to exponentially increase copy numbers also makes SuRe especially valuable for projects requiring the generation of a large number of transgene copies.

### Comparisons of SuRe to other means of transgene assembly

Although alternative approaches exist for multi-transgene assembly at the same locus, SuRe offers several advantages. Direct cloning of all transgenes into a single vector is limited by vector capacity. SuRe overcomes this by using multiple vectors, each containing part of the genetic circuit, and *in vivo* recombination, allowing assembly of larger constructs. Further, SuRe permits *in vivo* testing of individual components before assembly, for troubleshooting and replacement of malfunctioning parts.

Knapp *et al.* engineered mutated ΦC31 variants for multi-transgene array generation^64^. The mutated ΦC31 variants also catalyze the reverse reaction from *attL* and *attR* to *attP* and *attB*. It allows insertion of additional transgenes to a transgene made by ΦC31. However, this approach has some drawbacks compared to SuRe. First, the reversibility of mutated ΦC31 integration can lead to unintended gene excisions. Second, it offers limited control over precise transgene landing sites (upstream or downstream of the target site). Third, it lacks fluorescent markers for efficient screening, relying on laborious PCR genotyping. Fourth, it requires sequential gene insertion, which takes *N–*1 steps to insert *N* transgenes.

The SwAP-In (Switching Antibiotic resistance markers Progressively for Integration)^65^ is a method for assembling multiple transgenes and synthesizing genomes in yeast. Building on SwAP-In, the mSwAP-In (mammalian Switching Antibiotic resistance markers Progressively for Integration) system^66^ provides analogous seamless integration of large DNA fragments or multiple transgenes in animals. While SuRe typically leaves adaptor residuals between genes, a new design of variant of SuRe-CC can achieve seamless integration, albeit with a requirement for customized *R1* and *R2* strains for each reaction. mSwAP-In requires sequential integration of each component, requiring *N* steps to integrate *N* transgenes or DNA fragments, which is slower than the SuRe system. Besides, mSwAP-In introduces additional gRNA target sites with each step, increasing the risk of off-target CRISPR/Cas9 activity. SuRe, in contrast, uses a fixed number of gRNAs (4 for SuRe-CC, 2 for SuRe-CR), mitigating this risk.

Overall, the SuRe system constitutes a versatile and efficient platform for multi-transgene assembly, overcoming limitations of existing methods and providing distinct advantages in terms of scalability, fidelity, and ease of use.

### Application of the SuRe system in genome rewriting

The field of synthetic genomics has made remarkable progress, but synthesizing complete genomes for multicellular organisms remains a significant challenge. The largest synthetic genome made to date is that of yeast^67,68^, a unicellular organism, with a genome size of ∼12 Mb^69^. Scaling up genome rewriting technology to multicellular organisms with much larger genomes is a formidable challenge. Unlike yeast, which has a rapid life cycle, most multicellular animals have longer life cycles and more complex sexual reproduction processes. This complexity makes sequential assembly of transgenes inefficient due to fundamental genetic principles, particularly the laws of independent assortment and linkage and crossover, which hinder assembly of many transgenic elements. For example, the genomes of even simple animals like *C. elegans* and fruit flies are ∼100 Mb^70^ and ∼180 Mb^71^ respectively, ten times larger than that of yeast. Mammalian genomes are even larger, about two hundred times the size of yeast^72^. Given that the yeast genome project took nearly two decades^65,68^, directly applying current techniques to animal genomes would require prohibitively long timelines—potentially hundreds of years for simpler animals and thousands of years for mammals.

However, the SuRe system offers a groundbreaking solution by providing exponential acceleration in assembling genetic elements. Synthesizing the entire fruit fly genome (∼180 Mbp) would require the assembly of ∼3,600 transgenic strains, each carrying a 50 kbp synthesized transgene. SuRe can achieve this assembly in just 12 steps, theoretically allowing completion within two years. Moreover, we demonstrated SuRe’s capacity to recombine megabase-scale DNA segments (up to 4.2 Mb, **Figure 5**) in *Drosophila*. This capacity makes SuRe an ideal tool for genome rewriting in sexually reproducing multicellular animals.

### Outlook

Here, our studies with SuRe focused on *Drosophila* and *C. elegans*, but the SuRe-CR system also holds great potential for applications in a wide range of organisms. Both CRISPR/Cas9 and the serine recombinases employed in SuRe-CR (ΦC31 and Bxb1) have been successfully used in various species, including zebrafishes^73,74^, frogs^75,76^, mice^77,78^, human cells^79,80^, and plants^81–84^. This broad applicability suggests SuRe-CR can be readily adapted for efficient and precise transgene manipulation in various model systems. Owing to its capability to generate large-scale genomic duplications or deletions, including as byproduct of the duplication process (**Figure S5A,B**), one potential application of SuRe is to engineer animal models for studies of how pathogenic copy number variants (CNV) influence disease progression.

Beyond the synthesis of and large-scale modifications to the genome, SuRe’s versatility extends to creating novel metabolic pathways, engineering organisms with enhanced stress resistance, or developing new diagnostic tools. Its control over gene expression through copy number duplication unlocks broad possibilities for exploring and manipulating complex biological systems. Overall, we believe SuRe is a transformative tool for synthetic biology and a powerful approach to address challenging biological questions and drive scientific discovery and technological innovation.

### Limitations of the Study

One limitation of SuRe is its reliance on crossing transgenic organisms. Due to this constraint, the total time required for recombination is proportional to the length of the organism’s life cycle, making the system time-consuming in species with long life cycles. Future versions of SuRe that could be applied to embryonic stem cells would likely dramatically accelerate the recombination process.

Residual sequences between transgenic components may limit SuRe’s compatibility. These residuals include homologous arms necessary for adaptor insertion, the homologous sequence (HS) remaining after SuRe-CC application, and the attR site resulting from SuRe-CR recombination. The residual sequences diminish the fidelity of adaptor insertion to levels <100% (**Figures S3** and **S4**). Hence, PCR genotyping is required to ascertain accurate adaptor insertion. Further, in cases where a large gene necessitates division into smaller segments for transgenesis and SuRe-mediated recombination, these residual sequences can disrupt the contiguity of the reassembled gene, effectively partitioning it into discrete, non-functional units. Addressing these limitations requires a redesigned SuRe system that enables seamless recombination, followed by thorough testing.

The impact of genomic context on transgene expression, both in terms of levels and patterns, is well established^64,85^. Integrating transgenes at the same locus may lead to unpredictable expression of individual transgenes due to crosstalk between enhancers. Here, insulators were incorporated to minimize this potential crosstalk. Nevertheless, the efficacy of this approach may be limited in certain contexts. Consequently, future improvement to the SuRe system may necessitate the development of predictive algorithms to identify potential enhancer interference by predicting 3D genome organization and the design of effective strategies to abrogate such interactions.

## Acknowledgments

We gratefully acknowledge an NSF NeuroNex grant to M.J.S. and K. Deisseroth and the Bloomington Drosophila Stock Center (NIH P40OD018537) for providing fly stocks. We thank L. Luo, C. Lyu and J. Chen for helpful discussions.

---

**Figure S1.**
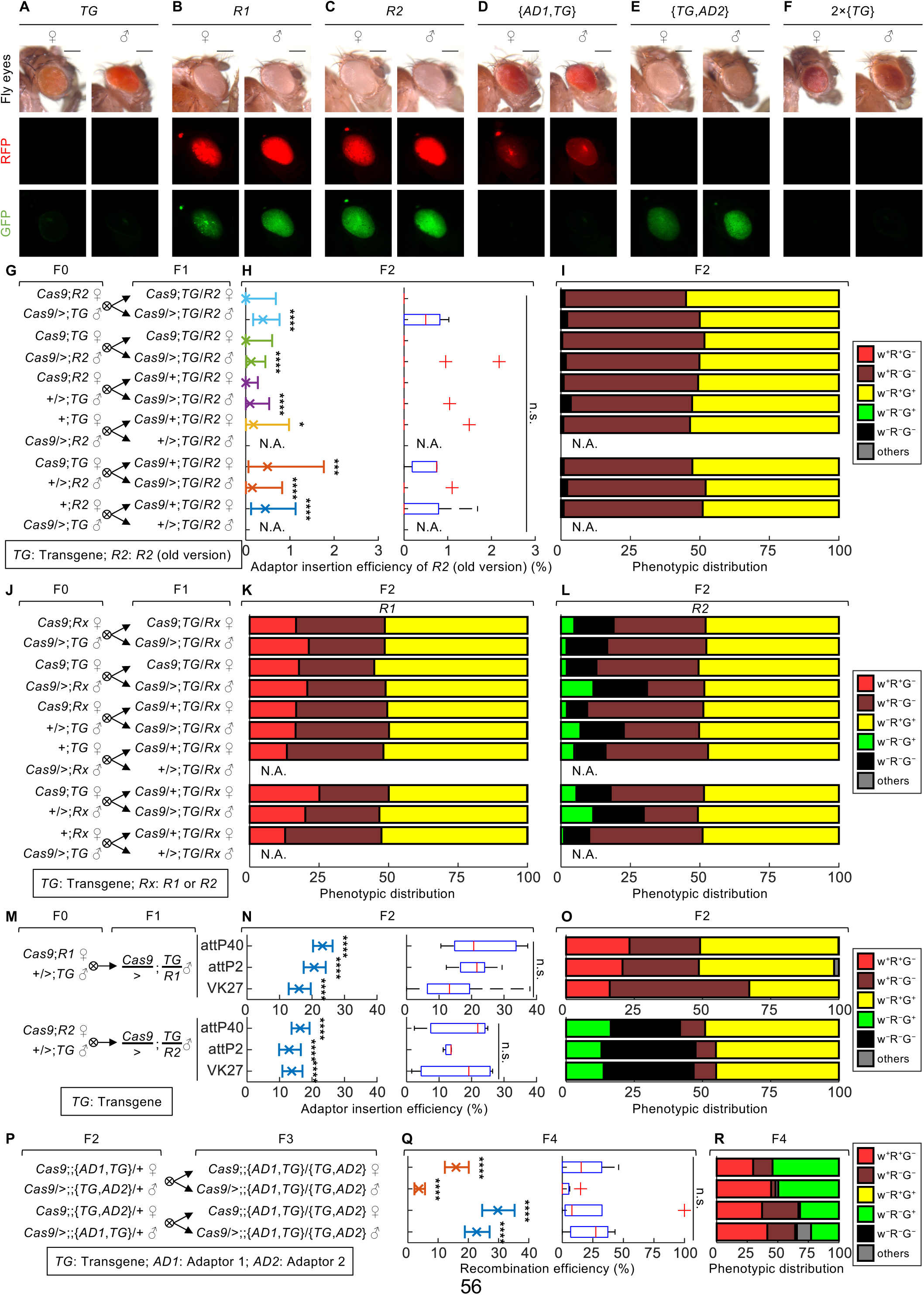
Extended analysis of adaptor insertion and recombination efficiencies in SuRe. **(A–F)** Brightfield (*top row*), red fluorescence (*middle row*) and green fluorescence (*bottom row*) images showing fly eye phenotypes for females and males of the different genotypes used with SuRe. Desired progeny from each cross were selected based on the presence or absence of the eye-specific fluorescent markers, *3×P3-RFP* and *3×P3-GFP*, and the eye pigmentation *mini-w* marker in the transgene vector. **(A)** Strains with a transgene (*TG*) to be recombined show orange-to-red-colored eyes. **(B)** The recombinator strain *R1* contains a *3×P3-GFP* and an *AD1* cassette harboring a *3×P3-RFP*, showing both red and green-fluorescent eyes. **(C)** The recombinator strain *R2* contains a *3×P3-RFP* and an *AD2* cassette harboring a *3×P3*-*GFP*, showing both green and red-fluorescent eyes. **(D)** The {*AD1*,*TG*} strain contains a *TG* and an *AD1* harboring a *3×P3-RFP*, showing both orange-to-red-colored and red-fluorescent eyes. **(E)** The {*TG*,*AD2*} strain contains a *TG* and an *AD2* harboring a *3×P3*-*GFP*, showing both orange-to-red-colored and green-fluorescent eyes. **(F)** The recombination product *2×*{*TG*} strain contains two *TG*s, showing orange-to-red-colored eyes that lack green or red fluorescence. Scale bars: 200 µm. **(G–I)** Experimental design and results for studies of adaptor-mediated recombination efficiency using an initial version of the recombinator strain *R2*. This initial version had suboptimal adaptor insertion efficiency. To improve this, we developed a new version by changing the gRNA target and modifying the homologous arm for adaptor insertion. The efficiency of the new *R2* is shown in Figures 1I**, S1L**. **(G)** Cross scheme for assaying adaptor-mediated recombination efficiency. We assayed the insertion of *AD2* into an example transgene (*TG*: *82C10-LexA*). To account for potential maternal effects on Cas9 and gRNA expression that may influence the efficiency, we conducted all possible reciprocal crosses between F0 parents to generate transheterozygote *TG/R2*. 10 out of the 12 F1 transheterozygous genotypes were crossed with the Cas9 strain for F2, whereas 2 male genotypes without Cas9 expression were omitted (marked as N.A.). **(H)** We calculated the adaptor insertion efficiency for *R2* as the percentage of desired transgene-adaptor animals {*TG*,*AD2*} among all F2 progeny. *Left panels,* Mean ± 95% C.I. weighted average adaptor insertion efficiency (n = 3–19 replicates, each with a parent of indicated F1 genotype crossed to a Cas9 strain). The average efficiency of this *R2* was ∼0–0.5%. The efficiency values of most cross designs were significantly higher than those for natural recombination (binomial test; *: *P* < 0.05, ***: *P* < 0.001, ****: *P* < 10^-6^). *Right panels,* Box-and-whisker plots of *R2* efficiency as measured from individual F1 animals (parent of F2). Adaptor insertion efficiencies did not vary significantly between different cross designs (Kruskal-Wallis one-way ANOVA followed by Tukey’s HSD test; n.s.: *P* > 0.05). **(I)** The distribution of indicated phenotypes among F2 animals. The desired genotype {*TG*,*AD2*} for assaying adaptor insertion efficiency shows a w^−^R^−^G^+^ phenotype. **(J–L)** Experimental design and results showing the phenotypic distribution of F2 progeny during adaptor insertion using the Recombinators *R1* and *R2*. **(J)** Cross scheme for assaying adaptor insertion efficiency, which is the same as that in Figure 1G. The transgene (*TG*) is *82C10-LexA* in *attP40*. **(K, L)** The phenotypic distributions of F2 progeny during the adaptor insertion process using the Recombinators *R1* and *R2*. The desired genotypes for assaying adaptor insertion efficiency are {*AD1*,*TG*} with a w^+^R^+^G^−^ phenotype for *R1* **(K)**, and {*TG*,*AD2*} with a w^−^R^−^G^+^ phenotype for *R2* **(L)**. The distribution corresponds to the experiments in Figure 1H**,I**. **(M–O)** Experimental designs and results for studies of adaptor insertion efficiency in different genomic docking sites. **(M)** Cross scheme for assaying adaptor insertion efficiency. To evaluate adaptor insertion efficiency, we inserted *AD1* or *AD2* into an example transgene (*TG*: *UAS-pAce*) in *attP40, VK27* and *attP2* sites. We crossed male *TG* flies and female Recombinator *R1* or *R2* flies to generate male *TG/R1* or *TG/R2* transheterozygotes for the assay. **(N)** Adaptor insertion efficiency of *R1* and *R2* in genomic docking sites *attP40*, *attP2*, and *VK27*. The adaptor insertion efficiency for *R1* or *R2* is calculated as the percentage of desired transgene-adaptor animals {*AD1*,*TG*} (w^+^R^+^G^−^ phenotype) or {*TG*,*AD2*} (w^−^R^−^G^+^ phenotype) among all F2 progeny, respectively. *Left panels,* Mean ± 95% C.I. weighted average adaptor insertion efficiency (n = 5–7 replicates, each with a parent of indicated F1 genotype crossed to a Cas9 strain). The average efficiency of *R1* was 23%, 21%, and 16%, and that of *R2* was 16%, 13%, and 14%, for *attP40*, *attP2*, and *VK27*, respectively, all significantly higher than that of the near-zero efficiency of natural recombination for generating transgene-adaptor strain (binomial test; ****: *P* < 10^-6^). *Right panels,* Boxplots showing the distribution of adaptor insertion efficiency measured. They were not significantly different among groups, demonstrating consistent and robust adaptor insertion efficiency across various integration sites (Kruskal-Wallis one-way ANOVA followed by Tukey’s HSD test; n.s.: *P* > 0.05). **(O)** The phenotype distributions obtained with each cross design. The desired genotypes for assaying adaptor insertion efficiency are: {*AD1*,*TG*} with a w^+^R^+^G^−^ phenotype for *R1*, and {*TG*,*AD2*} with a w^−^R^−^G^+^ phenotype for *R2*. **(P–R)** Experimental designs and results for studies of adaptor-mediated recombination in the *attP2* site. **(P)** Cross scheme for assaying adaptor-mediated recombination efficiency. To account for potential maternal effects on Cas9 and gRNA expression, we performed reciprocal crosses between F2 flies carrying the transgene-adaptor {*AD1*,*TG*} and {*TG*,*AD2*} (*TG*: *UAS-Ace::mNeon* in *attP2*). We crossed the resulting F3 transheterozygote {*AD1*,*TG*}/{*TG*,*AD2*} males and females with the Cas9 strain for F4 (the same as Figure 1J). **(Q)** Efficiency of adaptor-mediated recombination using each cross strategy of **(P)**. We determined the recombination efficiency as the percentage of the desired animals carrying the IGA *2×*{*TG*} (w^+^R^−^G^−^ phenotype) among all F4 progeny (the same as Figure 1K). *Left panels,* Mean ± 95% C.I. average recombination efficiency (n = 6–7 replicates, each with a parent of indicated F3 genotype crossed to a Cas9 strain). The recombination efficiency were ∼4–30%, significantly higher than the near-zero rate from the natural recombination (binomial test; ****: *P* < 10^-6^). *Right panels,* Boxplots showing the distribution of recombination efficiency measured. The recombination efficiencies among different groups were not significantly different (Kruskal-Wallis one-way ANOVA followed by Tukey’s HSD test; n.s.: *P* > 0.05). **(R)** Phenotype distributions obtained with each cross design. The desired genotypes *2×*{*TG*} for assaying adaptor-mediated recombination efficiency shows a w^+^R^−^G^−^ phenotype. Panels **A–F** show the genotypes and eye phenotypes corresponding to the color-coded phenotypes listed in **I**, **K**, **L**, **O**, and **R**: w^+^, red eye; w^−^: white eye; R^+^/G^+^: eye-specific RFP/GFP fluorescence; R^−^/G^−^: absence of eye-specific RFP/GFP fluorescence.

**Figure S2.**
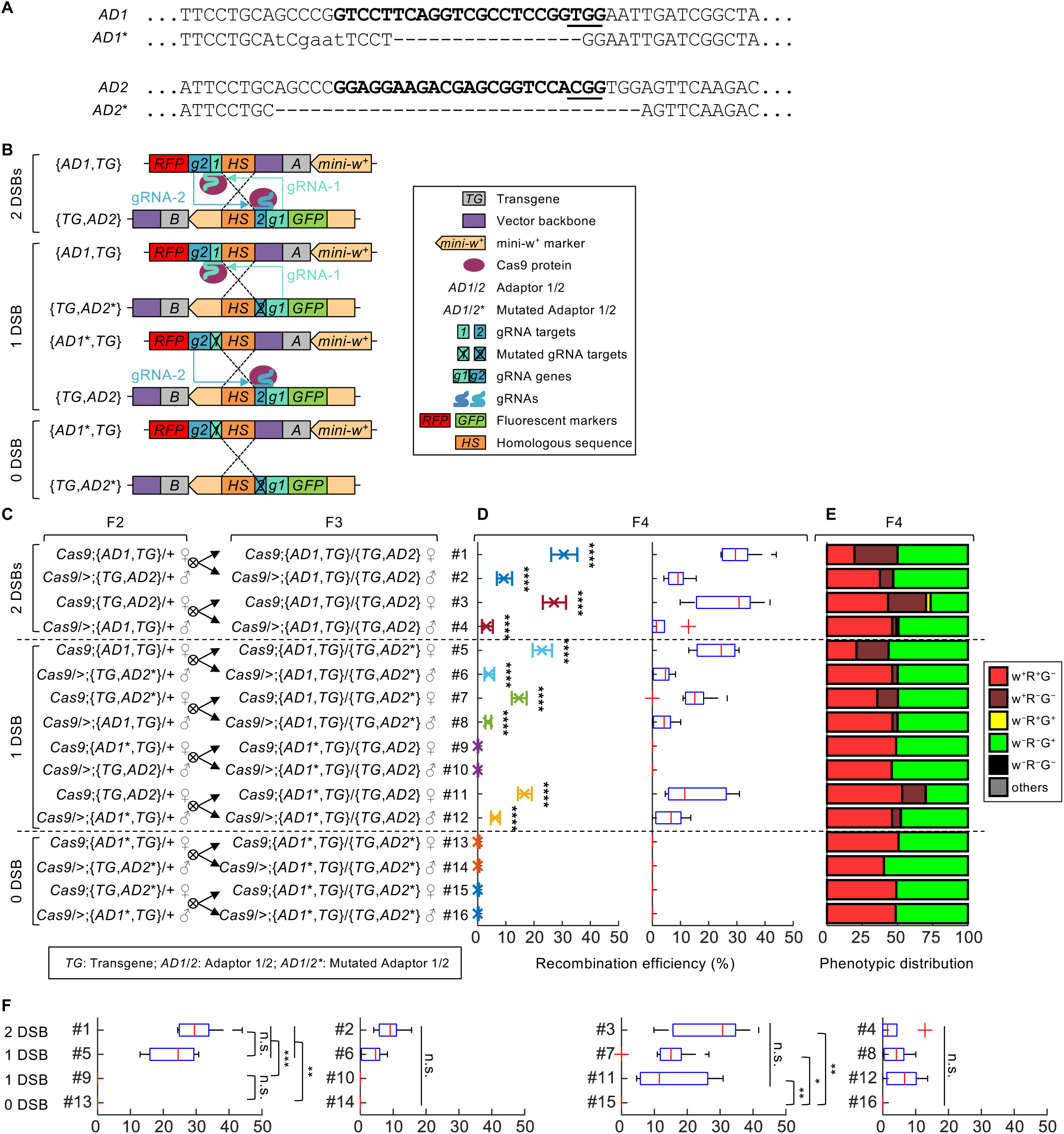
Effects of CRISPR/Cas9-induced double-strand breaks (DSBs) in adaptors on recombination efficiency. **(A)** gRNA target sequences in the original adaptors (*AD1* and *AD2*) and variants with mutated gRNA targets (*AD1*\* and *AD2*\*). The 23-nt gRNA targets are shown in bold, and Protospacer Adjacent Motifs (PAMs) are underlined. Deleted nucleotides are indicated by “–”. Mutated nucleotides are shown in lowercase. These mutations prevent Cas9 from binding and cutting the DNA, effectively disabling the generation of DSBs at these locations. **(B)** Schematic of the experimental design to investigate the number of DSBs required for recombination. Scenarios with 2, 1 or 0 DSBs were generated during adaptor-mediated recombination between two IGA-adaptors by crossing indicated strains. This allowed us to examine whether a single DSB is sufficient to trigger recombination. Our aim was to elucidate the mechanism for the generation of undesired recombination products. Assuming a single DSB can trigger recombination, the adaptor containing the break might recombine with residual homologous sequences present between transgenes lacking the break, leading to the formation of undesired products. Colored rectangles and pentagons denote distinct genetic elements. Magenta ovals denote the Cas9 protein, and thick curves depict gRNAs, of which the colors match those of their corresponding targets and expression genes. **(C–F)** Experimental designs and results for studies of adaptor-mediated recombination efficiency with different numbers (2, 1 or 0) of Cas9/gRNA-induced DSBs. **(C)** Cross scheme for assaying adaptor-mediated recombination efficiency. To account for potential maternal effects on Cas9 and gRNA expression, we performed reciprocal crosses between F2 flies carrying the transgene-adaptor {*AD1*,*TG*} and {*TG*,*AD2*} (*TG*: *82C10-LexA*). We crossed the resulting F3 transheterozygote {*AD1*,*TG*}/{*TG*,*AD2*} males and females with the Cas9 strain for F4 (the same as Figure 1J). **(D)** Efficiency of adaptor-mediated recombination using each cross strategy of **(C)**. We determined recombination efficiency as the percentage of the desired animals carrying the IGA *2×*{*TG*} among all F4 progeny. *Left panels,* Mean ± 95% C.I. weighted average recombination efficiency (n = 5–17 replicates, each with a parent of indicated F3 genotype crossed to a Cas9 strain). The average recombination efficiency with 2 DSBs was ∼3–31%, significantly higher than the near-zero rate from natural recombination (The data for 2 DSBs is the same as Figure 1H. They are included here for ease of comparison). The average recombination efficiency with 1 DSB was ∼0–23%; in particular, the cross scheme with the *AD1* mutation from maternal F2 led to 0% recombination efficiency, whereas the rest of the schemes had efficiencies that were significantly higher than those of natural recombination. The mean recombination efficiency with 0 DSBs was 0. Efficiency values were compared to natural recombination using a binomial test (****: *P* < 10^-6^). *Right panels,* Recombination efficiencies measured for individual F3 animals (parents of F4). **(E)** Phenotype distribution corresponding to each cross design. The desired genotypes *2×*{*TG*} for assaying adaptor-mediated recombination efficiency exhibit a w^+^R^−^G^−^ phenotype. Phenotype abbreviations: w^+^, red eye; w^−^, white eye; R^+^/G^+^, eye-specific RFP/GFP fluorescence; R^−^/G^−^, absence of eye-specific RFP/GFP fluorescence. **(F)** A re-ordered version of the set of box-and-whisker plots of **(D),** grouping together those with similar cross strategies to ease visual comparisons (Kruskal-Wallis one-way ANOVA followed by Tukey’s HSD test; n.s.: *P* > 0.05, *: *P* < 0.05, **: *P* < 0.01, ***: *P* < 0.001).

**Figure S3.**
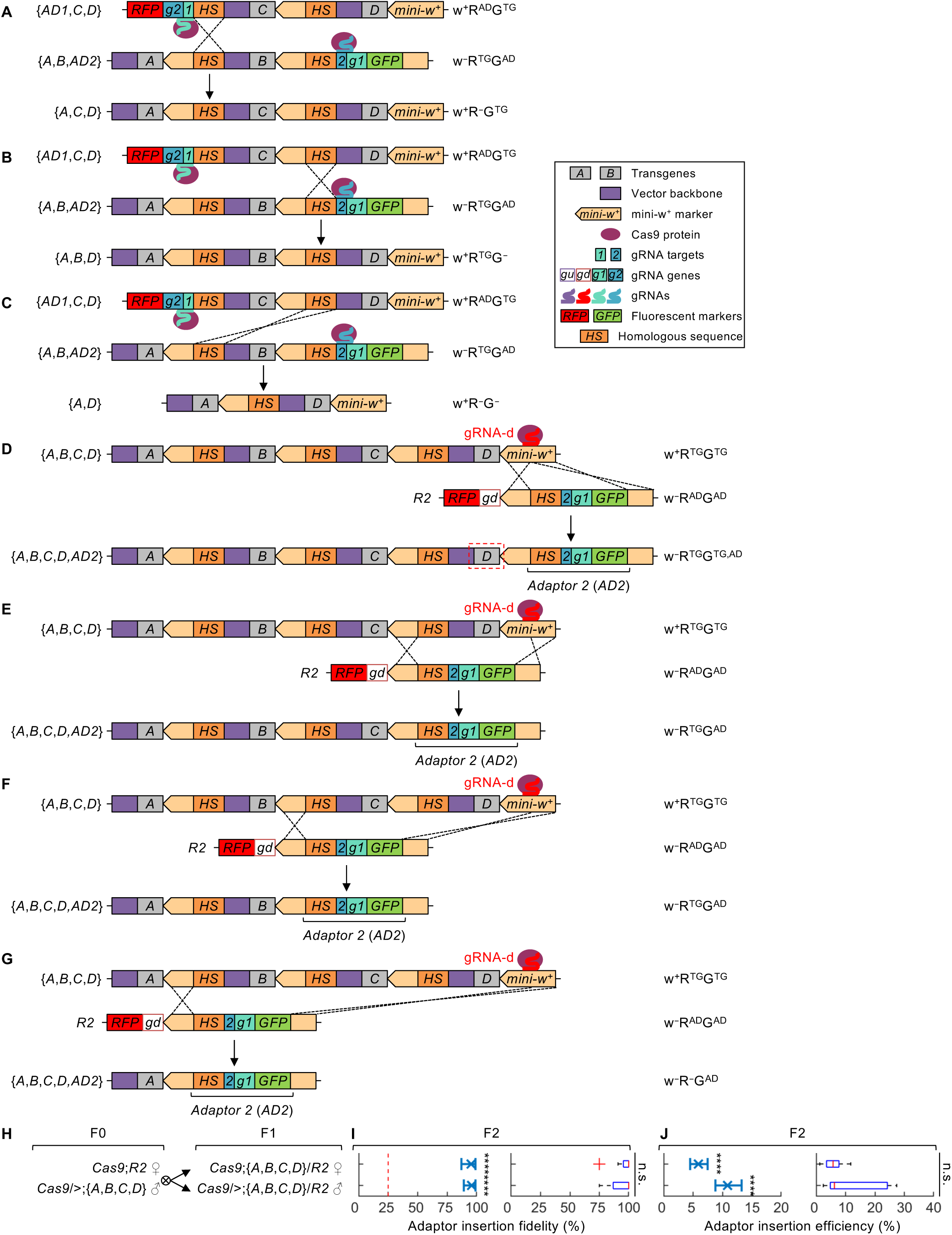
Undesired recombination and adaptor insertion products generated during CRISPR/Cas9-induced homology-directed repair in the SuRe system. **(A–C)** In addition to the desired homology sequence pairing shown in Figure 2A, three other pairing scenarios can occur during adaptor-mediated recombination between {*AD1*,*C*,*D*} and {*A*,*B*,*AD2*}. For simplicity, only shown for each pairing condition are the products selected in the initial screen for the presence of the *mini-w* marker and the absence of both eye-specific adaptor fluorescent markers (*3×P3-GFP* and *3×P3-RFP*). Colored rectangles and pentagons represent distinct genetic elements. Magenta ovals represent the Cas9 protein; thick curves depict gRNAs, of which the colors match those of their corresponding targets and expression genes. Legend also applies to panels **D–G**. **(D–G)** Four pairing scenarios can occur during the adaptor insertion step when inserting *AD2* into the IGA {*A*,*B*,*C*,*D*}. For simplicity, only shown for each pairing condition are products lacking the *mini-w* marker but containing the *AD2* fluorescent marker (*3×P3-GFP*), as these are the products identified after marker screening in the adaptor insertion step. **(D)** The desired homology pairing outcome is the insertion of *AD2* downstream of transgene *D*, creating {*A*,*B*,*C*,*D*,*AD2*}. **(E–G)** Undesired pairings lead to *AD2* insertion downstream of transgene *C*, *B*, or *A*, causing the loss of desired transgenes in the final product. When creating the desired product, both ends of the DNA break created by Cas9/gRNA align perfectly with the template DNA **(D)**. However, in generating the undesired products, only one end of the break achieves perfect alignment **(E–G)**. This inherent asymmetry in the DNA repair process creates a bias favoring the formation of the desired product during adaptor insertion. The red dashed box in **(D)** highlights a key criterion for genotyping: only the desired product retains the transgene adjacent to the DNA break (transgene *D*), while undesired products lose this gene. Following an initial screening using visible markers in the transgene vector and adaptors (selecting for w^−^R^AD–^G^AD+^ flies which have lost the *mini-w* marker and contain *AD2*), PCR genotyping for the presence of the transgene adjacent to the DNA cleavage site (transgene *D*) can identify the desired adaptor insertion product. **(H–J)** Experimental designs and results for studies of adaptor insertion into the IGA {*A*,*B*,*C*,*D*} using SuRe-CC. **(H)** Cross scheme for assaying adaptor insertion into the IGA {*A*,*B*,*C*,*D*} using SuRe-CC. We performed reciprocal cross between the F2 flies carrying the {*AD1*,*C*,*D*} and {*A*,*B*,*AD2*} adaptors. F3 transheterozygote *Cas9*;{*AD1*,*C*,*D*}/{*A*,*B*,*AD2*} males and females were then crossed with Cas9 for F4. Transgene strains *A*, *B*, *C* and *D* are as described in **(A, B)**. **(I)** The fidelity of adaptor insertion into the IGA {*A*,*B*,*C*,*D*} using *R2* of SuRe-CC. *Left panels*, Mean ± 95% C.I. weighted average adaptor insertion fidelity (n = 12, 7). The adaptor insertion fidelity is defined as the percentage of desired animals {*A*,*B*,*C*,*D*,*AD2*} (phenotype w^−^R^TG^G^TG,AD^) among the F2 progeny resulting from the initial screening for the present of adaptor *AD2* marker G^AD^ and the absence of the *mini-w*^+^ (including phenotypes w^−^R^TG^G^TG,AD^, w^−^R^TG^G^AD^, and w^+^R^−^G^AD^ shown in **(E-G)**). If the four adaptor insertion products occur with equal probability, the expected fidelity would be 25% (red dashed line). The adaptor insertion fidelity was significantly higher than 25% (binomial test; ****: *P* < 10^-6^), reaching ∼95–96%, suggesting a significant bias towards the desired product. *Right panels,* Boxplots showing the distribution of adaptor insertion fidelity measured for individual F1 animals (parent of F2). Adaptor insertion efficiencies among different cross designs were not significantly different (Kruskal-Wallis one-way ANOVA followed by Tukey’s HSD test; n.s.: *P* > 0.05). Each assay had at least seven individual F1 animals of the indicated genotype (n = 12 and 7). **(J)** Efficiency of adaptor insertion into the IGA {*A*,*B*,*C*,*D*} using *R2* of SuRe-CC. The adaptor insertion efficiency was determined as the percentage of the desired animals carrying {*A*,*B*,*C*,*D*,*AD2*} among all F2 progeny. *Left panels,* Mean ± 95% C.I. the weighted average efficiency (n = 12, 7). The adaptor insertion efficiency is defined as the percentage of the desired animals carrying {*A*,*B*,*C*,*D*,*AD2*} among all F2 progeny. The average adaptor insertion efficiency was ∼6–11%, which was significantly higher than that of the natural integration (binomial test; ****: *P* < 10^-6^). *Right panels,* Adaptor insertion efficiencies measured for individual F1 animals (parents of F2). Efficiencies did not vary significantly between the different cross designs (Kruskal-Wallis one-way ANOVA followed by Tukey’s HSD test; n.s.: *P* > 0.05). Transgenes were the same as those in Figure 2: *A*: *10×UAS-IVS-myr::tdTomato*, *B*: *R57C10-GAL4*, *C*: *13×LexAop2-mCD8::GFP*, and *D*: *R57C10-LexA*. Phenotype abbreviations in **(A–G)**: w^+^, red eye; w^−^, white eye; R^AD^/G^AD^, eye-specific RFP/GFP fluorescence from the adaptors; R^TG^/G^TG^, RFP/GFP fluorescence from the transgenes; R^−^/G^−^: absence of RFP/GFP fluorescence from both the adaptors and the transgenes. Symbols in **(A–G)**: Colored rectangles and pentagons represent distinct genetic elements. Magenta ovals denote the Cas9 protein. Thick curves depict gRNAs, and their colors are matched with those of their corresponding targets and expression genes.

**Figure S4.**
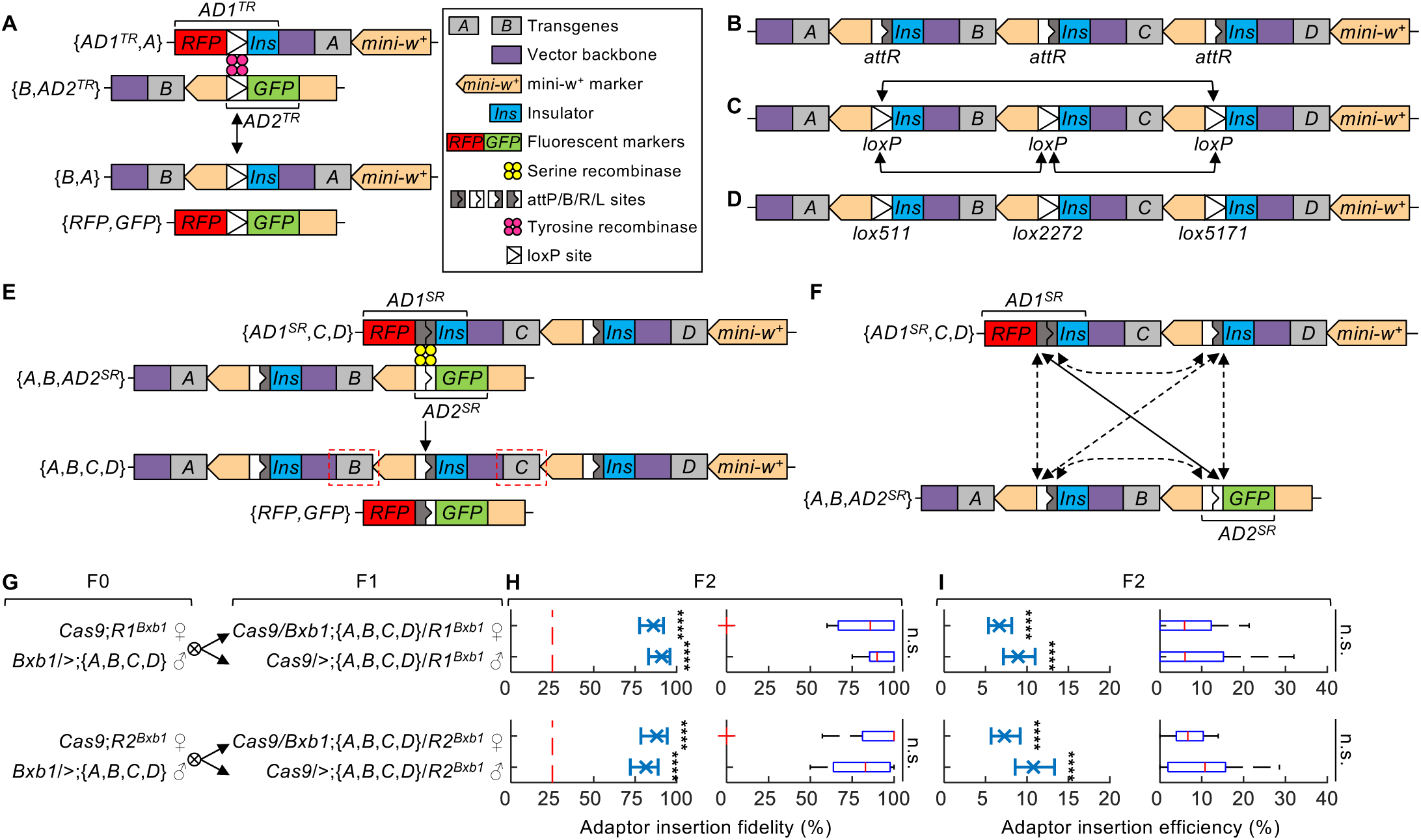
Comparison of serine and tyrosine recombinase usage in the SuRe-CR system. **(A)** Adaptor-mediated recombination in a hypothetical SuRe-CR system with tyrosine recombinase (TR). Both *AD1^TR^* and *AD2^TR^* adaptors contain *loxP* sites, which are targets for tyrosine recombinase. Tyrosine recombinases catalyze a bidirectional reaction, resulting in the retention of *loxP* sites in the recombination product. The hypothetical working mechanism for SuRe-CR with tyrosine recombinase applies to the intended recombination products presented in **C** and **D**, whereas panels **B**,**E–I** follow the working mechanism for SuRe-CR using serine recombinase (Figure 3A). Colored rectangles and pentagons denote distinct genetic elements. Yellow circles represent the serine recombinase protein. The four different types of rectangles with internal white and/or dark gray elements respectively denote the attP, attB, attR, and attL sites. Sets of 4 pink circles denote the tyrosine recombinase protein. Rectangles with triangles inside depict LoxP sites. The legend also applies to panels **B–F**. **(B)** The intended recombination product generated by the serine recombinase version of SuRe-CR has adaptor residuals with *attR* sites between the recombined transgenes. These *attR* sites are not substrates for the serine recombinase, preventing undesired excision of the recombined transgenes. **(C)** The intended recombination product generated by the hypothetical SuRe-CR using tyrosine recombinase has adaptor residuals with *loxP* sites between the recombined transgenes. Due to the presence of these *loxP* sites, the tyrosine recombinase could catalyze recombination between them (shown by black arrow), resulting in excision of the transgenes located between the paired *loxP* sites. **(D)** Multiple mutated *loxP* variants (e.g., *lox511*, *lox2272*, and *lox5171*) can prevent excision in **(C)**. However, each pair of SuRe Recombinator strains can only carry one *loxP* variant. Introducing these different *loxP* variants would require distinct Recombinator strain pairs for their insertion. Consequently, recombining *N* transgenes would necessitate engineering *N* – 1 pairs of Recombinator strains, each carrying a different mutated *loxP* variant. This requirement compromises the reusability of the Recombinator pairs and increases the complexity of the system. **(E)** Illustration of the desired products from further recombination with IGAs using the SuRe-CR system. As with Figure 2A, individual transgenes *A*, *B*, *C*, and *D* are at the same chromosomal locus and share identical flanking sequences. {*AD1^SR^*,*C*,*D*} and {*A*,*B*,*AD2^SR^*} are results from insertion of adaptors to the previous integrated {*C,D*} and {*A,B*} arrays, respectively. {*AD1^SR^*,*C*,*D*} and {*A*,*B*,*AD2^SR^*} contain *attP* and *attB* sites respectively. Products from the recombination between {*AD1^SR^*,*C*,*D*} and {*A*,*B*,*AD2^SR^*} were identified after an initial screening for the absence of eye-specific RFP (R^AD^) and GFP (G^AD^) markers (from the adaptors) and the presence of mini-w^+^ (from the IGA). Red dashed boxes highlight the two transgenes adjacent to adaptors (*B* and *C*). PCR genotyping for the presence of transgenes adjacent to the adaptors (*B* and *C*) can confirm the desired recombination product. **(F)** The desired product {*A*,*B*,*C*,*D*} is generated from pairing of *attP*-*attB* (solid black arrow). Serine recombinase does not catalyze alternative pairings (such as *attP*-*attR*, *attB*-*attR*, or *attR*-*attR*, dashed black arrows), preventing the formation of undesired products and transgene excision. This inherent specificity of serine recombinase ensures high fidelity in the SuRe-CR system. **(G–I)** Experimental designs and results showing the fidelity and efficiency of adaptor insertion into the IGA {*A*,*B*,*C*,*D*} with SuRe-CR. **(G)** Cross Strategies for assaying fidelity and efficiency of adaptor insertion into the IGA {A,B,C,D} using SuRe-CR(Bxb1). **(H)** The fidelity of adaptor insertion into the IGA {*A*,*B*,*C*,*D*} using *R1* or *R2* of SuRe-CR(Bxb1). *Left panels,* Mean ± 95% C.I. weighted average adaptor insertion fidelity (n = 9–17 replicates). Adaptor insertion fidelity was determined as the percentage of desired animals, either {*R1*,*A*,*B*,*C*,*D*} or {*A*,*B*,*C*,*D*,*R2*}, among the progeny initially screened for the presence of the adaptor marker and the absence of *mini-w*^+^. Similarly to **Figure S3D–G**, four potential insertion products are possible after this initial screening. If the four products occur with equal probability, the expected adaptor insertion fidelity would be 25% (red dashed line). The adaptor insertion fidelity was significantly higher than 25% (binomial test; ****: *P* < 10^-6^), reaching ∼86–91% for *R1* and ∼82–88% for *R2*, suggesting a significant bias towards the desired insertion product. *Right panels,* Fidelity values measured for individual F1 animals (parents of F2). Adaptor insertion fidelity did not vary significantly across different cross designs (Kruskal-Wallis one-way ANOVA followed by Tukey’s HSD test; n.s.: *P* > 0.05). Each assay had at least nine individual F1 animals of the indicated genotype (n = 9–17). **(I)** Efficiency of adaptor insertion into the IGA {*A*,*B*,*C*,*D*} using *R1* or *R2* of SuRe-CR(Bxb1). *Left panels,* Mean ± 95% C.I. weighted average adaptor insertion efficiency (n = 9–17 replicates). The average adaptor insertion efficiency was ∼7–11%, significantly higher than the efficiency of natural integration (binomial test; ****: *P* < 10^-6^). *Right panels,* Efficiencies measured for individual F1 animals (parents of F2). All efficiencies among different cross designs are not significantly different (Kruskal-Wallis one-way ANOVA followed by Tukey’s HSD test; n.s.: *P* > 0.05). Each assay had at least nine individual F1 animals of the indicated genotype (n = 9–17). Transgenes were the same as those in Figure 2: *A*: *10×UAS-IVS-myr::tdTomato*, *B*: *R57C10-GAL4*, *C*: *13×LexAop2-mCD8::GFP*, and *D*: *R57C10-LexA*.

**Figure S5.**
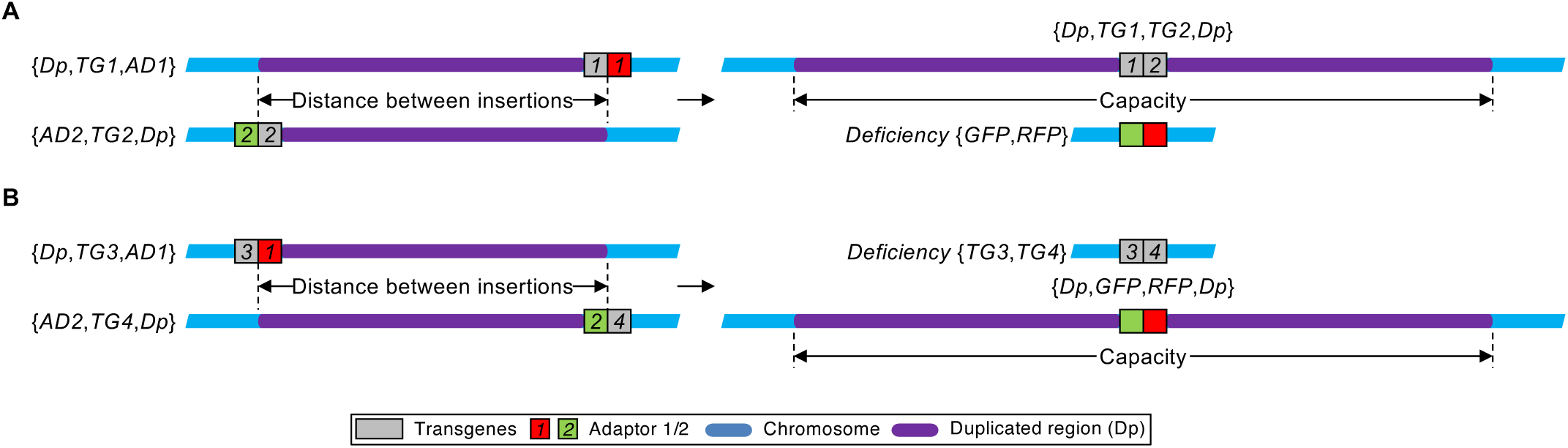
Creating gene duplications and deletions using SuRe. **(A, B)** Recombining *AD1* and *AD2* at distinct genomic loci with SuRe-CC typically yields either a deletion or a duplication, whereas SuRe-CR can produce both outcomes. This difference arises from the distinct mechanisms employed by each system, as illustrated in Figures 1B and **4A**. Although both systems can generate the desired recombination product, SuRe-CC rarely creates the byproduct containing {*GFP*,*RFP*}, because the *GFP* and *RFP* in the adaptors do not connect to the homologous sequence (*HS*) to facilitate their recombination (Figure 1B). In contrast, SuRe-CR can create the byproduct containing {*GFP*,*RFP*} because the *GFP* and *RFP* in the adaptors can be linked by an *attL* site (Figure 4A). **(A)** When *AD1* is downstream of *AD2*, SuRe-CC recombines {*Dp*,*TG1*,*AD1*} with {*AD2*,*TG2*,*Dp*} to generate a duplication strain {*Dp*,*TG1*,*TG2*,*Dp*}. In this scenario, SuRe-CR can generate both the duplication strain {*Dp*,*TG1*,*TG2*,*Dp*} and a deletion strain {*GFP*,*RFP*}. **(B)** When *AD1* is upstream of *AD2*, SuRe-CC recombines {*Dp*,*TG3*,*AD1*} with {*AD2*,*TG4*,*Dp*} to generate a deletion strain {*TG3*,*TG4*}. In this scenario, SuRe-CR can generate both a deletion strain {*TG3*,*TG4*} and a duplication strain {*Dp*,*GFP*,*RFP*,*Dp*}. Gray, red and green rectangles respectively represent transgenes *AD1* and *AD2*. Blue and purple bars respectively represent the chromosome and the duplicated region. Gray, red and green rectangles respectively denote the transgenes, *AD1,* and *AD2*. Blue and purple bars respectively denote the chromosome and the duplicated region.

**Figure S6.**
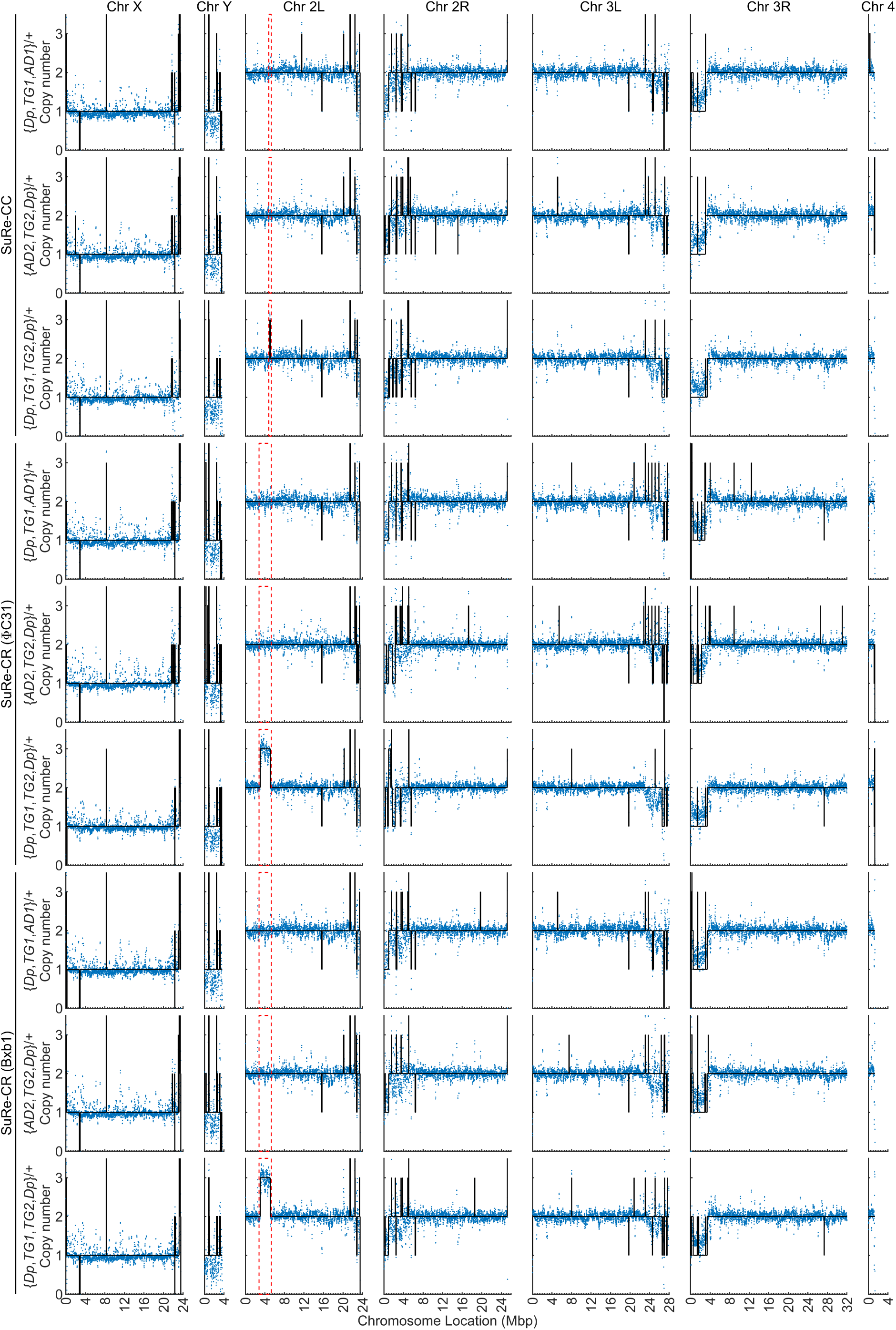
**Whole-genome copy number variation (CNV) analysis confirms successful generation of duplication and adaptor strains.** Full CNV profiles of parental strains carrying adaptors (AD) and the resulting duplication strains generated using SuRe-CC, SuRe-CR(ΦC31), and SuRe-CR(Bxb1) are shown. The x-axis shows the genomic coordinates; the y-axis shows the CNVs, visualized as the normalized sequencing read densities (blue dots) at the corresponding genomic coordinates along the x-axis. Black solid lines mark the estimated copy number inferred by the analysis. Red dashed lines highlight the regions further enlarged and detailed in Figure 5E. The *TG1* is *R57C10-LexA* in *attP40* (25C6). The *TG2* recombined by SuRe-CC is *P{lacW}Tfb5^k10127^*, which is 0.1 Mbp away from *attP40*. The *TG2* recombined by SuRe-CR is *P{lacW}FASN2^k05816^*, which is 2.1 Mbp away from *attP40*. In heterozygous transgene-adaptor strains {*Dp*,*TG1*,*AD1*}/+ and {*AD2*,*TG2*,*Dp*}/+, the CNVs in the DNA segment between *TG1* and *TG2* are the same as those in its surrounding DNA. In the heterozygous duplication strain {*Dp*,*TG1*,*TG2*,*Dp*}/+, the DNA segment between *TG1* and *TG2* increased to 3, one copy more than those in its surrounding DNA. These results confirm the successful duplication of the targeted DNA segment between *TG1* and *TG2*. Note that the DNA samples were from males and thus, the X and Y chromosomes show a copy number of 1.

**Figure S7.**
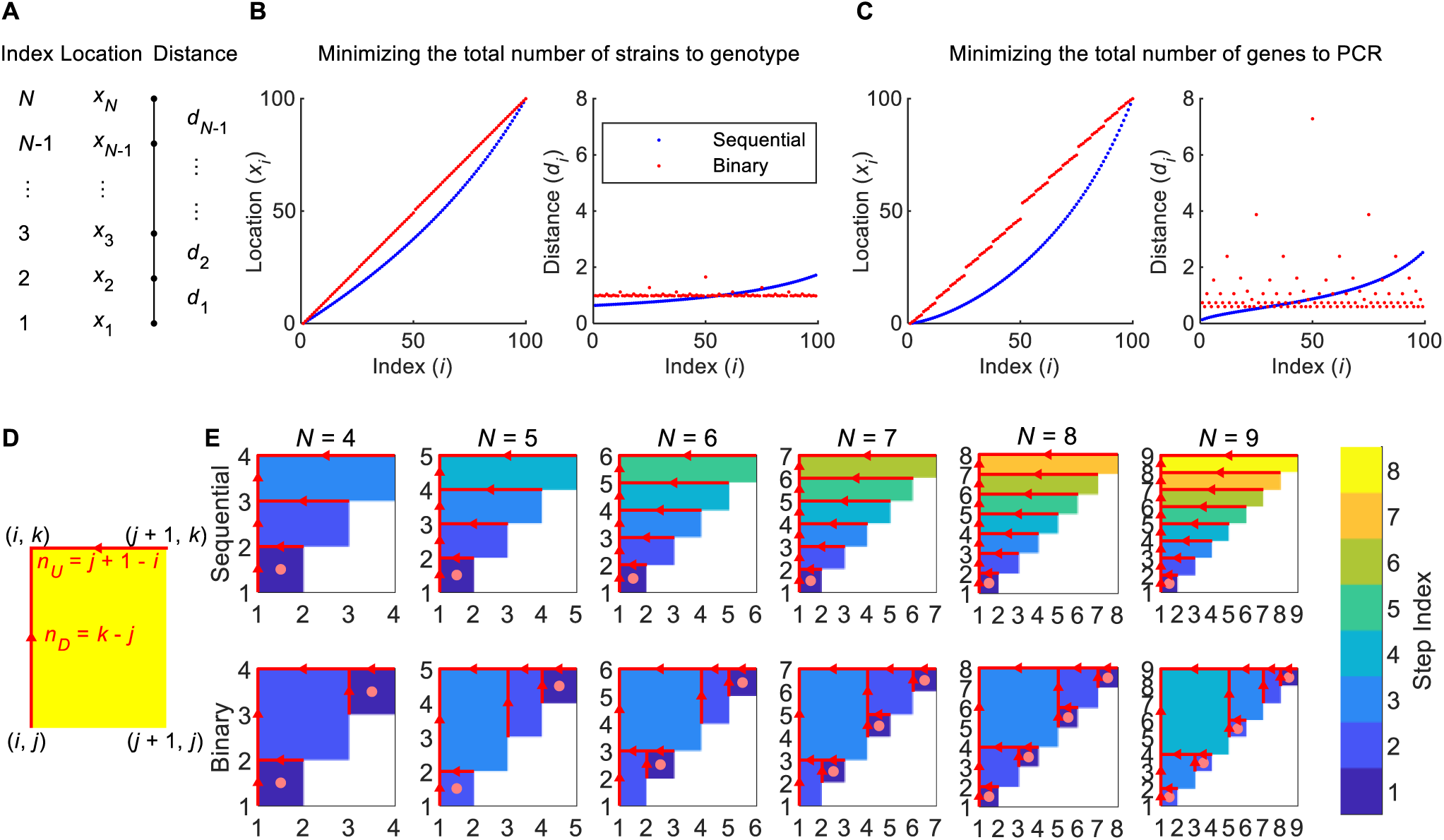
Mathematical modeling of transgene assembly with SuRe or natural recombination. **(A–C)** Modeling of the optimal arrangement of transgene locations on a chromosome (100 cM long) to minimize the screening effort in conventional approaches utilizing natural meiotic recombination. **(A)** Table showing the definition of transgene location along the chromosome (*x_i_*) and the distances between adjacent transgenes (*d_i_*). Both sequential and binary assembly processes are modeled, with optimization focused on minimizing either the number of strains requiring genotyping **(B)** or the total number of genes requiring PCR typing **(C)**. Panels **(B)** and **(C)** show the optimal distribution of transgenes along a chromosome for sequential (blue) and binary (red) assembly processes. Each data point denotes a transgene. *Left panels*, Locations of the transgenes along the chromosome. *Right panels*, Distances separating adjacent transgenes. **(D)** Geometric representation of one recombination step using the SuRe system. In this illustration, the coordinate (*i*, *j*) represents an Integrated Genetic Array (IGA) containing transgenes *i* through *j*. Red arrows, going from (*i*, *j*) and (*j* + 1, *k*) to (*i*, *k*), represent the recombination of the IGA containing transgene *i* through *j* with another IGA containing transgenes *j + 1* through *k*, resulting in a new IGA containing transgenes *i* through *k*. This recombination step is represented by the rectangle with vertices (*i*, *j*), (*j* + 1, *k*), (*i*, *k*), and (*j* + 1, *j*). The width and height of the rectangle are (*j* + 1 – *i*) and (*k* – *j*), respectively, which equal the numbers of transgenes in the two original IGAs (*n_U_* and *n_D_*). The area of this rectangle (*j* + 1 – *i*)(*k* – *j*) = *n_U_ n_D_* is proportional to the number of strains and genes that must be genotyped in this recombination step with the SuRe-CC system. In contrast to SuRe-CC, the number of strains and genes that require genotyping is constant for each recombination step in SuRe-CR. Hence, in this case, the total number of rectangles is proportional to the total number of strains and genes that must be genotyped. Furthermore, the number of strains and genes that require genotyping is constant for each adaptor insertion step in both SuRe-CC and SuRe-CR. In this case, the total number of rectangles is proportional to the total number of strains and genes that must be genotyped. **(E)** A geometric approach to calculate the total number of strains and genes that require genotyping in SuRe-CC and SuRe-CR. The rectangles defined in **(D)** serve as building blocks to represent the entire recombination process, from individual transgenes to the IGA containing transgenes 1 through *N*, where *N* is the total number of transgenes. The various colors of the rectangles indicate the index of the corresponding recombination steps, according to the color lookup table. Initial transgenes are represented by the coordinates (*i*, *i*) where {*i* = 1, 2, …, *N*}, along the diagonal of each plot. Following the red arrows, they are progressively recombined into the product, represented by the coordinate (1, *N*) at the upper left corner of each plot. Each column of panels represents the recombination process for a different value of *N*. The top and bottom rows of panels represent sequential and binary assembly processes respectively. Regardless of the recombination process, the rectangles representing each recombination step collectively fill the upper triangular area of the plot. The total area covered by these rectangles is *N* (*N* – 1) /2, and the total number of the rectangles is *N* – 1. Consequently, in SuRe-CC, the asymptotic numbers of strains and genes requiring genotyping for the recombination steps are both proportional to *N*(*N* – 1)/2, whereas in SuRe-CR, they are both proportional to *N*, independent of the recombination process. The asymptotic numbers of strains and genes requiring genotyping for the adaptor insertion steps are the same for both SuRe-CC and SuRe-CR. They are also proportional to the total number of rectangles (*N* – 1). In summary, the asymptotic numbers of strains and genes requiring genotyping in the SuRe-CC system scale with *N*^2^, while in SuRe-CR they scale with *N*. When using SuRe-CC with male F3 parents during the recombination, the geometric approach may introduce errors due to the rectangles highlighted by the pink dots. Typically, we estimate the fidelity of recombination with the SuRe-CC system for male F3 as 2/(*n_U_n_D_*). However, when *n_U_* = *n_D_* = 1, the fidelity for male F3 is 1, not 2/(*n_U_n_D_*) = 2. Therefore, the rectangles with an area of 1 (highlighted by pink dots) introduce these errors. Fortunately, the number of such rectangles does not exceed *N*/2. As *N* increases, this error becomes negligible relative to the total number, which scales with *N* ^2^.

## Methods

### Drosophila stocks

The *MB080C-split-GAL4*, *MB085C-split-GAL4*, *MB083C-split-GAL4* and *MB011B-split-GAL4* strains were kindly gifted by the FlyLight Project Team at Janelia Research Campus^51^. The *R82C10-LexA* (RRID:BDSC_54981)^29^, *10×UAS-IVS-myr::tdTomato* (RRID:BDSC_32222)^29^, *13×LexAop2-mCD8::GFP* (RRID:BDSC_32205)^29^, *R57C10-GAL4* (RRID:BDSC_81088)^25^, *R57C10-LexA* (RRID:BDSC_52817)^29^, *P{lacW}Tfb5^k10127^*(RRID:BDSC_10973)^86^, *P{lacW}Scox^SH1783^*(RRID:BDSC_29505)^86^, *P{lacW}ed^k01102^*(RRID:BDSC_10490)^86^, *P{lacW}FASN2^k05816^* (RRID:BDSC_10580)^86^, *P{lacW}dbe^k05428^* (RRID:BDSC_12169)^86^, *R14C08-GAL4* (RRID:BDSC_48606)^25^, *R52G04-GAL4*(RRID:BDSC_38843)^25^, *Act5C-Cas9*,*lig^4169^* (RRID:BDSC_58492)^87^, *nos-ΦC31*;*CyO/Sco* (RRID:BDSC_34770)^35^, *nos-ΦC31*;;*Dr^1^/TM3,Sb^1^*(RRID:BDSC_34771)^35^, *vas-Bxb1* (RRID:BDSC_67104)^36^, *vas-TP901-1* (RRID:BDSC_80078)^37^, *Act5C-Cas9(RFP+)*;*CyO/Sco* (RRID:BDSC_90363)^87^, *U6:3-GFP.gRNA* (RRID:BDSC_81897)^88^, *UAS(loxP.lexA)QF*;*TM3,Sb^1^*/*TM6B,Tb^1^* (RRID:BDSC_93912)^89^, *13×LexAop2-IVS-GCaMP6f-p10* (RRID:BDSC_44277)^53^ and *CyO*/*Sco*;*MKRS*/*TM6B*,*Tb1* (RRID:BDSC_3703) were obtained from Bloomington Drosophila Stock Center (BDSC). We have previously published descriptions of the voltage indicator strains *26×LexAop2*-*Ace::mNeon* (RRID:BDSC_90342)^41^, *13×LexAop2*-*VARNAM2* (RRID:BDSC_94206)^42^, and *20×UAS*-*Ace::mNeon2* (RRID:BDSC_94209)^42^. To prevent interference from the GFP marker in the *vas-TP901-1* strain (RRID:BDSC_80078)^37^ during selection, we used *Act5C-Cas9(RFP+);CyO/Sco* (RRID:BDSC_90363)^87^ and *U6:3-GFP.gRNA* (RRID:BDSC_81897)^88^ to mutate the GFP marker. Successful mutation was confirmed by PCR sequencing. We donated the *vas-TP901-1(GFP-)* strain to BDSC (RRID:BDSC_605091). Flies were raised at 25°C in 12 hr light/dark cycles on standard food at 50% humidity.

### *C. elegans* stocks and maintenance

We maintained *C. elegans* on nematode growth media (NGM) plates seeded with OP50 *E. coli* following standard protocols^90^. All transgenic strains were constructed in the wild-type N2 Bristol genetic background^90^. Imaging studies used L4 animals at 20°C. CRISPR injections were done with day 1 adult animals at 23°C.

### Generation of the constructs for transgenic *Drosophila*

To generate *R1* and *R2* transgenic strains for the SuRe-CC system, we created *R1* and *R2* constructs by PCR cloning the *3×P3-DsRed* from pHD-DsRed (Addgene#51434); *GFP* from UASpBacFPN (DGRC#1287); and *gypsy/Su(Hw)* insulator, *U6:3* promoter and gRNA scaffold from pCFD5 (Addgene#73914). We inserted these elements into the vector pBPLexA::p65Uw (Addgene#26231).

To choose the gRNA used in *R1* and *R2*, we predicted the efficiency of the gRNAs using the CRISPR Efficiency Predictor (https://www.flyrnai.org/evaluateCrispr/)^91^. We confirmed that the gRNAs used in the constructs do not have off-target effects by using flyCRISPR Target Finder (https://flycrispr.org/target-finder/)^92,93^. The exception was that the gRNA targeting to mini-white marker in the vector also targets the endogenous white gene on the X chromosome. This is unavoidable since mini-white is highly homologous to the white gene. Even if there are any off-target effects not predicted by the flyCRISPR Target Finder, we can backcross recombination products with the control flies, removing off-target mutants. Because the genes in the integrated genetic array generated by the SuRe system do not segregate during the backcross, the genetic background cleaning is easy and straightforward. The sequences of gRNAs used in *Drosophila* are in **Table S1**.

Based on the constructs in the SuRe-CC system, we replaced their adaptor cassettes and fluorescent markers to generate the constructs for making the *R1* and *R2* transgenic fly in the SuRe-CR system. We designed synthetic *2×r4* and *3×TpnC41C* promoters to drive expression of fluorescent markers and used them in SuRe-CR(Bxb1) and SuRe-CR(TP901-1) respectively. Their distinct expression pattern in specific tissues allowed us to distinguish the adaptors used in SuRe-CR(Bxb1) and SuRe-CR(TP901-1) from those in SuRe-CC and SuRe-CR(ΦC31), which use the *3×P3* promoter. Compared to endogenous promoters, these two synthetic promoters have multiple copies of the transcription factor binding sites to enhance expression and remove non-binding regions, yielding compact synthetic promoters for which the size reduction improves adaptor insertion efficiency. The *2×r4* promoter, comprising two tandem copies of the *r4* promoter from the *Drosophila Yp* gene^94^, drives strong expression in the fat body, mainly in the abdomen. The *3×TpnC41C* promoter, containing three copies of the MEF2 and SM3 binding sites from the *Drosophila TpnC41C* gene’s promoter^95^, drives strong expression in the flight muscles, mainly in the thorax.

The *2×r4* promoter sequence is:

CTAACTTCCGTGACCCGttaaaataatcaggcgTAGAttaaaataatcaggcgGTCAttaaaataatcaggcgGAGAttaaaataatcaggcgATGCATttaaaataatcaggcgTAGAttaaaataatcaggcgGTCAttaaaataatcaggcgGAGAttaaaataatcaggcgAGCGGAGACTCTA

As indicated by lowercase letters, “ttaaaataatcaggcg” is the r enhancer element sequence^94^.

The *3×TpnC41C* promoter sequence is:

CTAACTTCCGTGACCCGttcacaaataccatttCCctaaaaataaCCttcacaaataccatttCCctaaaaataaCCttcacaaataccatttCCctaaaaataaAGCGGAGACTCTA.

As indicated by lowercase letters, “ctaaaaataa” is the MEF2 binding sequence and “ttcacaaataccattt” is the SM3 binding sequence^95^.

The CFP coding sequence, *2×r4* promoter, *3×TpnC41C* promoter, and *attP* and *attB* sites of ΦC31, Bxb1 and TP901-1 were synthesized by Genscript Inc. Most of the molecular cloning was performed by Genscript Inc.

### Generation of transgenic *Drosophila*

We generated transgenic fly strains for the recombinator strains *R1* and *R2* of SuRe-CC, SuRe-CR(Bxb1), and SuRe-CR(TP901-1) in docking sites *attP40*, *attP2*, and *VK27* using the standard ΦC31 recombinase system^35,96^. However, constructing *R1* and *R2* of the ΦC31 version of SuRe-CR posed a challenge using the ΦC31 system. Since the *attP* site in *AD1^ΦC31^* and the *attB* site in *AD2^ΦC31^* are both substrates for ΦC31 integrase, using this enzyme for transgenesis would lead to unintended recombination events. If ΦC31 integrase is used to generate the *R2* transgenes of SuRe-CR(ΦC31), the *attB* site in the *AD2^ΦC31^* competes with the *attB* in the vector plasmid, leading to about a half of the transgenes lacking an intact *AD2^ΦC31^* cassette. We identified the correct *R2* transgenic strains of SuRe-CR(ΦC31) by PCR. Additionally, the *attP* in the *AD1^ΦC^*^31^ can react with the *attB* in the *R1* plasmid of SuRe-CR(ΦC31), hindering the insertion of the *R1* plasmid of SuRe-CR(ΦC31). To circumvent this issue and achieve site-specific integration for the *R1* of SuRe-CR(ΦC31), we employed CRISPR/Cas9-mediated transgenesis at the *attP40*, *attP2*, and *VK27* loci. Both ΦC31- and CRISPR/Cas9-mediated transgenesis were performed by commercial services from BestGene Inc.

### Measurements of adaptor insertion and recombination efficiency

When measuring adaptor insertion and recombination efficiency, we crossed individual F1 or F3 flies with balancer flies. Thus, we determined the efficiency of each cross as the fraction of desired progeny from individual F1 or F3 parents. This experimental design allowed us to assess the variability in efficiency among individuals. Given that expression levels of Cas9 and gRNA vary in the F1 animals, we made a small improvement in the standardized adaptor insertion protocol of *R2*. That is, picking F1 flies whose complex eyes are almost white to produce the F2 progeny. Since the gRNA of *R2* targets the *mini-white* marker, the eye color of the F1 flies is mosaic. The fraction of the white part in the eye indicates the expression level of Cas9 and gRNA. If a large fraction of the eye of an F1 fly is white, it is more likely to produce the transgene with *AD2* insertion in its F2 progeny. However, the gRNA of *R1* does not target a visible maker, so we cannot use the same approach to select its corresponding F1 flies.

### Standard protocol to create a transgene with adaptors

When using the SuRe system to insert adaptors into transgenes, it is not necessary to take individual male or females to cross with balancer flies as we did for measuring the adaptor insertion efficiency. We tested all 6 combinations of the Cas9, recombinator, and target transgene (**Figure 1G**), and found their efficiency to be acceptable (>1%). In our standardized cross protocol, we crossed female *Act5C-Cas9*;*R1* or *Act5C-Cas9*;*R2* with the male transgenic strain and then picked male F1 to cross with the *Act5C-Cas9* strain or one of the recombinase strains (*nos-ΦC31*, *vas-Bxb1*, or *vas-TP901-1(G*^−^)). Notably, for the SuRe-CC system, we selected the *Act5C-Cas9* in the second cross. Whe for the SuRe-CR system, we chose the corresponding recombinase strain. This approach has two significant advantages. First, it saves time by eliminating the need to combine the *Act5C-Cas9* element with the transgene before crossing with *R1* or *R2*. Second, it enhances the convenience of subsequent recombination steps by enabling easy switching of the *Act5C-Cas9* or the recombinase transgene onto the X chromosome.

SuRe-HACK facilitates the conversion of a *GAL4* transgene to a *p65.AD* hemidriver with concurrent *AD2^Bxb1^* adaptor insertion (**Figure 6I**). Its protocol is analogous to that of the *R2* recombinator in SuRe-CR(Bxb1). A typical Janelia *split-GAL4* strain comprises an *X-p65.AD* hemidriver integrated in the *attP40* or *VK27* locus and a *Y-GAL4.DBD* hemidriver in the *attP2* locus. To co-localize both hemidrivers at *attP2*, we obtained the corresponding *X-GAL4* strain in *attP2* from the Bloomington Drosophila Stock Center (BDSC). We used SuRe-HACK to convert this *X-GAL4* to *X-p65.AD* in *attP2* and then recombined it with the existing *Y-GAL4.DBD* in the same *attP2* locus. Specifically, we crossed female *Act5C-Cas9*;;*R2^HACK.p^*^65^*^.AD^*with male *X-GAL4* flies. Male F1 progeny were selected and crossed with flies carrying the Bxb1 recombinase-expressing gene. From the resulting F2 progeny, flies with genotype {*X-p65.AD*,*AD2^Bxb1^*} were chosen for further recombination with {*AD1^Bxb1^*,*Y-GAL4.DBD*} flies, which we generated by the standard SuRe-CR(Bxb1) protocol.

### Generation of transgenes with mutated adaptors

We generated mutated adaptor strains *Act5C-Cas9*;{*AD1\**,*TG*} and *Act5C-Cas9*;{*TG*,*AD2\**} by the following approach. Here, *TG* is the transgene *R82C10-LexA.* First, we crossed the *Act5C-Cas9*;{*AD1*,*TG*} and *Act5C-Cas9*;{*TG*,*AD2*} to generate the transheterozygous strain *Act5C-Cas9*;{*AD1*,*TG*}/{*TG*,*AD2*} and crossed it with *Act5C-Cas9*;*Sco*/*CyO*. In the transheterozygote, some *AD1* or *AD2* cassettes, which were cleaved by gRNA2 or gRNA1, did not undergo the recombination. Instead, their double-strand breaks (DSBs) were repaired by Non-Homologous End Joining (NHEJ), resulting in indels at the gRNA target sites. We selected individual w^+^RFP^+^GFP^−^ males or w^−^RFP^−^GFP^+^ males from the progeny of *Act5C-Cas9*;{*AD1*,*TG*}/{*TG*,*AD2*} and crossed them to *Act5C-Cas9*;*Sco*/*CyO* to establish candidate mutant strains. Finally, we sequenced the gRNA targets in these candidates and selected the ones with the largest deletion at the gRNA targets as *Act5C-Cas9*;{*AD1\**,*TG*} and *Act5C-Cas9*;{*TG*,*AD2\**} (**Figure S2A**). This experiment also highlights that the w^+^RFP^+^GFP^−^ or w^−^RFP^−^GFP^+^ flies in the F4 progeny are not the same as the {*AD1*,*TG*} and {*TG*,*AD2*} in the F2. Thus, w^+^RFP^+^GFP^−^ or w^−^RFP^−^GFP^+^ flies in the F4 progeny could not be reused for recombination.

### Genotyping of *Drosophila* strains

To genotype fly strains, we extracted genomic DNA with the G-spin™ Total DNA Extraction Mini Kit (iNtRON Biotechnology). We amplified the sequence to be detected using the REDExtract-N-Amp™ PCR ReadyMix™ kit (Sigma-Aldrich). The PCR amplification involved annealing at 62°C for 20 sec and elongating at 72°C for 2 min, for 35 cycles. **Table S2** lists the PCR primers for genotyping.

### *C. elegans* genome editing

We generated single-copy insertions via CRISPR-Cas9 through gonadal microinjection of Cas9 (IDT), gRNA (IDT), tracrRNA(IDT), *dpy-10* oligonucleotide repair template (IDT), and PCR-generated repair templates. All CRISPR experiments were performed using *dpy-10* co-CRISPR as previously described^97^. Plasmids containing the relevant transgenes, with 500 bp homology arms to the insertion site, were synthesized by GenScript. All plasmids were designed with a restriction site to produce a linear repair construct, but ethanol-precipitated PCR products were found to yield higher CRISPR efficiency. In brief, repair constructs were amplified from plasmids via PCR (600 µL total reaction volume) and ethanol precipitation was performed with pure ethanol (Sigma-Aldrich) and 3M sodium acetate (Thermo Scientific). PCR products were incubated with ethanol/NaOAc for 30 min at –80°C, centrifuged for 30 min at 4°C, washed with ice-cold 70% ethanol and centrifuged for 5 min at 4°C, air dried for 15 min, and resuspended in 30 µL water.

Microinjection mixtures were prepared fresh on the day of injection. In brief, gRNAs and tracrRNA were mixed and annealed, then incubated with Cas9 protein for 10 min. Repair templates were melted for 10 min, then added to the gRNA/Cas9 mix along with the *dpy-10* co-CRISPR repair template^97^. The final microinjection mix was centrifuged for 20 min at room temperature and was kept at room temperature during the injection^97^. **Table S1** lists the gRNAs used in this study, **Table S2** has the primers used for PCR genotyping, and **Table S3** lists the *C. elegans* strains that we generated.

### Imaging studies of *C. elegans*

We imaged *C. elegans* at room temperature using live, immobilized animals. Hermaphrodites at the L4 stage were anesthetized with 10 mM levamisole (Sigma-Aldrich) and mounted on 4% agarose pads. We obtained the images of **Figure 3C–E** and **J** using a Zeiss Axio Observer Z1 microscope controlled by 3i Slidebook (V6) software and equipped with a C-Apochromat 40×/0.9 NA water immersion objective lens, a Yokogawa CSU-W1 spinning disk unit, and a Prime 95B Scientific CMOS camera. We obtained the images of **Figure 3G,H** using a Zeiss Axio Observer Z1 microscope controlled by MetaMorph software (version 7.8.12.0) and equipped with a Yokagawa CSU-X1 spinning disk unit, a a plan-apochromat 100×/1.4 oil NA immersion objective lens, and a Hamamatsu EM-CCD digital camera. We performed image processing and quantification using Fiji.

### Generation of transgenes with *AD2* for testing the capacity of SuRe

To test the capacity of different versions of the SuRe system, we create a series of transgenes carrying *AD2* ({*AD2*,*TG2*}) located at progressively distant genomic loci and recombined it with a transgene carrying *AD1* ({*TG1*,*AD1*}) in the *attP40* site (**Figure 5B**). The *R2* produced by all versions of the SuRe systems target the *mini-w* marker, allowing for easy identification of suitable transgenes at varying distances for *AD2* insertion. We selected transgenes with a *mini-w* marker oriented identically to the homology arms in *R2* to ensure proper *AD2* insertion (**Figures 1B** and **4A**). This consistent orientation allowed recombination between *AD1* and *AD2*, facilitating the formation of duplications. The *TG2*s we selected for the capacity experiment were *P{lacW}Tfb5^k10127^* (RRID:BDSC_10973), *P{lacW}Scox^SH1783^* (RRID:BDSC_29505), *P{lacW}ed^k01102^* (RRID:BDSC_10490), *P{lacW}FASN2^k05816^* (RRID:BDSC_10580), *P{lacW}dbe^k05428^* (RRID:BDSC_12169), situated at distances of 0.1 Mbp, 0.14 Mbp, 1.1 Mbp, and 2.1 Mbp, respectively, from the *attP40* site^86^.

### Whole genome sequencing for detecting the duplication in *Drosophila*

We extracted genomic DNA from fly strains using the G-spin™ Total DNA Extraction Mini Kit (iNtRON Biotechnology). To guarantee that the amount of genomic DNA from each strain was not <1.5 μg, we took ∼60 male flies and used two G-spin columns to extract their genomic DNA. Male flies were used for whole-genome sequencing because their single X and Y chromosomes provide an internal control for copy number analysis of autosomes and duplicated regions. The genomic DNA samples were made into a PCR free library and then sequenced with the DNBSEQ PE100 platform. We mapped the sequencing data to the *Drosophila melanogaster* genome (Release 6 plus ISO1 MT, GCF_000001215.4)^98^ For each sample, we obtained at least 4.5 Gb clean data yielding at least 29× coverage of the genome. To analyze copy number variations (CNVs), we used Control-FREEC software^99^ using a 30-kb window with 10-kb sliding step. The 30-kb window dictates that sequencing read counts were aggregated within consecutive 30,000 base-pair segments along the genome. The 10-kb sliding step indicates that the starting position of each subsequent window advanced by 10,000 base pairs relative to the previous one, resulting in a 20-kb overlap between adjacent windows.

### Imaging fluorescent markers in *Drosophila*

We manually sorted flies using fluorescent markers that we visualized under a stereomicroscope equipped with SFA-RB and SFA-GR fluorescence adaptors (Nightsea Inc.). We imaged the samples in **Figure S1A–F** with a Zeiss Axio Zoom.V16 stereomicroscope, and those in **Figures 2C** and **6E** with a Zeiss LSM 700 confocal laser-scanning microscope equipped with either a 10×/0.8 Plan-Apochromat DIC objective (**Figure 2B**) or a 20×/0.8 Plan-Apochromat DIC objective (**Figure 6E**). We processed images with ZEN (Zeiss Inc.) and MATLAB R2022a (Mathworks Inc.) software.

### Fluorescence voltage and Ca^2+^ imaging of neural activity

We performed fluorescence voltage imaging as detailed in our prior publications^41,100^. Ca^2+^ imaging studies involved the same fly preparation and imaging equipment as those for voltage imaging, with the key distinction being the slower frame acquisition rate used for Ca^2+^ imaging. All imaging experiments used female flies (7–14 days old at the time of laser surgery). To prepare flies for voltage or Ca^2+^ imaging experiments, we first opened an imaging window on the fly head using a laser microsurgery system based on a 193-nm-wavelength excimer laser (GamLaser; EX5 ArF), as \ in our prior work^100^. We imaged neural voltage dynamics with a custom-built upright epi-fluorescence microscope equipped with a 1.0 NA water-immersion 20× XLUMPlanFLN objective lens (N20X-PFH, Olympus) and a 360–780 nm wavelength solid-state light source (Spectra X, Lumencor). To image the green fluorescent indicators Ace::mNeon, Ace::mNeon2, and GCaMP6f, we used a 475/28-nm excitation filter (10-10508, Lumencor), a 515-nm dichroic mirror (T515lpxr, Chroma) and a 531/40-nm emission filter (FF01-531/40-25, BrightLine). To image the red fluorescent voltage indicator VARNAM2, we used a 555/28-nm excitation filter (10-10579, Lumencor), 585-nm dichroic mirror (T585lpxr, Chroma), and 641/75-nm emission filter (FF02-641/75-25, BrightLine). The illumination intensity was approximately 3–7 mW/mm^2^ at the specimen plane. We acquired images at 1000 Hz and 30 Hz for voltage and Ca^2+^ imaging, respectively, using a scientific-grade sCMOS camera (Zyla 4.2, Andor) and 2 x 2 pixel binning (**Figure 6C,F,K**).

In the Ca^2+^ imaging studies (**Figure 6K–M**), we tested flies’ responses to an attractive odor, 100% apple cider vinegar (ACV; Bragg) and a repellent odor, benzaldehyde (BEN; CAS 100-52-7, Sigma-Aldrich; dissolved in mineral oil (3% v/v). We delivered these odorants to the flies’ antennae using the same custom olfactometer and airflow parameters as in our prior research, with100 a constant 200 mL/min airflow. To do this, we flowed air through either mineral oil (baseline), ACV (attractive odor) or mineral oil with dissolved BEN (repellent odor). A probe needle directed the airflow to the fly at a 45-deg angle on its right side in the horizontal plane, ∼3 mm from the fly’s antenna. We imaged neural Ca^2+^ dynamics through the transparent window created by laser microsurgery. Each animal underwent three 203-s-long trials, with a rest of at least 3 min between trials. Each trial included a 5-s-long baseline period, three 3-s-periods of odor delivery (*i.e.*, ACV or BEN were each presented three times per trial), and 30-s-intervals of rest between odor deliveries.

### Statistical analysis

We estimated the efficiency and fidelity of SuRe at the adaptor insertion and recombination steps through maximum likelihood estimation. Here, we define *p_i_* as the efficiency or fidelity of the experimental condition *i*, and *p_ij_* as the efficiency or fidelity of the experimental condition *i* and biological repeat *j*. In each such condition, we observed *N_ij_* progeny and found that *n_ij_* out of them were the desired recombination products. One biological repeat here means crossing an individual male or female with a wildtype or balancer strain and sorting their progeny by phenotypes. The number of desired recombination products followed a binomial distribution Binorm(*N_ij_*, *p_i_*). The likelihood of the experimental data is:

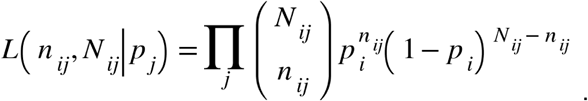

The maximum likelihood estimate of *p_i_* is then:

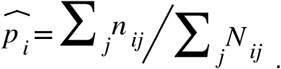

Along the same lines, the maximum likelihood estimate of *p_ij_* is:

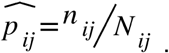

The estimate of *p_i_* equals the weighted average of the estimate of *p_ij_*, which has a weight of 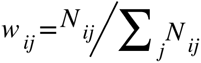. Therefore, we determined the estimate of *p_i_* as the weighted average of the efficiency or fidelity.

We calculated 1-*α* confidence interval of *p_i_* using the Clopper-Pearson method^101^.

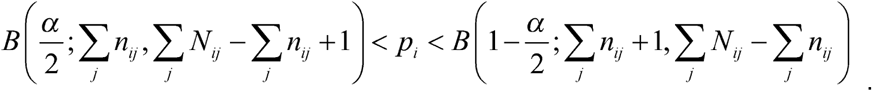

Here, the function *B*(*x*;*b*_1_,*b*_2_) is the inverse cumulative distribution function of a Beta distribution with shape parameters *b*_1_ and *b*_2_. We performed maximum likelihood estimation and calculated the confidence interval using the binomial parameter estimation function *binofit()* in MATLAB (version R2022a, Mathworks).

To evaluate the efficiency and fidelity of our system, we used the binomial test to compare the efficiency with the natural recombination rate, and to compare the measured fidelity with its expected value under the assumption that all possible recombination products occur with equal probability. Let *π_i_* represent the natural recombination rate or the expected fidelity of experimental condition *i*. We tested the null hypothesis *H*_0_: *p_i_* ≤ *π_i_*using a one-sided binomial test. The *P*-value of this test is:

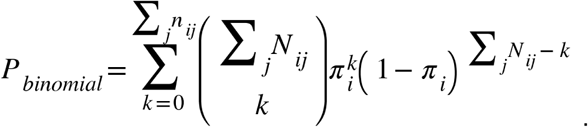

To compare the efficiency to the natural recombination rate, we calculated the natural recombination rate *π_i_* as the length of the homologous sequence (*L_HS_*), which refers to the region of DNA shared between the recombining elements. In *D. melanogaster*, males exhibit no meiotic recombination (0 cM/Mbp), while females have a recombination frequency of 2.46 cM/Mbp^102,103^. Consequently, for males, *π_i_* = 0; for females, *π_i_* = *L_HS_*× 2.46 × 10^-8^, where the unit of *L_HS_* is bp. We then applied this calculation to both adaptor insertion and recombination steps of the SuRe system.

During the adaptor insertion step, the sequence between the adaptor insertion site and the edge of the target transgene serves as a homology arm for CRISPR/Cas9 induced homology-directed repair. In the absence of CRISPR/Cas9 cutting, natural recombination must occur in this region to insert the adaptor adjacent to the transgene. Hence, we used the distance between the adaptor insertion site and the edge of the target transgene as *L_HS_*to calculate the probability of natural recombination-mediated adaptor insertion in females: for *R1*, *L_HS_* = 1813 bp, corresponding to *π_i_* = 4.46 × 10^-5^ (used for the binomial test in **Figures 1H, S1N** and **S4I**); for the initial version of *R2*, *L_HS_* = 1026 bp, corresponding to *π_i_* = 2.52 × 10^-5^ (used for the binomial test in **Figure S1H**); for improved *R2*, *L_HS_*= 1488 bp, corresponding to *π_i_* = 3.66 × 10^-5^ (used for the binomial test in **Figures 1I, S1N, S3J** and **S4I**).

During the recombination step, the length of the homologous sequence in the SuRe-CC system is 900 bp, corresponding to *π_i_* = 2.21 × 10^-5^ in females (used for the binomial test in **Figures 1K, 2F, S1Q** and **S2D**). The sequences of *attP* and *attB* sites are not the same. Thus, we did not use the length of *attP* or *attB* to calculate the natural recombination rate. However, the serine recombinase catalyzes crossover of *attP* and *attB* at the central dinucleotide^35–37,104^. Therefore, for the SuRe-CR system, we set *L_HS_* = 2 bp, corresponding to *π_i_* = 4.92 × 10^-8^ in females (used for the binomial test in **Figure 4D,E**).

We used a similar approach to compare the recombination efficiency of SuRe to the natural recombination in *C. elegans*. Here, the recombination frequency is 1.32 cM/Mbp at the transgene insertion site (V:8,644,770) for male and hermaphrodite meiosis^105^. The length of the homologous sequence is *L_HS_* = 500 bp, corresponding to *π_i_* = 0.66 × 10^-5^ (used for the binomial test in **Figure 3I**).

We compared measured fidelity values with the value expected under the hypothesis of a uniform distribution, which assumes that all possible recombination products occur with equal probability. We calculated the expected fidelity *π_i_* for both adaptor insertion and recombination steps of the SuRe system. During the adaptor insertion step, if an IGA has *n* transgenes, the expected fidelity of adaptor insertion is 1/*n* (**Figure S3D–G**). During the recombination step, when recombining two IGAs containing *n_U_* and *n_D_* transgenes respectively, the expected fidelity of recombination is 1/(*n_U_n_D_*) (**Figures 2A** and **S3A–C**). We used the binomial test to compare the measured and expected fidelity values (**Figures 2E, 4F, S3I** and **S4H**).

To compare the efficiency of individual flies in multiple groups, we used the non-parametric Kruskal-Wallis one-way ANOVA followed by Tukey’s honestly significant difference (HSD) test. Statistical analyses were performed using the Kruskal-Wallis test with multiple comparisons corrections using the functions *kruskalwallis()* and *multcompare()* in MATLAB (version R2022a). To compare the data in two groups, we performed a non-parametric Wilcoxon rank sum test using the MATLAB function *ranksum()*.

### Mathematical modeling of transgene assembly using SuRe and convention methods

To evaluate the scalability of transgene assembly with SuRe and conventional methods, we assessed two key metrics: assembly time and workload, both as a function of the number of transgenes. These metrics respectively correspond to the time and spatial complexity metrics used in computer science.

With sequential assembly, assembly time increases linearly with the number of transgenes (**Figure 7A,B**). In contrast, with binary assembly the assembly time rises logarithmically with transgene number (**Figure 7C,D**). Traditional genetic methods and the SuRe system enable binary assembly, significantly reducing assembly time for large numbers of transgenes as compared to sequential integration by genome editing (**Figure 7E**).

When one uses natural meiotic recombination methods or the SuRe system to recombine multiple transgenes, one must choose the correct recombination product by genotyping. There are two possible approaches to the genotyping, either genotype individual genes by PCR or sequence the entire genome of the strain. If there are more than hundreds of target genes in a strain to be typed, whole genome sequencing is less expensive than PCR. If PCR is used for genotyping, the cost rises linearly with the number of genes. With whole genome sequencing, the cost rises linearly with the number of strains examined. We evaluate these two numbers to compare natural meiotic recombination to the SuRe system.

In each recombination step, if the probability of getting one desired strain by genotyping is *p* and one seeks *r* independent strains, the number of genes to be genotyped, *X*, follows a negative binomial distribution:

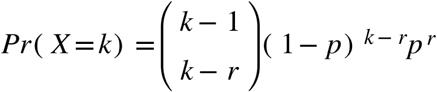

The expectation value of *X* is *r*/*p*, implying that, on average, we need to genotype *r*/*p* strains to obtain *r* independent desired strains. Alternatively, if we use natural meiotic recombination approaches for recombination, we need to genotype all the transgenes in the desired strain. The number of genes to be genotyped at this recombination step is the *r*/*p* times the number of transgenes in the desired strain. If we use the SuRe system for recombination, we only need to genotype the genes adjacent to the adaptors. The number of genes to be genotyped at the recombination step is the *r*/*p* times 2. The number of genes to be genotyped at the adaptor insertion step is the *r*/*p*. We used the expectation value of the total number of strains or genes in all the steps

to represent the total genotyping workload. Notably, we are interested in how the total genotyping workload grows with the total number of transgenes (*N*). We derived the asymptotic number of strains or genes needed to be genotyped in the regime that *N* is large. The **Supplementary Appendix** has the details of this derivation.

