## Supplemental Appendix for "Super Recombinator (SuRe): An *in vivo* recombination system for scalable and efficient transgene assembly at a single genomic locus"

### Supplementary Appendix

#### §1. Overview.

This appendix evaluates the workload involved in recombining multiple transgenes using traditional genetic methods or the SuRe system. Genotyping, the process of identifying correctly recombined strains, is the most time-intensive task after the transgenic strains with single transgenes are made. In **§2**, we establish two criteria for comparison: the total number of genes and strains that require genotyping, which we evaluate quantitatively in the subsequent sections.

Recombination processes typically fall into two categories: sequential recombination (introducing one transgene per step, with $N$ steps for $N$ transgenes; **Figure 7A,B**) and binary recombination (simultaneously combining transgenes from a pair of strains, reducing the number of steps to $\log_2 N$; **Figure 7C,D**).

In **§3** and **§4**, we calculate the genotyping workload for traditional genetics approaches using sequential (**§3**) and binary (**§4**) recombination processes. In **§5** and **§6**, we compute the genotyping workload for the SuRe-CC (**§5**) and SuRe-CR (**§6**) systems.

Our comparisons between sequential and binary recombination reveal that, regardless of whether traditional genetic approaches or the SuRe system are used, binary recombination speeds the recombination process and reduces the genotyping workload. Further comparisons between traditional genetic approaches and the SuRe systems (SuRe-CC and SuRe-CR) show that the SuRe-CR system, particularly when combined with binary recombination, requires the least genotyping workload and thus has the best time efficiency. **Supplemental Table 1** summarizes the relevant equations.

#### §2. Metrics of genotyping workload.

This section focuses on estimating the genotyping workload required for transgene recombination, a crucial aspect of comparisons between SuRe and traditional genetic methods. First, we derive a formula for the number of strains that need to be evaluated to identify the desired recombinant in each step, based on the probability of obtaining this recombinant. We discuss two main genotyping techniques, Polymerase Chain Reaction (PCR) and Whole-Genome Sequencing (WGS), highlighting their respective cost implications, particularly when there are a large number of target genes. Finally, we define key metrics, the total numbers of strains and genes that must be genotyped, to quantify the overall workload and facilitate selection of a cost-effective strategy.

To estimate the genotyping workload, we first consider the number of strains needed to identify the desired recombinant in each step. Let $P$ be the probability that any given strain is the desired recombination product, assuming independence between strains. We examine two distinct genotyping methods: sequential and parallel genotyping.

**Sequential Genotyping:** If strains are genotyped one at a time until $r$ desired recombinants are found, the number of genotyped strains ($X$) follows a negative binomial distribution:

$$\Pr\left(X=k\right)=\binom{k-1}{k-r}\left(1-P\right)^{k-r}P^{r} \tag{2.1}$$

It is easy to derive that the expected value of $X$ is:

$$\mathrm{E}\left(X\right)=r/P\,. \tag{2.2}$$

**Parallel Genotyping:** Alternatively, if $r/P$ strains are genotyped in parallel, the number of desired strains within this batch ($Y$) follows a binomial distribution:

$$\Pr\left(Y=k\right)=\binom{r/P}{k}\left(1-P\right)^{r/P-k}P^{k}\quad. \tag{2.3}$$

The distribution of $Y$ has an expectation value of $r$ desired strains ($\mathrm{E}\left(Y\right)=r$). The probability of obtaining at least one desired recombinant in this batch is:

$$\Pr\left(\text{Getting desired strains in } r/P \text{ strains}\right) = 1-\left(1-P\right)^{r/P} > 1-e^{-r} \quad . \tag{2.4}$$

Typically, a redundancy factor of $r$ = 3 is used. For parallel genotyping, this ensures a probability >95% of obtaining at least one desired recombinant per batch for any value of $P$.

In summary, for both sequential and parallel genotyping, we can use $r/P$ to evaluate the number of strains that require genotyping in each step. When using traditional genetic approaches to recombine transgenes that share the same markers (*e.g., mini-w*$^{+}$), $P$ equals the proportion of desired strains in the progeny. In this case, the value of $P$ is determined by the genomic locations of the transgenes, as detailed in **§3** and **§4**. With the SuRe system, fluorescent markers on the adaptors allow for eliminating unrecombined strains, simplifying genotyping. In this case, $P$ equals the fidelity of recombination and adaptor insertion. We have experimentally measured the fidelity of recombination and adaptor insertion (**Figures 2E**, **4F**, **S3H**, **S4H**). Detailed methods for estimating the fidelity of recombination and adaptor insertion across varying transgene numbers are described in **§5** and **§6**.

To genotype transgenic strains, we consider two primary techniques: polymerase chain reaction (PCR) and whole-genome sequencing (WGS). PCR is a targeted approach and detects the presence of a single gene of interest. Conversely, whole-genome sequencing (WGS) provides a comprehensive view, identifying all target genes within a strain in a single assay. The cost of PCR scales linearly with the number of genes to be analyzed per strain. WGS, however, offers a constant cost per strain, irrespective of the number of genes interrogated. Although individual PCR assays are less expensive than WGS, for applications requiring genotyping of numerous genes (hundreds or more) within each strain, WGS becomes a more cost-effective alternative. Based on these cost considerations, we designed three genotyping strategies for transgene assembly.

The first strategy employs WGS for all genotyping steps. In this approach, the cost per step is directly proportional to the number of strains genotyped at that step. Let $P_i$ be the probability that any given strain is the desired recombination product in step *i*. The total workload, characterized by the total number of strains that require genotyping ($S_{total}$), can be expressed as:

$$S_{total} = r\sum_i \frac{1}{P_i} \quad . \tag{2.5}$$

The second strategy uses PCR for all genotyping. Here, the cost per step scales with both the number of strains genotyped and the number of genes analyzed per strain. Let $P_i$ be the probability that any given strain is the desired recombination product and $g_i$ be the number of genes genotyped per strain in step *i*. The total workload, characterized by the total number of genes that require genotyping ($G_{total}$), is given by:

$$G_{total} = r\sum_{i}\frac{g_i}{P_i} \quad . \tag{2.6}$$

The third strategy is a hybrid approach, combining the strengths of both WGS and PCR. This strategy aims to enhance efficiency by initially using PCR to genotype for a select few genes, thereby eliminating a significant portion of undesired strains early in the process. Subsequently, WGS is applied to the remaining candidate strains to definitively confirm the presence of all other target genes. This combined approach offers a substantial advantage over purely PCR- or WGS-based strategies, particularly in using traditional genetic approaches to recombine transgenes on a single chromosome. In such scenarios, PCR-based genotyping of just two genes can effectively filter out most non-recombinant strains. Consequently, the overall genotyping workload for this hybrid strategy is primarily dependent on the total number of strains ($S_{total}$). We will delve into the detailed implementation and advantages of this strategy in **§3** and **§4**.

In summary, to evaluate the genotyping workload across the three strategies, we primarily rely on two key metrics: $S_{total}$ and $G_{total}$. These criteria allow fair comparisons between traditional genetic approaches and the SuRe system. Although mathematical expressions for $S_{total}$ and $G_{total}$ can be complex, we apply two asymptotic estimators to simplify estimations of their values as the number of transgenes to be assembled (*N*) rises. These asymptotic estimators obey the following:

$$\lim_{N\to+\infty} S_{total}/S_{asymptotic} = 1 \quad , \tag{2.7}$$

$$\lim_{N\to+\infty} G_{total}/G_{asymptotic} = 1 \quad . \tag{2.8}$$

#### §3. Sequential recombination with traditional genetics approaches.

This section considers the sequential recombination process with traditional genetics approaches. We calculate the numbers of strains and genes that must be genotyped under different scenarios, including those in which transgenes lie on either separate chromosomes or on a single chromosome. If the transgenes are on a single chromosome, we consider cases in which they are either randomly or evenly spaced, as well as cases in which the insertion sites are optimized to minimize the genotyping workload. Mathematical formulae and asymptotic expressions are derived below to estimate the numbers of strains and genes needed, using recombination probabilities based on chromosome distances and crossover rates.

To recombine *N* transgenes, the total number of steps needed is $N-1$. The number of genes that must be genotyped per strain in step *i* is:

$$g_i = i+1 \quad . \tag{3.1}$$

If all *N* transgenes are located on separate chromosomes, the probability of obtaining the desired strain in step *i* is:

$$P_i = 2^{-i-1} \quad . \tag{3.2}$$

By substituting Eqs. (3.1) and (3.2) into Eqs. (2.5) and (2.6), we can derive the number of strains or genes that require genotyping:

$$S_{total} = r\sum_{i=1}^{N-1} 2^{i+1} = r\left(2^{N+1} - 4\right) \quad , \tag{3.3}$$

$$G_{total} = r\sum_{i=1}^{N-1} \left(i+1\right) 2^{i+1} = r 2^{N+1}\left(N-1\right) \quad . \tag{3.4}$$

However, if all $N$ transgenes are on one chromosome, to calculate the recombination probability we need to consider how these genes are distributed on the chromosome. To facilitate this calculation, we describe the length of the chromosome, $D$, in centimorgan (cM) units. A 1 cM distance means that the expected average number of intervening chromosomal crossovers in a single generation is 0.01. Two transgenes are recombined if there are an odd number of chromosomal crossovers between them. If the distance between two transgenes is $d$ cM, the probability for them to recombine is:

$$\Pr\left(\text{Recombination}\right) = \frac{1 - e^{-2d/100}}{2} \quad . \tag{3.5}$$

In female Drosophila Melanogaster, 1 cM corresponds to ~2 Mbp[1]. The total lengths of chromosomes X, II, III, and IV are about 66 cM, 112 cM, 99 cM, and 0 cM[1].

The total numbers of strains and genes required for genotyping depend on the locations of the $N$ transgenes on the chromosome. If 2 transgenes are located randomly on one chromosome whose total length is $D$, the distribution of their distance follows the probability density function:

$$f_{D_1}\left(d_1\right) = \frac{2\left(D - d_1\right)}{D^2} \quad . \tag{3.6}$$

The expectation values of the net numbers of strains and genes that must be genotyped are:

$$S_{total} = \frac{1}{2} G_{total} = r\int_0^D \frac{2}{1 - e^{-2d_1/100}} \frac{2\left(D - d_1\right)}{D^2} \partial d_1 = +\infty \quad . \tag{3.7}$$

The net numbers of strains and genes to be genotyped when $N \geq 2$ are not less than those when $N = 2$ case, irrespective of whether recombination is sequential or binary. Therefore, if the genes are randomly inserted into one chromosome, the asymptotic expression of $S_{total}$ and $G_{total}$ is:

$$S_{asymptotic} = +\infty \quad , \tag{3.8}$$

$$G_{asymptotic} = +\infty \quad . \tag{3.9}$$

If $N$ transgenes are located evenly on the chromosome, the distance between a pair of adjacent transgenes is $D/(N-1)$. In step $i$, the desired strain must retain the $i$ transgenes recombined in prior steps and introduce transgenes $i$+1. The probability of obtaining the desired strain in this step is:

$$P_i = (1-p)^{i-1}\frac{p}{2} \quad , \tag{3.10}$$

Here $p = \frac{1-e^{-2(D/100)/(N-1)}}{2}$ is the probability of the recombination between two adjacent transgenes.

By substituting Eqs. (3.1) and (3.10) into Eqs. (2.5) and (2.6), we can derive the number of strains or genes that require genotyping:

$$S_{total} = r\frac{2(1-p)}{p^2}\left[(1-p)^{-N+1}-1\right] \quad , \tag{3.11}$$

$$G_{total} = r\frac{2(1-p)}{p^3}\left[1-2p+(1-p)^{-N+1}\left(p(N+1)-1\right)\right] \quad . \tag{3.12}$$

Substituting the expression of $p$ into the above two equations will complicate these two equations. Here, we derive the asymptotic expression of $S_{total}$ and $G_{total}$ when $N$ is large:

$$S_{asymptotic} = r\frac{2\left(e^{D/100}-1\right)}{\left(D/100\right)^2}N^2 \quad , \tag{3.13}$$

$$G_{asymptotic} = r\frac{2\left[1+\left(D/100-1\right)e^{D/100}\right]}{\left(D/100\right)^3}N^3 \quad . \tag{3.14}$$

We note that if we fine-tune the insertion sites of the transgenes, it is possible to reduce the total number of strains or genes that require genotyping. Let $d_i$ to be the distance between transgene $i$ and $i$ + 1. To minimize the total number of strains or genes that require genotyping, we can minimize the following two functions, using the vector, $\vec{d}$, to represent ($d_1, d_2, \ldots d_{N-1}$):

$$S_{total}\left(\vec{d}\right) = r\sum_{i=1}^{N-1} 2\left[\frac{1-e^{-2d_i/100}}{2}\prod_{j=1}^{i-1}\left(1-\frac{1-e^{-2d_j/100}}{2}\right)\right]^{-1} \quad , \tag{3.15}$$

$$G_{total}\left(\vec{d}\right) = r\sum_{i=1}^{N-1}(i+1)2\left[\frac{1-e^{-2d_i/100}}{2}\prod_{j=1}^{i-1}\left(1-\frac{1-e^{-2d_j/100}}{2}\right)\right]^{-1} \quad , \tag{3.16}$$

under the constraint:

$$\sum_{i=1}^{N-1} d_i = D \quad . \tag{3.17}$$

It is hard to derive the closed forms of the minimum values of $S_{total}\left(\vec{d}\right)$ and $G_{total}\left(\vec{d}\right)$. However, we can derive asymptotic forms for their minimum values and corresponding $\vec{d}$ when $N$ is large. Here, we define a function characterizing the distribution of the transgene as:

$$f_{d_i}\left(x\right) = \lim_{N\to\infty} Nd_{xN}/100 \quad . \tag{3.18}$$

From Eq. (3.17), we know that the function above follows the constraint:

$$\int_0^1 f_{d_i}\left(x\right)dx = D/100 \quad . \tag{3.19}$$

Then, Eqs. (3.15) and (3.16) can be converted into the following two functionals of $f_{d_i}\left(x\right)$:

$$\lim_{N\to\infty} \frac{S_{total}\left(\vec{d}\right)}{rN^2} = \int_0^1 \frac{2}{f_{d_i}\left(x\right)} \exp\left(\int_0^x f_{d_i}\left(y\right)dy\right)dx \quad , \tag{3.20}$$

$$\lim_{N\to\infty} \frac{G_{total}\left(\vec{d}\right)}{rN^3} = \int_0^1 x\frac{2}{f_{d_i}\left(x\right)} \exp\left(\int_0^x f_{d_i}\left(y\right)dy\right)dx \quad . \tag{3.21}$$

If $f_{d_i}\left(x\right) = D/100$, it implies the transgenes are located evenly on the chromosome. By substituting this expression into Eqs. (3.20) and (3.21), we reach the same asymptotic estimation Eqs. (3.13) and (3.14).

We can use the Euler-Lagrange equation from the calculus of variations to derive the optimal $f_{d_i}\left(x\right)$ for $S_{total}\left(\vec{d}\right)$ and $G_{total}\left(\vec{d}\right)$. For $S_{total}\left(\vec{d}\right)$, the optimal $f_{d_i}\left(x\right)$ follows the differential equation:

$$\frac{2}{f_{d_i}\left(x\right)} \exp\left(\int_0^x f_{d_i}\left(y\right)dy\right) - \frac{\partial}{\partial x}\left(\frac{-2}{f_{d_i}^2\left(x\right)} \exp\left(\int_0^x f_{d_i}\left(y\right)dy\right)\right) = 0 \quad . \tag{3.22}$$

The differential equation above can be simplified into:

$$\frac{\partial f_{d_i}\left(x\right)}{\partial x} = f_{d_i}^2\left(x\right) \quad . \tag{3.23}$$

From the differential equation (3.23) and the constraint in Eq. (3.19), we can derive the closed form of optimal $f_{d_i}\left(x\right)$ for $S_{total}\left(\vec{d}\right)$:

$$f_{d_i}\left(x\right)=\left[\left(1-e^{-D/100}\right)^{-1}-x\right]^{-1} \quad . \tag{3.24}$$

By substituting the optimal $f_{d_i}\left(x\right)$ into Eq. (3.20), we obtain the corresponding minimum value of $S_{total}\left(\vec{d}\right)$:

$$S_{asymptotic}=r\frac{2}{1-e^{-D/100}}N^2 \quad . \tag{3.25}$$

With the same approach, we can derive the optimal $f_{d_i}\left(x\right)$ for $S_{gene}\left(\vec{d}\right)$ using

$$x\frac{2}{f_{d_i}\left(x\right)}\exp\left(\int_0^x f_{d_i}\left(y\right)dy\right)-\frac{\partial}{\partial x}\left(x\frac{-2}{f_{d_i}^2\left(x\right)}\exp\left(\int_0^x f_{d_i}\left(y\right)dy\right)\right)=0 \quad . \tag{3.26}$$

This differential equation can be simplified into:

$$\frac{\partial f_{d_i}\left(x\right)}{\partial x}=f_{d_i}^2\left(x\right)+\frac{1}{2x}f_{d_i}\left(x\right) \quad . \tag{3.27}$$

From (3.27) and the constraint in (3.19), we derive the closed form of optimal $f_{d_i}\left(x\right)$ for $S_{gene}\left(\vec{d}\right)$:

$$f_{d_i}\left(x\right)=\frac{3}{2}x^{1/2}\left[\left(1-e^{-D/100}\right)^{-1}-x^{3/2}\right]^{-1}. \tag{3.28}$$

By substituting the optimal $f_{d_i}\left(x\right)$ into (3.20), we obtain the corresponding minimum of $G_{total}\left(\vec{d}\right)$:

$$G_{asymptotic}=r\frac{8/9}{1-e^{-D/100}}N^3 \quad . \tag{3.29}$$

**§4. Binary recombination with traditional genetics approaches.**

This section details the genotyping workload associated with using binary recombination with traditional genetic approaches. It calculates the number of strains and genes needed for genotyping when combining transgenes via a binary process, which reduces the number of steps to the logarithm of the transgene number. We consider various scenarios, with the transgenes on separate or a single chromosome. For transgenes on a single chromosome, we consider both randomly and evenly spaced transgene locations, as well as optimized insertion sites. Mathematical formulas and asymptotic expressions are derived to estimate the number of strains and genes required, based on the recombination probabilities and the symmetry of the binary recombination process.

To recombine $N$ transgenes, the total number of steps is $n=\lceil\log_2 N\rceil$. The symbol $\lceil\ \rceil$ here represents the ceiling function.

If $\log_2 N$ is an integer, the recombination process is highly symmetric. In step *i*, we create $N2^{-i}$ strains, each of which contains $2^i$ transgenes. If $\log_2 N$ is not an integer, the recombination process is more complex, but we can still use the symmetry to simplify the derivation. In step *i*, we create two types of strains, which have $g_{i,1} = \lfloor N2^{-n+i} \rfloor$ and $g_{i,2} = \lceil N2^{-n+i} \rceil$ transgenes. The symbol $\lfloor \ \rfloor$ here represents the floor function. When $g_{i,1} = g_{i,2}$, without loss of generality, we consider all the strains to be type 2 strains at step *i*.

We define the recombination coefficient $r_{i,j,j'}$ as how many type *j* strains at step *i* – 1 are used to create one type *j* strain at step *i*. If $g_{i,1}$ is even, $g_{i,1} = \lfloor N2^{-n+i} \rfloor = 2\lfloor N2^{-n+i-1} \rfloor = 2g_{i-1,1}$. That means the type 1 strain at step *i* can be made from a pair of type 1 strains at step *i* – 1. As this example shows, if $g_{i,1}$ is even, $r_{i,1,1} = 2, r_{i,1,2} = 0$. Similarly, if $g_{i,2}$ is even, $r_{i,1,1} = 0, r_{i,1,2} = 2$; if $g_{i,j}$ is odd, $r_{i,j,1} = 1, r_{i,j,2} = 1$. In summary, we can obtain the value of the recombination coefficient matrix $R_i$ in the four cases in which both $g_{i,1}$ and $g_{i,2}$ are even or odd:

$$R_i = \begin{pmatrix} r_{i,1,1} & r_{i,1,2} \\ r_{i,2,1} & r_{i,2,2} \end{pmatrix} = \begin{cases} \begin{pmatrix} 2 & 0 \\ 0 & 2 \end{pmatrix} & g_{i,1} \text{ and } g_{i,2} \text{ are even.} \\ \begin{pmatrix} 1 & 1 \\ 1 & 1 \end{pmatrix} & g_{i,1} \text{ and } g_{i,2} \text{ are odd.} \\ \begin{pmatrix} 2 & 0 \\ 1 & 1 \end{pmatrix} & g_{i,1} \text{ is even; } g_{i,2} \text{ is odd.} \\ \begin{pmatrix} 1 & 1 \\ 0 & 2 \end{pmatrix} & g_{i,1} \text{ is odd; } g_{i,2} \text{ is even.} \end{cases} \tag{3.30}$$

This expression of $R_i$ can be further simplified into:

$$R_i = 2\begin{pmatrix} 1 & 0 \\ 0 & 1 \end{pmatrix} + \begin{pmatrix} g_{i,1} - 2g_{i-1,1} \\ g_{i,2} - 2g_{i-1,2} \end{pmatrix} \begin{pmatrix} -1 & 1 \end{pmatrix} \ . \tag{3.31}$$

Because each strain at step *i* is made from two strains from step *i* – 1, and $g_{i,j} = \sum_{j'=1}^{2} r_{i,j,j'} g_{i-1,j'}$. It is straightforward to confirm that $R_i$ follows the two equations:

$$R_i \begin{pmatrix} 1 & g_{i-1,1} \\ 1 & g_{i-1,2} \end{pmatrix} = \begin{pmatrix} 2 & g_{i,1} \\ 2 & g_{i,2} \end{pmatrix} \ . \tag{3.32}$$

To create the type $j$ strain at step $i$, we need to recombine the transgenes in the two strains from the previous step, and the transgenes from these two strains do not segregate. Therefore, the probability of getting the desired type $j$ strain at step $i$ is:

$$P_{i,j} = \frac{1}{2} p_{i,j} Q_{i-1,1}^{r_{i,j,1}} Q_{i-1,2}^{r_{i,j,2}}; \left(P_{1,j} = p_{1,j}\right) \quad . \tag{3.33}$$

Here, $p_{i,j}$ is the probability of recombination between the two strains. $Q_{i,j}$ is the probability that the transgenes on the type $j$ strain at step $i$ do not segregate in the next step. Then, the probability $Q_{i,j}$ follows the recursive equation:

$$Q_{i,j} = \left(1 - p_{i,j}\right) Q_{i-1,1}^{r_{i,j,1}} Q_{i-1,2}^{r_{i,j,2}}; \left(Q_{1,j} = 1 - p_{1,j}\right) \quad . \tag{3.34}$$

From this recursive equation, we know $Q_{i,j}$ can be represented as the product of $\left(1 - p_{i',j'}\right)$ raised to the power of $m_{i,i',j,j'}$

$$Q_{i,j} = \prod_{i'=1}^{i} \prod_{j'=1}^{2} \left(1 - p_{i',j'}\right)^{m_{i,i',j,j'}} \quad . \tag{3.35}$$

Here, we note that if $\log_2 N$ is not an integer, $g_{1,1} = \left\lfloor N 2^{-n+1} \right\rfloor = 1$. Then, there is only one transgene on the strain corresponding to $p_{1,1}$. In this case, we can let $p_{1,1} = 0$ to satisfy Eq. (3.35). However, when we estimate the number of strains and genes to genotype ($S_{strain}$ and $S_{gene}$), we should eliminate the term $\frac{2}{p_{1,1}}$, because the type 1 strain with only one transgene does not need to be genotyped in the combination step 1. We will discuss this again later when deriving expressions for $S_{strain}$ and $S_{gene}$. By substituting Eq. (3.35) into the recursive equation (3.34), we derive a recursive equation for $m_{i,i',j,j'}$ :

$$M_{i,i'} = \begin{pmatrix} m_{i,i',1,1} & m_{i,i',1,2} \\ m_{i,i',2,1} & m_{i,i',2,2} \end{pmatrix} = R_i \begin{pmatrix} m_{i-1,i',1,1} & m_{i-1,i',1,2} \\ m_{i-1,i',2,1} & m_{i-1,i',2,2} \end{pmatrix} = R_i M_{i-1,i'}; \left( M_{i,i} = \begin{pmatrix} 1 & 0 \\ 0 & 1 \end{pmatrix} \right). \tag{3.36}$$

When $i > i'$, we derive the equation:

$$M_{i,i'} = R_i R_{i-1} \ldots R_{i'+1} = \prod_{j=0}^{i-i'-1} R_{i-j} \quad . \tag{3.37}$$

Using Eq. (3.32), we can simplify Eq. (3.37) into:

$$M_{i,i'}\begin{pmatrix}1 & g_{i',1}\\ 1 & g_{i',2}\end{pmatrix}=\begin{pmatrix}2^{i-i'} & g_{i,1}\\ 2^{i-i'} & g_{i,2}\end{pmatrix} \quad . \tag{3.38}$$

Finally, we can solve for the closed form of $M_{i,i'}$ :

$$M_{i,i'}=\begin{cases}\begin{pmatrix}0 & 0\\ 0 & 0\end{pmatrix} & i<i'\\ 2^{i-i'}\begin{pmatrix}1 & 0\\ 0 & 1\end{pmatrix}+\begin{pmatrix}g_{i,1}-g_{i',1}2^{i-i'}\\ g_{i,2}-g_{i',2}2^{i-i'}\end{pmatrix}\begin{pmatrix}-1 & 1\end{pmatrix} & i\geq i'\end{cases} \quad . \tag{3.39}$$

Here, we should notice that if $g_{i'1}=g_{i',2}$, the matrix $\begin{pmatrix}1 & g_{i',1}\\ 1 & g_{i',2}\end{pmatrix}$ is irreversible. But we can confirm that Eq. (3.39) satisfies Eqs. (3.36) and (3.37), regardless if $g_{i'1}=g_{i',2}$ or $g_{i'1}\neq g_{i',2}$.

Finally, we need to obtain one type 2 strain at step *n*. We define $s_{i,j}$ as the number of type *j* strains at step *i*. Because the total number of strains at step *i* is $2^{n-i}$, and the total number of genes at step *i* is *N*, the expression for $s_{i,j}$ satisfies the following,

$$\begin{cases}s_{i,1}+s_{i,2}=2^{n-i}\\ g_{i,1}s_{i,1}+g_{i,2}s_{i,2}=N\end{cases} \quad . \tag{3.40}$$

It is straightforward to confirm that $s_{i,j}=m_{n,i,2,j}$ is the solution of the above equation by substituting the expression of $m_{n,i,2,j}$ in Eq. (3.39) into Eq. (3.40).

From Eqs. (3.33), (3.34), (3.35) and (3.39), we can simplify the expression of $P_{i,j}$ into:

$$P_{i,j}=\frac{1}{2}p_{i,j}\prod_{i'=1}^{i-1}\prod_{j'=1}^{2}\left(1-p_{i',j'}\right)^{m_{i,i',j,j'}} \quad . \tag{3.41}$$

By substituting Eq. (3.41) into Eqs. (2.5) and (2.6), we obtain the expression of the total number of strains ($S_{total}$) or genes ($G_{total}$) to genotype:

$$S_{total}=r\left[m_{n,1,2,2}\frac{2}{p_{1,2}}+\sum_{i=2}^{n}\sum_{j=1}^{2}m_{n,i,2,j}\frac{2}{p_{i,j}}\left(1-p_{1,2}\right)^{-m_{i,1,j,2}}\prod_{i'=2}^{i-1}\prod_{j'=1}^{2}\left(1-p_{i',j'}\right)^{-m_{i,i',j,j'}}\right] \quad . \tag{3.42}$$

$$G_{total}=r\left[g_{1,2}m_{n,1,2,2}\frac{2}{p_{1,2}}+\sum_{i=2}^{n}\sum_{j=1}^{2}g_{i,j}m_{n,i,2,j}\frac{2}{p_{i,j}}\left(1-p_{1,2}\right)^{-m_{i,1,j,2}}\prod_{i'=2}^{i-1}\prod_{j'=1}^{2}\left(1-p_{i',j'}\right)^{-m_{i,i',j,j'}}\right] \quad . \tag{3.43}$$

In these two equations, we eliminated terms containing $p_{1,1}$ by considering the following two cases. For the first case, $\log_2 N$ is not an integer, $g_{1,1} = \lfloor N2^{-n+1} \rfloor = 1$. We do not need to genotype for the type 1 strain at step 1. Then, we can eliminate the term $m_{n,1,2,1}\frac{2}{p_{1,1}}$ in $S_{total}$ and the term $g_{1,1}m_{n,1,2,1}\frac{2}{p_{1,1}}$ in $G_{total}$ to avoid $p_{1,1} = 0$ making these terms infinite. In this case, because $p_{1,1} = 0$, we can eliminate the term $\left(1-p_{1,1}\right)^{m_{i,1,j,1}}$ in the expression of $P_{i,j}$ to get $P_{i,j} = \frac{1}{2}p_{i,j}\left(1-p_{1,2}\right)^{m_{i,1,j,2}}\prod_{i'=2}^{i-1}\prod_{j'=1}^{2}\left(1-p_{i',j'}\right)^{m_{i,i',j,j'}}$. For the second case, $\log_2 N$ is an integer, $\lfloor N2^{-n+i} \rfloor = \lceil N2^{-n+i} \rceil$, $m_{i,i',2,1} = 0$. We can still eliminate the term $m_{n,1,2,1}\frac{2}{p_{1,1}}$ in $S_{total}$ and the term $g_{1,1}m_{n,1,2,1}\frac{2}{p_{1,1}}$ in $G_{total}$ because these terms are 0. For the same reason, we can eliminate all the terms corresponding to $j = 1$. To satisfy the expressions for $S_{total}$ and $G_{total}$ in this case, we only need to check the expression of $P_{i,j}$ in the terms with $j = 2$. Because $m_{i,1,2,1} = 0$, we can eliminate the term $\left(1-p_{1,1}\right)^{m_{i,1,2,1}}$ in the expression of $P_{i,2}$ to get $P_{i,2} = \frac{1}{2}p_{i,2}\left(1-p_{1,2}\right)^{m_{i,1,2,2}}\prod_{i'=2}^{i-1}\prod_{j'=1}^{2}\left(1-p_{i',j'}\right)^{m_{i,i',2,j'}}$. In summary, Eqs. (3.42) and (3.43) are true whether or not $\log_2 N$ is an integer. We use these two equations to calculate the total number of strains and genes that require genotyping. These two equations made full use of the symmetry of binary recombination, even if $\log_2 N$ is not an integer. We need to consider no more than $2\lceil \log_2 N \rceil - 1$ variables $p_{i,j}$ in the calculation.

If all $N$ transgenes are located on separate chromosomes, these transgenes independently assort at each step. Thus, it is straightforward to see that the probability $p_{i,j} = 1/2$. From Eqs. (3.39), (3.42) and (3.43), we derive the number of strains or genes that require genotyping:

$$S_{total} = r\left[4\left(N-2^{n-1}\right) + \sum_{i=2}^{n} 2^{n-i+g_{i,2}} - \left(g_{i,2}2^{n-i} - N\right)\left(2^{g_{i,2}} - 2^{g_{i,1}}\right)\right] . \tag{3.44}$$

$$G_{total} = r\left[8\left(N-2^{n-1}\right) + \sum_{i=2}^{n} g_{i,2}2^{n-i+g_{i,2}} - \left(g_{i,2}2^{n-i} - N\right)\left(g_{i,2}2^{g_{i,2}} - g_{i,1}2^{g_{i,1}}\right)\right] . \tag{3.45}$$

The last term of $S_{total}$, in which $i = n$, equals $2^N$, and other terms are non-negative but not more than $N2^{N/2}$. Hence, $r2^N \le S_{total} \le r\left[\left(n-1\right)N2^{N/2} + 2^N\right]$. Similarly, the last term of $G_{total}$, in which $i = n$, equals $N2^N$, and the other terms are non-negative but not more than $2N2^{N/2}$, implying $rN2^N \le G_{total} \le r\left[\left(n-1\right)2N2^{N/2}+N2^N\right]$. By the Squeeze Theorem, we can derive asymptotic estimators for $S_{total}$ and $G_{total}$ when *N* is large:

$$S_{asymptotic}=r2^N \quad , \tag{3.46}$$

$$G_{asymptotic}=rN2^N \quad . \tag{3.47}$$

These two equations indicate that when *N* is large, the workload of the last step is dramatically larger than the sum of all previous steps.

If all the *N* transgenes are evenly spaced on one chromosome, as mentioned in **§3**, recombining 2 genes requires genotyping an infinite set of strains and genes. The asymptotic expressions for $S_{total}$ and $G_{total}$ are:

$$S_{asymptotic}=+\infty \quad , \tag{3.48}$$

$$G_{asymptotic}=+\infty \quad . \tag{3.49}$$

If all *N* transgenes are evenly located on one chromosome, the distance between two adjacent genes is $D/\left(N-1\right)$. Thus, it is straightforward to see that $p_{i,j}=p=\frac{1-e^{-2(D/100)/(N-1)}}{2}$. From Eqs. (3.39), (3.42) and (3.43), we derive the number of strains or genes that require genotyping:

$$S_{total}=r\frac{2}{p}\left[\left(N-2^{n-1}\right)+\sum_{i=2}^{n}\frac{2^{n-i}}{\left(1-p\right)^{g_{i,2}-2}}-\left(g_{i,2}2^{n-i}-N\right)\left(\frac{1}{\left(1-p\right)^{g_{i,2}-2}}-\frac{1}{\left(1-p\right)^{g_{i,1}-2}}\right)\right] \quad , \tag{3.50}$$

$$G_{total}=r\frac{2}{p}\left[2\left(N-2^{n-1}\right)+\sum_{i=2}^{n}\frac{2^{n-i}g_{i,2}}{\left(1-p\right)^{g_{i,2}-2}}-\left(g_{i,2}2^{n-i}-N\right)\left(\frac{g_{i,2}}{\left(1-p\right)^{g_{i,2}-2}}-\frac{g_{i,1}}{\left(1-p\right)^{g_{i,1}-2}}\right)\right] \quad . \tag{3.51}$$

We use these two equations to compute exact values of $S_{total}$ and $G_{total}$. We can also derive asymptotic estimators for $S_{total}$ and $G_{total}$ when *N* is large. Because $m_{n,i,2,1}=g_{i,2}2^{n-i}-N\ge 0$ and $m_{n,i,2,2}=2^{n-i}-\left(g_{i,2}2^{n-i}-N\right)\ge 0$, we can obtain lower bounds for $S_{total}$ and $G_{total}$ :

$$S_{total}>r\frac{2}{p}\left[\left(N-2^{n-1}\right)+\sum_{i=2}^{n}2^{n-i}\right]=r\frac{2}{p}\left(N-1\right) \quad , \tag{3.52}$$

$$G_{total} > r\frac{2}{p}\left[\left(N-2^{n-1}\right)+\sum_{i=2}^{n}2^{n-i}\left(N2^{-n+i}-1\right)\right]=r\frac{2}{p}\left[N-2^{n}+1+N\left(n-1\right)\right] \quad . \tag{3.53}$$

Because $m_{n,i,2,1}=g_{i,2}2^{n-i}-N\geq 0$ and $N2^{-n+i}-1<g_{i,1}\leq g_{i,2}<N2^{-n+i}+1$, we can obtain the upper bound of $S_{total}$ and $G_{total}$:

$$S_{total} < r\frac{2}{p}\left[\left(N-2^{n-1}\right)+\sum_{i=2}^{n}2^{n-i}\left(1-p\right)^{-N2^{-n+i}+1}\right] \quad . \tag{3.54}$$

$$G_{total} < r\frac{2}{p}\left[\left(N-2^{n-1}\right)+\sum_{i=2}^{n}2^{n-i}\left(N2^{-n+i}+1\right)\left(1-p\right)^{-N2^{-n+i}+1}\right] \quad . \tag{3.55}$$

It is clear that $\lim_{N\to+\infty}Np=D/100$. Here, we need to find the limit of $\sum_{i=2}^{n}2^{n-i}\left(1-p\right)^{-N2^{-n+i}}$ and $\sum_{i=2}^{n}\left(1-p\right)^{-N2^{-n+i}}$ to derive the limit of the upper bounds of $S_{total}$ and $G_{total}$. If $A>0$ and $\lim_{N\to+\infty}B\left(N\right)=+\infty$, we can prove the following limit by L'Hôpital's rule:

$$\lim_{N\to+\infty}\frac{\int_{b}^{B(N)}e^{A/x}dx}{B\left(N\right)}=\lim_{N\to+\infty}\frac{e^{A/B(N)}B'\left(N\right)}{B'\left(N\right)}=\lim_{N\to+\infty}e^{A/B(N)}=1 \quad . \tag{3.56}$$

Because the function $e^{\left[-N\ln(1-p)\right]/x}$ monotonically decreases with *x*, we can further prove the following two inequalities:

$$\sum_{i=2}^{n}2^{n-i}\left(1-p\right)^{-N2^{-n+i}}=2\sum_{i=0}^{n-2}\left(2^{i}-2^{i-1}\right)\left(1-p\right)^{-N2^{-i}}<2\int_{1/2}^{2^{n-2}}e^{\left[-N\ln(1-p)\right]/x}dx=2^{n-1}+o\left(N\right) \quad ,\tag{3.57}$$

$$\begin{aligned}\sum_{i=2}^{n}\left(1-p\right)^{-N2^{-n+i}}&=\sum_{i=0}^{n-2}\left[i-\left(i-1\right)\right]\left(1-p\right)^{-N2^{-i}}\\&<\left(1-p\right)^{-N}+\left(1-p\right)^{-N/2}+\int_{1}^{n-2}e^{\left[-N\ln(1-p)\right]/x}dx=\log_2 N+o\left(\log_2 N\right)\end{aligned} \quad . \tag{3.58}$$

These equations allow us to derive asymptotic estimators of $S_{total}$ and $G_{total}$:

$$S_{asymptotic}=r\frac{2}{D/100}N^{2} \quad , \tag{3.59}$$

$$G_{asymptotic}=r\frac{2}{D/100}N^{2}\log_2 N \quad . \tag{3.60}$$

To derive the equation for optional $S_{total}$ and $G_{total}$, we can optimize Eqs. (3.42) and (3.43) under the constraint of the total length of the chromosome. To generate type *i* strain at step *j*, we need the recombination to happen at the chromosome region with length $d_{i,j}$, and the transgenes at other chromosome regions do not segregate. Thus, the chromosome region with the length $d_{i,j}$ corresponding to the recombination probability $p_{i,j}$ which follows the equation:

$$p_{i,j} = \frac{1-e^{-2\left(d_{i,j}/100\right)}}{2} \quad . \tag{3.61}$$

$D_{i,j}$ is defined as the distance between the first and last transgene in this strain. Thus, $D_{i,j}$ follows the recursive equation:

$$D_{i,j} = d_{i,j} + r_{i,j,1}D_{i-1,1} + r_{i,j,2}D_{i-1,2};\left(D_{1,j} = d_{1,j}\right) \quad . \tag{3.62}$$

Here, we note that if $g_{1,1} = 1$, there is only one transgene on the strain corresponding to $d_{1,1}$. In this case, we can let $d_{1,1} = 0$ without losing generality. Eq. (3.61) still holds true.

From Eq. (3.62), we know $D_{i,j}$ is a linear combination of $d_{i',j'}$. Comparing Eq. (3.62) with Eq. (3.34), we can derive the expression of $D_{i,j}$ and $Q_{i,j}$ from the recursive equations (3.62):

$$D_{i,j} = \sum_{i'=1}^{i}\sum_{j'=1}^{2} m_{i,i',j,j'}d_{i',j'} \quad . \tag{3.63}$$

The final product of the recombination is the type 2 strain at step *n*. Therefore, the total distance obeys the constraint $D \geq D_{n,2}$. Similar to the elimination of the term containing $p_{1,1}$ in Eqs. (3.42) and (3.43), we can eliminate the term containing $d_{1,1}$ in the expression of $D_{n,2}$. Thus, the constraint of $d_{i,j}$ satisfies:

$$D \geq m_{n,1,2,2}d_{1,2} + \sum_{i=2}^{n}\sum_{j=1}^{2} m_{n,i,2,j}d_{i,j} \quad . \tag{3.64}$$

Eqs. (3.42), (3.43), and (3.64) can be used to compute the optimal values of $S_{total}$ and $G_{total}$ numerically. In the following derivation, we will obtain asymptotic estimators of the optimal $S_{total}$ and $G_{total}$. Eq. (3.61) indicated that $d_{i,j}/100 \geq p_{i,j}$. Combine this inequality with Eq. (3.64), we can further derive the constraint:

$$\frac{D}{100} > m_{n,1,2,2}p_{1,2} + \sum_{i=2}^{n}\sum_{j=1}^{2} m_{n,i,2,j}p_{i,j} \quad . \tag{3.65}$$

From Eqs. (3.42) and (3.43), we derive lower bounds on the optimal values of $S_{total}$ and $G_{total}$ using the Cauchy-Schwarz inequality:

$$\begin{aligned}\frac{D}{100}\frac{S_{total}}{r} &> \left(m_{n,1,2,2}p_{1,2}+\sum_{i=2}^{n}\sum_{j=1}^{2}m_{n,i,2,j}p_{i,j}\right)\left[m_{n,1,2,2}\frac{2}{p_{1,2}}+\sum_{i=2}^{n}\sum_{j=1}^{2}m_{n,i,2,j}\frac{2}{p_{i,j}}\right] \\ &\geq \left[m_{n,1,2,2}\sqrt{2}+\sum_{i=2}^{n}\sum_{j=1}^{2}m_{n,i,2,j}\sqrt{2}\right]^2 = 2(N-1)^2\end{aligned} \quad . \quad (3.66)$$

$$\begin{aligned}\frac{D}{100}\frac{G_{total}}{r} &> \left(m_{n,1,2,2}p_{1,2}+\sum_{i=2}^{n}\sum_{j=1}^{2}m_{n,i,2,j}p_{i,j}\right)\left[g_{1,2}m_{n,1,2,2}\frac{2}{p_{1,2}}+\sum_{i=2}^{n}\sum_{j=1}^{2}g_{i,j}m_{n,i,2,j}\frac{2}{p_{i,j}}\right] \\ &\geq \left[m_{n,1,2,2}\sqrt{2g_{1,2}}+\sum_{i=2}^{n}\sum_{j=1}^{2}m_{n,i,2,j}\sqrt{2g_{i,j}}\right]^2 = 2\left[m_{n,1,2,2}g_{1,2}^{1/2}+\sum_{i=2}^{n}\sum_{j=1}^{2}m_{n,i,2,j}g_{i,j}^{1/2}\right]^2\end{aligned} \quad . \quad (3.67)$$

These inequalities suggest that the optimal $p_{i,j}$ approximate the following equation for large *N*,

$$p_{i,j} \approx \frac{D/100}{2^n \bar{g}_{n,2}(a)} g_{i,j}^a e^{(D/100)2^{i-n-1}} \quad . \quad (3.68)$$

Here, we define a set of functions $\bar{g}_{i,j}(a)$ as:

$$\bar{g}_{i,j}(a) = 2^{-i}\left(m_{i,1,j,2}g_{1,2}^a e^{(D/100)2^{-n}} + \sum_{i'=2}^{i}\sum_{j'=1}^{2}m_{i,i',j,j'}g_{i',j'}^a e^{(D/100)2^{i'-n-1}}\right) \quad . \quad (3.69)$$

The normalization factor $C_a = \frac{D/100}{2^n \bar{g}_{n,2}(a)}$ in Eq. (3.68) makes $p_{i,j}$ approximately match the constraint in Eq. (3.65).

We can substitute Eq. (3.68) into Eqs. (3.42) and (3.43) to estimate the upper bounds of the optimal $S_{total}$ and $G_{total}$. Eqs. (3.42) and (3.43) contain the term:

$$\left(1-p_{1,2}\right)^{-m_{i,1,j,2}}\prod_{i'=2}^{i-1}\prod_{j'=1}^{2}\left(1-p_{i',j'}\right)^{-m_{i,i',j,j'}} < e^{-m_{i,1,j,2}\ln\left(1-p_{1,2}\right)-\sum_{i'=2}^{i}\sum_{j'=1}^{2}m_{i,i',j,j'}\ln\left(1-p_{i',j'}\right)} = e^{C_a 2^i \bar{g}_{i,j}(a)\left[1+o(1)\right]} \,. \quad (3.70)$$

We need to estimate the upper bound of this term to estimate the upper bounds of the optimal $S_{total}$ and $G_{total}$. From this definition equation (3.69) and Eqs. (3.31) and (3.36), we can prove $\bar{g}_{i,j}(a)$ increases with *i* and *j* for any $a \geq 0$. Therefore, $\bar{g}_{i,j}(a) \leq \bar{g}_{n,2}(a)$. This allows us to estimate the upper bound of $\left(1-p_{1,2}\right)^{-m_{i,1,j,2}}\prod_{i'=2}^{i-1}\prod_{j'=1}^{2}\left(1-p_{i',j'}\right)^{-m_{i,i',j,j'}}$ :

$$\left(1-p_{1,2}\right)^{-m_{i,1,j,2}}\prod_{i'=2}^{i-1}\prod_{j'=1}^{2}\left(1-p_{i',j'}\right)^{-m_{i,i',j,j'}}<e^{(D/100)\frac{2^{i}\bar{g}_{i,j}(a)}{2^{n}\bar{g}_{n,2}(a)}\left[1+o(1)\right]}\le e^{(D/100)2^{i-n}\left[1+o(1)\right]}\quad. \tag{3.71}$$

Interestingly, this upper bound does not contain the parameter *a*.

By substituting $a=0$, Eqs. (3.68), and (3.71) into Eq. (3.42), we can prove:

$$\frac{S_{total}}{r}<\frac{2}{C_0}\left[m_{n,1,2,2}e^{-(D/100)2^{-n}}+\sum_{i=2}^{n}\sum_{j=1}^{2}m_{n,i,2,j}e^{(D/100)2^{i-n-1}\left[1+o(1)\right]}\right]<\frac{2^{1+2n}\bar{g}_{n,2}^{2}\left(0\right)}{D/100}\left[1+o\left(1\right)\right]\ . \tag{3.72}$$

By substituting $a=1/2$, Eqs. (3.68) and (3.71) into Eq. (3.43), we can prove:

$$\frac{G_{total}}{r}<\frac{2}{C_{1/2}}\left[m_{n,1,2,2}g_{1,2}^{1/2}e^{-(D/100)2^{-n}}+\sum_{i=2}^{n}\sum_{j=1}^{2}m_{n,i,2,j}g_{i,j}^{1/2}e^{(D/100)2^{i-n-1}\left[1+o(1)\right]}\right]<\frac{2^{1+2n}\bar{g}_{n,2}^{2}\left(1/2\right)}{D/100}\left[1+o\left(1\right)\right] \tag{3.73}$$

Based on Eq. (3.56), we can prove:

$$\begin{aligned}2^{n}\bar{g}_{n,2}\left(0\right)&=\left(N-2^{n-1}\right)e^{(D/100)2^{-n}}+\sum_{i=2}^{n}\left(2^{n-i+1}-2^{n-i}\right)e^{(D/100)2^{i-n-1}}\\&<\left(N-2^{n-1}\right)e^{(D/100)2^{-n}}+\int_{1}^{2^{n-1}}e^{(D/100)/x}dx=N+o\left(N\right)\end{aligned}\quad. \tag{3.74}$$

Combining Eqs. (3.66), (3.72), and (3.74), we can derive:

$$S_{asymptotic}=r\frac{2}{D/100}N^{2}\quad. \tag{3.75}$$

The $p_{i,j}$ corresponding to the $S_{asymptotic}$ can be estimated via the equation:

$$p_{i,j}\approx\frac{D/100}{2^{n}\bar{g}_{n,2}\left(0\right)}e^{(D/100)2^{i-n-1}}\quad. \tag{3.76}$$

Based on the inequality $\sum_{j=1}^{2}m_{n,i,2,j}g_{i,j}^{1/2}\le\sqrt{\left(\sum_{j=1}^{2}m_{n,i,2,j}\right)\left(\sum_{j=1}^{2}m_{n,i,2,j}g_{i,j}\right)}=\sqrt{2^{n-i}N}$, we can prove:

$$\begin{aligned}&m_{n,1,2,2}g_{1,2}^{1/2}+\sum_{i=2}^{n}\sum_{j=1}^{2}m_{n,i,2,j}g_{i,j}^{1/2}\le\left(N-2^{n-1}\right)2^{1/2}+\sum_{i=2}^{n}\sqrt{2^{n-i}N}\\&=\left[2^{1/2}-2^{-1/2}\left(2^{n}/N\right)+\sqrt{2^{n}/N}\left(1+2^{-1/2}\right)\right]N-o\left(N\right)\\&=\left[\sqrt{2}+1+2^{-1/2}\left(\sqrt{2}-\sqrt{2^{n}/N}\right)\left(\sqrt{2^{n}/N}-1\right)\right]N-o\left(N\right)\end{aligned}\quad, \tag{3.77}$$

$$\begin{aligned}
&2^n \bar{g}_{n,2}(1/2) - \left( m_{n,1,2,2} g_{1,2}^{1/2} + \sum_{i=2}^{n} \sum_{j=1}^{2} m_{n,i,2,j} g_{i,j}^{1/2} \right) \\
&\le \left( N - 2^{n-1} \right) 2^{1/2} \left[ e^{(D/100)2^{-n}} - 1 \right] + \sum_{i=2}^{n} \left[ e^{(D/100)2^{i-n-1}} - 1 \right] \sqrt{2^{n-i} N} \\
&= \left[ \sum_{j=1}^{+\infty} \frac{(D/200)^j}{j!} \frac{1 - 2^{(-j+1/2)(n-1)}}{1 - 2^{-j+1/2}} \right] \sqrt{N} + o\left(\sqrt{N}\right) < \frac{e^{D/200} - 1}{1 - 2^{-1/2}} \sqrt{N} + o\left(\sqrt{N}\right)
\end{aligned} \quad . \tag{3.78}$$

Based on the inequality $g_{i,j}^{1/2} \ge g_{1,2}^{1/2} = \sqrt{2}$, we can prove:

$$m_{n,1,2,2} g_{1,2}^{1/2} + \sum_{i=2}^{n} \sum_{j=1}^{2} m_{n,i,2,j} g_{i,j}^{1/2} > \left( N - 2^{n-1} \right) \sqrt{2} + \sum_{i=2}^{n} \sum_{j=1}^{2} m_{n,i,2,j} \sqrt{2} = (N-1)\sqrt{2} \quad . \tag{3.79}$$

Combining Eqs. (3.77), (3.78) and (3.79), we can prove:

$$\lim_{N \to +\infty} \frac{2^n \bar{g}_{n,2}(1/2)}{m_{n,1,2,2} g_{1,2}^{1/2} + \sum_{i=2}^{n} \sum_{j=1}^{2} m_{n,i,2,j} g_{i,j}^{1/2}} = 1 \quad . \tag{3.80}$$

Combining Eqs. (3.67), (3.73) and (3.80), we can derive:

$$G_{asymptotic} = r \frac{2^{1+2n} \bar{g}_{n,2}^2 (1/2)}{D/100} \quad . \tag{3.81}$$

When $\log_2 N$ is an integer, $g_{i,j} = 2^i$. The equation above can be simplified into

$$G_{asymptotic} \approx r \frac{2\left(1+\sqrt{2}\right)^2}{D/100} N^2 \quad . \tag{3.82}$$

Eqs. (3.77) and (3.81) indicate:

$$G_{asymptotic} \le r \frac{2}{D/100} \left[ \sqrt{2} + 1 + 2^{-1/2} \left( \sqrt{2} - \sqrt{2^n/N} \right) \left( \sqrt{2^n/N} - 1 \right) \right]^2 N^2 \quad . \tag{3.83}$$

Because $1 \le 2^n/N < 2$, the factor $\left[ \sqrt{2} + 1 + 2^{-1/2} \left( \sqrt{2} - \sqrt{2^n/N} \right) \left( \sqrt{2^n/N} - 1 \right) \right]^2$ is in the range $\left[ \left(1+\sqrt{2}\right)^2, \left( 1 + \sqrt{2} + 2^{-1/2} \left( \frac{\sqrt{2}-1}{2} \right)^2 \right)^2 \right] \approx [5.83, 5.98)$. When $\log_2 N$ is an integer, this factor equals $\left(1+\sqrt{2}\right)^2$. When $\log_2 N$ is not an integer, if we use the simplified Eq. (3.82) for estimation, the error is < 2.53%.

The $p_{i,j}$ corresponding to the $G_{asymptotic}$ can be estimated using the equation:

$$p_{i,j} \approx \frac{D/100}{2^n \bar{g}_{n,2}(1/2)} g_{i,j}^{1/2} e^{(D/100)2^{i-n-1}} \quad . \tag{3.84}$$

**§5. Recombination with SuRe-CC.**

Here we estimate the number of strains or genes that must be genotyped when we use the SuRe-CC system. These numbers include the strains or genes genotyped in the recombination and adaptor insertion steps. Interestingly, the numbers of strains or genes that must be genotyped in the recombination steps and adaptor insertion steps are unrelated to whether recombination is sequential, binary, or another irregular process. Below, we first discuss genotyping for the recombination steps and next for the adaptor insertion steps.

As noted in **§2**, the probability of obtaining the desired recombination products at each recombination step equals the fidelity of the SuRe-CC system. The number of strains or genes we need to genotype is inversely proportional to the fidelity of recombination. When using SuRe-CC for recombination, the homologous sequences in the residual adaptor may also serve as the homologous arm in the recombination. To recombine two transgenic tandems with $n_U$ and $n_D$ transgenes, there are $n_U n_D$ possible recombination products (**Figures 2A,B** and **S3A–C**). Our experimental results suggest that the progeny of female F3 exhibit these recombination products with equal probability (**Figure 2E**). The results also suggest that the fidelity of the male F3 is about two-fold that of the female F3 (**Figure 2E**). Hence, the fidelity of the SuRe-CC system can be described by the following equation:

$$P_{RS,CC} = \begin{cases} 1/(n_U n_D) & Female \\ \min\left(2/(n_U n_D),1\right) & Male \end{cases} \quad . \tag{4.1}$$

Here, RS is the abbreviation of the recombination step. The expression, $\min\left(2/(n_U n_D),1\right)$, implies that the fidelity of male F3 is not larger than 1 when $n_U = 1, n_D = 1$. In other words, when $n_U = 1, n_D = 1$, the fidelity of male F3 is 1; otherwise, the fidelity of the male F3 is $2/(n_U n_D)$.

We noted that it is not necessary to type all the transgenes in the transgenic tandem to identify the desired recombination products. To pick the desired recombination product, we only need to confirm the existence of the two genes adjacent to the two adaptors (for example, genes B and C in **Figure 2A,B**). Thus, the relationship between the total numbers of strains and genes is:

$$G_{RS,total} = 2S_{RS,total} \quad . \tag{4.2}$$

To estimate the numbers of strains and genes to genotype, we compute the sum of all the recombination steps. We use the plots in **Figure S7D,E** to compute this sum and show that it is not related to the recombination process. In these plots, the coordinate (*i*, *j*) represents a transgenic tandem containing genes *i* to *j*. Recombining this tandem with another one containing genes *j* + 1 to *k* produces a new transgenic tandem containing gene *i* to *k* (**Figure S7D**). This recombination step is represented by the rectangle with vertices (*i*, *j*), (*j* + 1, *k*), (*i*, *k*), and (*j* + 1, *j*). The width and height of the rectangle are (*j* + 1 – *i*) and (*k* – *j*), corresponding to the numbers of genes in the two original transgenic tandems ($n_U$ and $n_D$). The area of this rectangle is $\left(j+1-i\right)\left(k-j\right)=n_U n_D$ (**Figure S7D**). It is proportional to the numbers of strains and genes that require genotyping (Eqs. (4.1) and (4.2)). The initial transgenes are denoted by the coordinates (*i*, *i*) {*i* = 1, 2, … *N*} at the diagonal of the plot. The final recombination product is represented by the coordinate (1, *N*) at the upper left corner of each plot. Regardless of the recombination process, the rectangles representing each recombination step collectively fill the upper triangular area of each plot (**Figure S7E**). The total area of these rectangles is $N\left(N-1\right)/2$. As shown in Eq. (4.1), when $n_U=1, n_D=1$, the fidelity of the male F3 is 1 instead of $2/\left(n_U n_D\right)$. However, the number of rectangles with area 1 (the rectangles with pink dots in **Figure S7E**) does not exceed $N/2$. This effect becomes negligible as *N* rises. Thus, asymptotic values of $S_{RS,total}$ and $G_{RS,total}$ for the SuRe-CC system are:

$$S_{RS,asymptotic}=r\frac{1}{2a_{RS,CC}}N^2 \quad , \tag{4.3}$$

$$G_{RS,asymptotic}=r\frac{1}{a_{RS,CC}}N^2 \quad . \tag{4.4}$$

Here, $a_{RS,CC}$ is the fidelity factor of the recombination step of the SuRe-CC system. It equals 1 for females and 2 for males (**Figure 2E**).

The cause of incorrect adaptor insertion differs from that of incorrect recombination. Adaptor insertion also needs the homologous arm flanking the adaptor sequence to induce the adaptor insertion, and there are $n_T$ copies of the same homologous sequence in a transgenic tandem with $n_T$ transgenes. However, the desired and undesired products do not appear with equal probability. When the desired product is created, both ends of the double-strand break generated by Cas9/gRNA-d align perfectly with the template DNA (**Figure S3D**). However, in generating the undesired product, only one end of the double-strand break achieves perfect alignment (**Figure S3E–G**). This inherent asymmetry in the repair process creates a bias favoring the formation of the desired product during adaptor insertion. Here, we assume the probability of creating desired or undesired products declines exponentially with the length of the unaligned DNA. Namely, the proportion of the $k^{\text{th}}$ undesired product is $a_{AIS}^k$ times the proportion of the desired product, where $a_{AIS}$ is a constant describing the exponential decline of this proportion. Thus, the fidelity of the adaptor insertion is:

$$P_{AIS} = \frac{1}{\sum_{k=0}^{n_T-1} a_{AIS}^k} = \frac{1-a_{AIS}}{1-a_{AIS}^{n_T}} \quad . \tag{4.5}$$

Our experiments showed that $P_{AIS} \approx 0.90$ and $a_{AIS} \approx 0.10$ when $n_T = 4$ (**Figure S3**). This implies $a_{AIS}^{n_T} << 1$. To simplify the derivation, we approximate the fidelity of the adapter insertion to be $P_{AIS} \approx 0.90$ for any $n_T$.

We only need to type the transgene adjacent to the adaptor to confirm that the adaptor is inserted correctly. Thus, the relationship between the total numbers of strains and genes is:

$$G_{AIS,total} = S_{AIS,total} \quad . \tag{4.6}$$

For any recombination process, no matter sequential, binary, or another irregular recombination process, there are $2(N-1)$ adaptor insertion steps to recombine $N$ transgenes. When $N$ is large, the asymptotic values of $S_{AIS,total}$ and $G_{AIS,total}$ are:

$$S_{AIS,asymptotic} = G_{AIS,asymptotic} = r\frac{2}{P_{AIS}}N \quad . \tag{4.7}$$

Because the SuRe-CC and SuRe-CR systems use the same mechanism for adaptor insertion, Eq. (4.7) applies to both systems.

Comparing Eqs. (4.3), (4.4), and (4.7), we found that the genotyping workload in the adaptor insertion steps is much less than that in the recombination steps. Therefore, we can derive the asymptotic values of $S_{total}$ and $G_{total}$ for the SuRe-CC system:

$$S_{asymptotic} = r\frac{1}{2a_{RS,CC}}N^2 \quad , \tag{4.8}$$

$$G_{asymptotic} = r\frac{1}{a_{RS,CC}}N^2 \quad . \tag{4.9}$$

##### §6. Recombination with SuRe-CR.

In this section, we estimate the number of strains or genes that must be genotyped when we use the SuRe-CR system. As with SuRe-CC, these numbers include the strains or genes that must be genotyped in the recombination and adaptor insertion steps. Both parts are independent of the recombination process.

No matter whether recombination is sequential, binary, or another irregular recombination process, there are $N-1$ reactions to recombine $N$ transgenes. Because Serine recombinase only catalyzes the recombination between attP and attB sites, the attR sites in the adaptor residuals do not recombine with attP or attR sites in adaptors. So, unlike the SuRe-CC system, the fidelity of the SuRe-CR system is not related to the number of transgenes. In our experiments, we observed that the fidelity of the three serine recombinases was close to but not equal to 100%. Thus, asymptotic values of $S_{RS,total}$ and $G_{RS,total}$ for the SuRe-CR system are:

$$S_{RS,asymptotic} = r\frac{1}{P_{RS,CR}}N \quad , \tag{5.1}$$

$$G_{RS,asymptotic} = r\frac{2}{P_{RS,CR}}N \quad . \tag{5.2}$$

Here, $P_{RS,CR}$ is the fidelity of the recombination step of the SuRe-CR system. It is approximately 98% for PhiC31, 99% for Bxb1, and 87% for TP901-1 (**Figure 4F**).

Since the adaptor insertion steps of the SuRe-CR system are similar to those of SuRe-CC, the asymptotic values of $S_{AIS,total}$ and $G_{AIS,total}$ for the SuRe-CR system also follow Eq. (4.7). Therefore, we can derive asymptotic values of $S_{total}$ and $G_{total}$ for the SuRe-CR system:

$$S_{asymptotic} = r\left(\frac{1}{P_{RS,CR}} + \frac{2}{P_{AIS}}\right)N \quad , \tag{5.3}$$

$$G_{asymptotic} = r\left(\frac{2}{P_{RS,CR}} + \frac{2}{P_{AIS}}\right)N \quad . \tag{5.4}$$

#### §7. Comparing the genotyping workload among different recombination approaches.

**Supplemental Table 1** compares the number of strains or genes required for genotyping using different recombination approaches. Among all the methods, binary recombination requires less genotyping workload than sequential recombination. Additionally, the SuRe system offers a significant reduction in genotyping workload compared to traditional approaches. Finally, the SuRe-CR system requires the lowest genotyping workload for genotyping among all the methods compared.

#### §8. Supplementary Table

**Supplementary Table 1 | Numbers of strains or genes required for genotyping in the limit of large *N*.**

| | | | | Number of strains to genotype | | Number of genes to PCR | |
|---|---|---|---|---|---|---|---|
| Traditional genetic approach | Separate chromosomes | Sequential | | $r2^{N+1}$ | (3.3) | $r2^{N+1}N$ | (3.4) |
| | | Binary | | $r2^{N}$ | (3.46) | $r2^{N}N$ | (3.47) |
| | Single chromosome* | Sequential | Random | $+\infty$ | (3.8) | $+\infty$ | (3.9) |
| | | | Even | $r\frac{2\left(e^{D/100}-1\right)}{\left(D/100\right)^2}N^2$ | (3.13) | $r\frac{2\left[1+\left(D/100-1\right)e^{D/100}\right]}{\left(D/100\right)^3}N^3$ | (3.14) |
| | | | Optimized | $r\frac{2}{1-e^{-D/100}}N^2$ | (3.25) | $r\frac{8/9}{1-e^{-D/100}}N^3$ | (3.29) |
| | | Binary | Random | $+\infty$ | (3.48) | $+\infty$ | (3.49) |
| | | | Even | $r\frac{2}{D/100}N^2$ | (3.59) | $r\frac{2}{D/100}N^2\log_2 N$ | (3.60) |
| | | | Optimized | $r\frac{2}{D/100}N^2$ | (3.75) | $r\frac{\left(2+\sqrt{2}\right)^2}{D/100}N^2$ ** | (3.82) |
| SuRe | The SuRe-CC system# | | | $r\frac{1}{2a_{RS,CC}}N^2$ | (4.8) | $r\frac{1}{a_{RS,CC}}N^2$ | (4.9) |
| | The SuRe-CR system## | | | $r\left(\frac{1}{P_{RS,CR}}+\frac{2}{P_{AIS}}\right)N$ | (5.3) | $r\left(\frac{2}{P_{RS,CR}}+\frac{2}{P_{AIS}}\right)N$ | (5.4) |

*: $D$ is the length of the chromosome in unit centimorgan (cM).

**: This equation estimates $G_{asymptotic}$ when $\log_2 N$ is an integer. If $\log_2 N$ is not an integer, the error of this equation is < 2.53% (Eq. (3.83)).

#: $a_{RS,CC}$ is the fidelity factor of the recombination step of the SuRe-CC system. It equals 1 for females and 2 for males (**Figure 2E**).

##: $P_{RS,CR}$ is the fidelity of the recombination step of the SuRe-CR system. It approximates 98% for PhiC31, 99% for Bxb1, and 87% for TP901-1 (**Figure 4F**). $P_{AIS}$ is the fidelity of the adaptor insertion step. It approximates 90% for both SuRe-CC and SuRe-CR systems (**Figures S3I, S4H**).

#### §9. References.

1. Comeron JM, Ratnappan R, Bailin S. The Many Landscapes of Recombination in Drosophila melanogaster. PLoS Genetics 2012 Oct 11;8(10):e1002905. Available from: http://dx.doi.org/10.1371/journal.pgen.1002905.
