## Supplemental Tables for "Super Recombinator (SuRe): An *in vivo* recombination system for scalable and efficient transgene assembly at a single genomic locus"

**Table S1 | gRNAs used in this research.**

| | gRNA name | gRNA target | Description |
|---|---|---|---|
| ***D. melano-gaster*** | gRNAu | ACCGTCGACGATGTAGGTCANGG | AD1 insertion site, upstream of transgene, targeting the vector backbone |
| | gRNAd | CCGCAGTCCGATCATCGGATNGG | AD2 insertion site, downstream of transgene, targeting the mini-white marker in the vector |
| | gRNAd(old) | TGTCGGCTACTCCTTGCGTCNGG | The old version of *AD2* insertion site, downstream of transgene, targeting the mini-white marker in the vector |
| | gRNA1 | GTCCTTCAGGTCGCCTCCGGNGG | gRNA in *AD2* of SuRe-CC, targeting *AD1* to mediate recombination |
| | gRNA2 | GGAGGAAGACGAGCGGTCCANGG | gRNA in *AD1* of SuRe-CC, targeting *AD2* to mediate recombination |
| ***C. elegans*** | oxTi365_sgRNA | AACAAGTTGGGACAATACGGNGG | Chr V single-copy insertion site |
| | AD1_sgRNA | TACACTAAACTACTAAAGTANGG | AD1 insertion site, 482 bp upstream of oxTi365_sgRNA |
| | AD2_sgRNA | AAATGAAATCTAACTAGCAGNGG | AD2 insertion site, 532 bp downstream of oxTi365_sgRNA |
| | dpy10_sgRNA | GCTACCATAGGCACCACGAGNGG | dpy-10 co-CRISPR |
| | reco_sgRNA | GTCCTTCAGGTCGCCTCCGGNGG | gRNA to mediate recombination |

**Table S2 | Primers used in this research.**

| | Primer names | Primer sequences | Primer locations | Products to genotype | Figures |
|---|---|---|---|---|---|
| *D. melanogaster* | HS1f | CTCCACCTTCCGCTTTTTCTTGGG | Edge of AD1 | AD1 insertion | 1F, 6A |
| | JT1r | TCTTGAAGACGAAAGGGCCTCGTG | Edge of the transgene | AD1 insertion | 1F, 6A |
| | GL2f | CGTTTAGATCCACTAGTTCTAGAGCGG | Edge of the Janelia GAL4 or LexA transgene | AD2 insertion to Janelia GAL4 or LexA | 1F |
| | UL2f | GGCATGCCCTAGGGCTAGAGTC | Edge of the UAS or LexAop transgene | AD2 insertion to UAS or LexAop | 6A |
| | HS2r | TTCTTGCCATACCATTTAGCCGATC | Edge of AD2 | AD2 insertion | 1F, 6A |
| | MWf | GGGATGTTCGAAGGGGAAATACTTG | Flanking the adaptor residual | Recombination | 1F, 5F, 6A |
| | VBr | GGGTTTCGCCACCTCTGACTTG | Flanking the adaptor residual | Recombination | 1F, 5F, 6A |
| | PE3f | TATCGCTGTCTCACTCAGACTCAATACGAC | 3' end of the P-element vector | The upstream edge of TG1 and the downstream edge of TG2 | 5F |
| | PE5r | CGACGCATTTCGTACTCCAAAGTACG | 5' end of the P-element vector | The upstream edge of AD2 and the downstream edge of AD1 | 5F |
| | Dp5r | ACGGAGAGTACGTGGTAAACAACATCCATC | Upstream of TG1 insertion site | The upstream edge of TG1 | 5F |
| | G3f | CTTCTCGCAGTCCCACAAA | Downstream of TG1 insertion site | The downstream edge of TG1 | 5F |
| | G5r (Tfb5[k10127]) | TTTGATGTTGCCGGCATCTTTAGCCATGAC | Upstream of TG2 (Tfb5[k10127]) insertion site | The upstream edge of TG2 (Tfb5[k10127]) | 5F |
| | Dp3f (Tfb5[k10127]) | CAGCCTGAAGACCGGCAAGGAGAAGAAAGG | Downstream of TG2 (Tfb5[k10127]) insertion site | The downstream edge of TG2 (Tfb5[k10127]) | 5F |
| | G5r (FASN2[k05816]) | GGTTCGCTTTCGCGATCATCTTTCAGAGCG | Upstream of TG2 (FASN2[k05816]) insertion site | The upstream edge of TG2 (FASN2[k05816]) | 5F |
| | Dp3f (FASN2[k05816]) | CAGCATCCGCATGCAGGGGTCCATCAGCTC | Downstream of TG2 (FASN2[k05816]) insertion site | The downstream edge of TG2 (FASN2[k05816]) | 5F |
| *C. elegans* | 1F | GGGCTCTCCCAGTTCGTTCTTCTC | Upstream of AD1 insertion site | AD1 insertion | 3F |
| | 1R | GGACATTTTTCTTTGGCCATATAGGAGG | In the promoter of the TG | AD1 insertion | 3F |
| | 2F | GGCGAACGATGAGAATTGTCCTCGC | In the UTR of the TG | AD2 insertion | 3F |
| | 2R | GGGTACAGAATCTACCTGCAACAGTTC | Downstream of AD1 insertion site | AD2 insertion | 3F |
| | inF | GGAGCATCCAGCTTTCAAATCTTTGTG | Flanking the adaptor residual | Recombination | 3F |
| | inR | GCATAACTTTTTTATGGGAATTTTACCTACAAGAC | Flanking the adaptor residual | Recombination | 3F |
| | oxTi365_F | TGAACATCAGTAGATACCTAGAAGTACACT | Upstream of TG insertion site | TG insertion | 3F |
| | oxTi365_R | CAAACTATCCAGTGCTTTTTTATGGTGTCAAC | Downstream of TG insertion site | TG insertion | 3F |

**Table S3 | *C. elegans* strains used in this research.**

| Strain Name | *C. elegans* Genotype | Description |
|---|---|---|
| TV26083 | *wySi916 V* | wySi916 = pdes2::ANv3-TSER (PVD/FLP voltage sensor. Not Visible) |
| TV26857 | *wySi925 wySi916 V* | wySi925 = AD1 cassette w/ pric-19::GFP (pan-neuronal selection marker) with wySi916 |
| TV26696 | *wySi916 wySi917 V* | wySi917 = AD2 cassette w/ punc-122::GFP (coelomocyte selection marker) with wySi916 |
| TV26999 | *wySi929 V* | wySi929 = recombination of wySi916 to create 2x tandem transgene (PVD/FLP voltage sensor, 2x. Not Visible) |
| TV26733 | *wySi919 V* | wySi919 = pdes-2::myr-mScarlet::let-858 3'UTR (PVD/FLP mScarlet) |
| TV26954 | *wySi926 wySi919 V* | wySi926 = AD1 cassette w/ pric-19::GFP (pan-neuronal selection marker) with wySi919 |
| TV26955 | *wySi919 wySi927 V* | wySi927 = AD2 cassette w/ punc-122::GFP (coelomocyte selection marker) with wySi919 |
| TV29645 | *wySi1021[wySi919 wySi919] V* | wySi1021 = recombination of wySi919 to create 2x tandem transgene (PVD/FLP mScarlet, 2x) |
| TV27388 | *wySi926 wySi951 V* | wySi951 = prab-3::myr-mScarlet::let-858 3'UTR (pan-neuronal mScarlet) with AD1 |
| TV27387 | *wySi955 wySi927 V* | wySi955 = pdes-2::myrBFP::let-858 3'UTR (PVD/FLP BFP) with AD2 |
| TV27805 | *wySi973[wySi951 wySi955] V* | wySi973 = recombination of wySi951 (pan-neuronal mScarlet) and wySi955 (PVD/FLP BFP) |
| TV27128 | *wySi937 wySi927 V* | wySi937 = pdes-2::FLPase::let-858 3'UTR (PVD/FLP FLPase) with AD2 |
| TV29681 | *wySi1027[wySi919 wySi937] V* | wySi1027 = recombination of wySi919 V (PVD/FLP mScarlet) & wySi937 V (PVD/FLP FLPase) |
| TV27075 | *wySi926 wySi934 V* | wySi934 = pser-2::myr-mScarlet::let-858 3' UTR with AD1 |
| TV29644 | *wySi966 wySi927 V* | wySi966 = pdes-2::myr-mNeonGreen::let-858 3'UTR with AD2 |
| TV27582 | *wySi965 [wySi934, wySi966] V* | wySi965 = recombination of wySi934 (PVD-only mScarlet) and wySi966 (PVD/FLP mNeonGreen) |

**Table S4 | *D. melanogaster* strains used in this research.**

| Stock # | *D. melanogaster* Genotype |
|---|---|
| 605091 | y[1] M{vas-NLS-TP901-1.nanos3'UTR.GFP-.RFP-}ZH-2A w[*]; MKRS/TM6B, Tb[1] |
| 605092 | y[1] w[*]; P{y[+t7.7] w[+mC]=13XlexAop-pAce.x2}attP40; MKRS/TM6B, Tb[1] |
| 605093 | y[1] w[*]; P{y[+t7.7] w[+mC]=20XUAS-pAce.x2}attP40; TM3, Sb[1]/TM6B, Tb[1] |
| 605094 | y[1] w[*]; P{y[+t7.7] w[+mC]=20XUAS-pAce.x2}attP2 |
| 605095 | w[*]; P{y[+t7.7] w[+mC]=UAS-sytjGCaMP7f.x2}attP2 |
| 605096 | y[1] w[*]; P{y[+t7.7] w[+mC]=20XQUAS-Ace2N-2AA-mNeon2}attP2 |
| 605097 | y[1] w[*]; P{y[+t7.7] w[+mC]=13XLexAop-Ace2N-2AA-mNeon2.x2}attP2 |
| 605098 | y[1] w[*]; P{y[+t7.7] w[+mC]=13XlexAop-Ace2N-2AA-mNeon2,20XUAS-pAce}attP40; MKRS/TM6B, Tb[1] |
| 605099 | y[1] w[*]; P{y[+t7.7] w[+mC]=20XUAS-pAce,13XlexAop-Ace2N-2AA-mNeon2}attP40 |
| 605100 | y[1] w[*]; P{y[+t7.7] w[+mC]=13XLexAop-jGCaMP8m}attP40 |
| 605101 | y[1] w[*]; P{y[+t7.7] w[+mC]=13XLexAop-jGCaMP8m}attP2 |
| 605102 | y[1] w[*]; PBac{y[+mDint2] w[+mC]=13XLexAop-jGCaMP8m}VK00027 |
| 605103 | y[1] M{y[+mDint]=vas-Bxb1.nanos3'UTR.RFP-}ZH-2A w[*]; P{RFP[mCh.2xr4] y[+t7.7] w[+mC]=attP(Bxb1),13XLexAop-jGCaMP8m}attP40/CyO |
| 605104 | y[1] M{y[+mDint]=vas-Bxb1.nanos3'UTR.RFP-}ZH-2A w[*]; P{GFP[CFP.2xr4] y[+t7.7]=13XLexAop-jGCaMP8m,attB(Bxb1)}attP40 |
| 605105 | y[1] M{y[+mDint]=vas-Bxb1.nanos3'UTR.RFP-}ZH-2A w[*]; P{RFP[mCh.2xr4] y[+t7.7] w[+mC]=attP(Bxb1),R14C08-GAL4.DBD}attP2/TM6B, Tb[1] |
| 605106 | y[1] M{y[+mDint]=vas-Bxb1.nanos3'UTR.RFP-}ZH-2A w[*]; P{RFP[mCh.2xr4] y[+t7.7] w[+mC]=attP(Bxb1),R19F09-GAL4.DBD}attP2/TM6B, Tb[1] |
| 605107 | y[1] M{y[+mDint]=vas-Bxb1.nanos3'UTR.RFP-}ZH-2A w[*]; P{RFP[mCh.2xr4] y[+t7.7] w[+mC]=attP(Bxb1),R25D01-GAL4.DBD}attP2 |
| 605108 | y[1] M{y[+mDint]=vas-Bxb1.nanos3'UTR.RFP-}ZH-2A w[*]; P{RFP[mCh.2xr4] y[+t7.7] w[+mC]=attP(Bxb1),R13F04-GAL4.DBD}attP2 |
| 605109 | y[1] M{y[+mDint]=vas-Bxb1.nanos3'UTR.RFP-}ZH-2A w[*]; P{RFP[mCh.2xr4] y[+t7.7] w[+mC]=attP(Bxb1),R27G01-GAL4.DBD}attP2/TM6B, Tb[1] |
| 605110 | y[1] M{y[+mDint]=vas-Bxb1.nanos3'UTR.RFP-}ZH-2A w[*]; P{RFP[mCh.2xr4] y[+t7.7] w[+mC]=attP(Bxb1),R52H01-GAL4.DBD}attP2/TM6B, Tb[1] |
| 605111 | y[1] M{y[+mDint]=vas-Bxb1.nanos3'UTR.RFP-}ZH-2A w[*]; P{RFP[mCh.2xr4] y[+t7.7] w[+mC]=attP(Bxb1),R24E12-GAL4.DBD}attP2/TM6B, Tb[1] |
| 605112 | y[1] M{y[+mDint]=vas-Bxb1.nanos3'UTR.RFP-}ZH-2A w[*]; P{RFP[mCh.2xr4] y[+t7.7] w[+mC]=attP(Bxb1),R30E11-GAL4.DBD}attP2 |
| 605113 | y[1] M{y[+mDint]=vas-Bxb1.nanos3'UTR.RFP-}ZH-2A w[*]; P{GFP[CFP.2xr4] y[+t7.7] w[+mC]=R25D01-p65.AD.H,attB(Bxb1)}attP2/TM6B, Tb[1] |
| 605114 | y[1] M{y[+mDint]=vas-Bxb1.nanos3'UTR.RFP-}ZH-2A w[*]; P{GFP[CFP.2xr4] y[+t7.7] w[+mC]=R19F09-p65.AD.H,attB(Bxb1)}attP2/TM6B, Tb[1] |
| 605115 | y[1] M{y[+mDint]=vas-Bxb1.nanos3'UTR.RFP-}ZH-2A w[*]; P{GFP[CFP.2xr4] y[+t7.7] w[+mC]=R20A02-p65.AD.H,attB(Bxb1)}attP2/TM6B, Tb[1] |
| 605116 | y[1] M{y[+mDint]=vas-Bxb1.nanos3'UTR.RFP-}ZH-2A w[*]; P{GFP[CFP.2xr4] y[+t7.7] w[+mC]=R93D10-p65.AD.H,attB(Bxb1)}attP2/TM6B, Tb[1] |
| 605117 | y[1] M{y[+mDint]=vas-Bxb1.nanos3'UTR.RFP-}ZH-2A w[*]; P{GFP[CFP.2xr4] y[+t7.7] w[+mC]={R15B01-p65.AD.H,attB(Bxb1)}attP2 |
| 605118 | y[1] M{y[+mDint]=vas-Bxb1.nanos3'UTR.RFP-}ZH-2A w[*]; P{GFP[CFP.2xr4] y[+t7.7]=R52B07-p65.AD,attB(Bxb1)}attP2/TM6B, Tb[1] |
| 605119 | y[1] M{y[+mDint]=vas-Bxb1.nanos3'UTR.RFP-}ZH-2A w[*]; P{GFP[CFP.2xr4] y[+t7.7] w[+mC]=R53C03-p65.AD.H,attB(Bxb1)}attP2/TM6B, Tb[1] |
| 605120 | y[1] M{y[+mDint]=vas-Bxb1.nanos3'UTR.RFP-}ZH-2A w[*]; P{GFP[CFP.2xr4] y[+t7.7] w[+mC]=R30E08-p65.AD.H,attB(Bxb1)}attP2/TM6B, Tb[1] |
| 605121 | y[1] M{Act5C-Cas9.P.RFP-}ZH-2A w[1118] DNAlig4[169]; P{GFP[EGFP.2xr4] y[+t7.7]=p65AD,attB(Bxb1).G4HACK}attP2/TM6B, Tb[1] |
| 605122 | y[1] M{Act5C-Cas9.P.RFP-}ZH-2A w[1118] DNAlig4[169]; P{GFP[EGFP.2xr4] y[+t7.7]=QF2DBD,attB(Bxb1).G4HACK}attP2 |
| 605123 | y[1] M{y[+mDint]=vas-Bxb1.nanos3'UTR.RFP-}ZH-2A w[*]; P{RFP[mCh.2xr4] y[+t7.7] w[+mC]=attP(Bxb1),MB011B}attP2/TM6B, Tb[1] |
| 605124 | y[1] M{y[+mDint]=vas-Bxb1.nanos3'UTR.RFP-}ZH-2A w[*]; P{RFP[mCh.2xr4] y[+t7.7] w[+mC]=attP(Bxb1),MB077B}attP2/TM6B, Tb[1] |
| 605125 | y[1] M{y[+mDint]=vas-Bxb1.nanos3'UTR.RFP-}ZH-2A w[*]; P{RFP[mCh.2xr4] y[+t7.7] w[+mC]=attP(Bxb1),MB085C}attP2/TM6B, Tb[1] |
| 605126 | y[1] M{y[+mDint]=vas-Bxb1.nanos3'UTR.RFP-}ZH-2A w[*]; P{RFP[mCh.2xr4] y[+t7.7] w[+mC]=attP(Bxb1),MB110C}attP2/TM6B, Tb[1] |
| 605127 | y[1] M{y[+mDint]=vas-Bxb1.nanos3'UTR.RFP-}ZH-2A w[*]; P{RFP[mCh.2xr4] y[+t7.7] w[+mC]=attP(Bxb1),MB112C}attP2/TM6B, Tb[1] |
| 605128 | y[1] M{y[+mDint]=vas-Bxb1.nanos3'UTR.RFP-}ZH-2A w[*]; P{RFP[mCh.2xr4] y[+t7.7] w[+mC]=attP(Bxb1),MB262B}attP2 |
| 605129 | y[1] M{y[+mDint]=vas-Bxb1.nanos3'UTR.RFP-}ZH-2A w[*]; P{GFP[CFP.3xP3] y[+t7.7]=MB083C,attB(Bxb1)}attP2/TM6B, Tb[1] |
| 605130 | y[1] M{y[+mDint]=vas-Bxb1.nanos3'UTR.RFP-}ZH-2A w[*]; P{RFP[mCh.2xr4] y[+t7.7] w[+mC]=attP(Bxb1),R53C10-GAL4.DBD}attP2 |
| 605702 | y[1] w[*] P{y[+t7.7]=nanos-phiC31\int.NLS}X; P{RFP[mCh.3xP3] y[+t7.7] w[+mC]=attP(phiC31),R57C10-GAL4}attP40/CyO |
| 605703 | y[1] w[*] P{y[+t7.7]=nanos-phiC31\int.NLS}X; P{RFP[mCh.3xP3] y[+t7.7] w[+mC]=attP(phiC31),R57C10-P-lexA::p65}attP40/CyO |
| 605704 | y[1] w[*] P{y[+t7.7]=nanos-phiC31\int.NLS}X; P{GFP[EGFP.3xP3] y[+t7.7]=13XLexAop2-IVS-myr::GFP,attB(phiC31)}attP40/CyO |
| 605705 | y[1] w[*] P{y[+t7.7]=nanos-phiC31\int.NLS}X; P{RFP[mCh.3xP3] y[+t7.7] w[+mC]=attP(phiC31),13XLexAop2-IVS-myr::GFP,R57C10-P-lexA::p65}attP40/CyO |

| | |
|---|---|
| 605706 | y[1] w[*] P{y[+t7.7]=nanos-phiC31\int.NLS}X; P{GFP[EGFP.3xP3] y[+t7.7]=10XUAS-IVS-myr::tdTomato,R57C10-GAL4,attB(phiC31)}attP40/CyO |
| 605707 | y[1] M{y[+mDint]=vas-Bxb1.nanos3’UTR.RFP-}ZH-2A w[*]; P{RFP[mCh.2xr4] y[+t7.7] w[+mC]=attP(Bxb1),R57C10-GAL4}attP40 |
| 605708 | y[1] M{y[+mDint]=vas-Bxb1.nanos3’UTR.RFP-}ZH-2A w[*]; P{RFP[mCh.2xr4] y[+t7.7] w[+mC]=attP(Bxb1),R57C10-P-lexA::p65}attP40/CyO |
| 605709 | y[1] M{y[+mDint]=vas-Bxb1.nanos3’UTR.RFP-}ZH-2A w[*]; P{GFP[CFP.2xr4] y[+t7.7]=10XUAS-IVS-myr::tdTomato,attB(Bxb1)}attP40/CyO |
| 605710 | y[1] M{y[+mDint]=vas-Bxb1.nanos3’UTR.RFP-}ZH-2A w[*]; P{GFP[CFP.2xr4] y[+t7.7]=13XLexAop2-IVS-myr::GFP,attB(Bxb1)}attP40/CyO |
| 605711 | y[1] M{y[+mDint]=vas-Bxb1.nanos3’UTR.RFP-}ZH-2A w[*]; P{GFP[CFP.2xr4] y[+t7.7]=10XUAS-IVS-myr::tdTomato,R57C10-GAL4,attB(Bxb1)}attP40/CyO |
| 605712 | y[1] M{y[+mDint]=vas-Bxb1.nanos3’UTR.RFP-}ZH-2A w[*]; P{RFP[mCh.2xr4] y[+t7.7] w[+mC]=attP(Bxb1),R57C10-GAL4}attP2 |
| 605713 | y[1] M{y[+mDint]=vas-Bxb1.nanos3’UTR.RFP-}ZH-2A w[*]; P{GFP[CFP.2xr4] y[+t7.7]=R57C10-GAL4,attB(Bxb1)}attP2 |
| 605714 | y[1] M{vas-NLS-TP901-1.nos3’UTR.RFP-.GFP-}ZH-2A w[*]; P{GFP[CFP.TpnC41C] y[+t7.7]=10XUAS-IVS-myr::tdTomato,R57C10-GAL4,attB(TP901-1)}attP40/CyO |
| 605715 | y[1] M{Act5C-Cas9.P.RFP-}ZH-2A w[1118] DNAlig4[169]; P{RFP[mCh.3xP3] y[+t7.7] w[+mC]=AD1(Cas9),13XLexAop2-IVS-myr::GFP,R57C10-P-lexA::p65}attP40/CyO |
| 605716 | y[1] M{Act5C-Cas9.P.RFP-}ZH-2A w[1118] DNAlig4[169]; P{GFP[EGFP.3xP3] y[+t7.7]=10XUAS-IVS-myr::tdTomato,R57C10-GAL4,AD2(Cas9)}attP40/CyO |
| 605717 | y[1] w[*] P{y[+t7.7]=nanos-phiC31\int.NLS}X; P{GFP[EGFP.3xP3] y[+t7.7]=13XLexAop2-IVS-myr::GFP,attB-CA(phiC31)}attP40/CyO |
| 605718 | y[1] w[*] P{y[+t7.7]=nanos-phiC31\int.NLS}X; P{GFP[EGFP.3xP3] y[+t7.7]=13XLexAop2-IVS-myr::GFP,attB-AC(phiC31)}attP40/CyO |
| 605719 | y[1] w[*] P{y[+t7.7]=nanos-phiC31\int.NLS}X; P{RFP[mCh.3xP3] y[+t7.7] w[+mC]=attP(phiC31),R82C10-lexA}attP40/CyO |
| 605720 | w[1118]; nub[1] Adc[b-1] sna[Sco] lt[1] stw[3]/CyO; P{y[+t7.7] w[+mC]=MB083C,MB011B.x2}attP2/TM6B, Tb[1] |
| 605721 | w[*]; P{y[+t7.7] w[+mC]=MB011B,MB110C}attP2/TM6B, Tb[1] |
| 605722 | w[1118]; nub[1] Adc[b-1] sna[Sco] lt[1] stw[3]/CyO; P{y[+t7.7] w[+mC]=MB110C,MB011B}attP2/TM6B, Tb[1] |
| 605723 | w[1118]; nub[1] Adc[b-1] sna[Sco] lt[1] stw[3]/CyO; P{y[+t7.7] w[+mC]=MB083C,MB011B,R52B07-GAL4.DBD}attP2 PBac{y[+mDint2] w[+mC]=R52H01-p65.AD}VK00027 |
| 605724 | w[1118]; nub[1] Adc[b-1] sna[Sco] lt[1] stw[3]/CyO; P{y[+t7.7] w[+mC]=MB083C,MB011B,MB262B}attP2/TM6B, Tb[1] |
| 605725 | w[*]; P{y[+t7.7] w[+mC]=MB262B,MB083C,MB011B.x2}attP2/TM6B, Tb[1] |
| 605728 | y[1] M{y[+mDint]=vas-Bxb1.nanos3’UTR.RFP-}ZH-2A w[*]; P{RFP[mCh.2xr4] GFP[CFP.TpnC41C] y[+t7.7]=AD1(Bxb1),R52B07-GAL4.DBD,AD2(TP901-1)}attP2/TM6B, Tb[1] |
| 605729 | y[1] M{y[+mDint]=vas-Bxb1.nanos3’UTR.RFP-}ZH-2A w[*]; P{RFP[mCh.2xr4] GFP[CFP.TpnC41C] y[+t7.7]=AD1(Bxb1),R94B10-GAL4.DBD,AD2(TP901-1)}attP2/TM6B, Tb[1] |
| 605730 | y[1] M{y[+mDint]=vas-Bxb1.nanos3’UTR.RFP-}ZH-2A w[*]; P{RFP[mCh.2xr4] GFP[CFP.TpnC41C] y[+t7.7]=AD1(Bxb1),R15B01-GAL4.DBD,AD2(TP901-1)}attP2 |
| 605731 | y[1] M{vas-NLS-TP901-1.nanos3’UTR.RFP-.GFP-}ZH-2A w[*]; P{RFP[mCh.TpnC41C] GFP[EGFP.3xP3] y[+t7.7] w[+m*]=AD1(TP901-1),R52H01-p65.AD,AD2(phiC31)}attP2 |
| 605732 | y[1] M{vas-NLS-TP901-1.nanos3’UTR.RFP-.GFP-}ZH-2A w[*]; P{RFP[mCh.TpnC41C] GFP[EGFP.3xP3] y[+t7.7] w[+m*]=AD1(TP901-1),R52G04-p65.AD,AD2(phiC31)}attP2 |
| 605733 | y[1] M{vas-NLS-TP901-1.nanos3’UTR.RFP-.GFP-}ZH-2A w[*]; P{RFP[mCh.TpnC41C] GFP[EGFP.3xP3] y[+t7.7] w[+m*]=AD1(TP901-1),R14C08-p65.AD,AD2(phiC31)}attP2 |
| 605734 | y[1] w[*] P{y[+t7.7]=nanos-phiC31\int.NLS}X; P{RFP[mCh.3xP3] GFP[CFP.2xr4] y[+t7.7]=AD1(phiC31),20xUAS-pAce,AD2(Bxb1)}attP2 |
| 605735 | w[*]; P{y[+t7.7] w[+mC]=MB083C,MB011B,MB112C.x2}attP2/TM6B, Tb[1] |
| 605736 | w[1118]; nub[1] Adc[b-1] sna[Sco] lt[1] stw[3]/CyO; P{y[+t7.7] w[+mC]=MB112C,MB083C,MB011B}attP2/TM6B, Tb[1] |
| 605737 | w[*]; P{y[+t7.7] w[+mC]=MB262B,MB077B}attP2/TM6B, Tb[1] |
| 605738 | y[1] w[*] P{y[+t7.7]=nanos-phiC31\int.NLS}X; P{GFP[EGFP.3xP3] y[+t7.7] w[+m*]=10XUAS-IVS-myr::tdTomato,AD2(phiC31)}attP40 |
| 605739 | y[1] w[*] P{y[+t7.7]=nanos-phiC31\int.NLS}X; P{RFP[DsRed.3xP3] y[+t7.7]=13XLexAop2-mCD8::GFP,AD2-GG(phiC31)}attP40/CyO |
| 605740 | y[1] w[*] P{y[+t7.7]=nanos-phiC31\int.NLS}X; PBac{GFP[EGFP.3xP3] y[+mDint2] w[+m*]=20XUAS-Ace2N-2AA-mNeon2,AD2(phiC31)}VK00027 |
| 605741 | w[1118]; P{y[+t7.7] w[+mC]=UAS-sytjGCaMP7f,MB083C,MB011B}attP2 |
| 606065 | w[*]; P{y[+t7.7] w[+mC]=R82C10-lexA.x2,13XlexAop-VARNAM2}attP40/CyO; P{y[+t7.7] w[+mC]=R52B07-GAL4.DBD}attP2 PBac{y[+mDint2] w[+mC]=20XUAS-Ace2N-2AA-mNeon2,R52H01-p65.AD}VK00027 |
| 606126 | y[1] M{Act5C-Cas9.P.RFP-}ZH-2A w[1118] DNAlig4[169]; P{GFP[EGFP.3xP3] RFP[mCh.3xP3] y[+t7.7]=SuRe.AD1(phiC31)1}attP40/CyO |
| 606127 | y[1] M{Act5C-Cas9.P.RFP-}ZH-2A w[1118] DNAlig4[169]; P{GFP[EGFP.3xP3] RFP[mCh.3xP3] y[+t7.7]=SuRe.AD1(phiC31)1}attP2 |
| 606128 | y[1] M{Act5C-Cas9.P.RFP-}ZH-2A w[1118] DNAlig4[169]; PBac{GFP[EGFP.3xP3] RFP[mCh.3xP3] y[+mDint2]=SuRe.AD1(phiC31)1}VK00027/TM3, Sb[1] |
| 606129 | y[1] M{Act5C-Cas9.P.RFP-}ZH-2A w[1118] DNAlig4[169]; P{RFP[DsRed.3xP3] GFP[EGFP.3xP3] y[+t7.7] w[+mC]=SuRe.AD2(phiC31)1}attP40/CyO |
| 606130 | y[1] M{Act5C-Cas9.P.RFP-}ZH-2A w[1118] DNAlig4[169]; P{RFP[DsRed.3xP3] GFP[EGFP.3xP3] y[+t7.7] w[+mC]=SuRe.AD2(phiC31)1}attP2 |
| 606131 | y[1] M{Act5C-Cas9.P.RFP-}ZH-2A w[1118] DNAlig4[169]; PBac{RFP[DsRed.3xP3] GFP[EGFP.3xP3] y[+mDint2] w[+mC]=SuRe.AD2(phiC31)1}VK00027 |
| 606132 | y[1] M{Act5C-Cas9.P.RFP-}ZH-2A w[1118] DNAlig4[169]; P{GFP[EGFP.2xr4] RFP[mCh.2xr4] y[+t7.7]=SuRe.AD1(Bxb1)1}attP40 |
| 606133 | y[1] M{Act5C-Cas9.P.RFP-}ZH-2A w[1118] DNAlig4[169]; P{GFP[EGFP.2xr4] RFP[mCh.2xr4] y[+t7.7]=SuRe.AD1(Bxb1)1}attP2/TM6B, Tb[1] |
| 606134 | y[1] M{Act5C-Cas9.P.RFP-}ZH-2A w[1118] DNAlig4[169]; PBac{GFP[EGFP.2xr4] RFP[mCh.2xr4] y[+mDint2]=SuRe.AD1(Bxb1)1}VK00027/TM6B, Tb[1] |
| 606135 | y[1] M{Act5C-Cas9.P.RFP-}ZH-2A w[1118] DNAlig4[169]; P{RFP[DsRed.2xr4] GFP[CFP.2xr4] y[+t7.7]=SuRe.AD2(Bxb1)1}attP40/CyO |

| | |
|---|---|
| 606136 | y[1] M{Act5C-Cas9.P.RFP-}ZH-2A w[1118] DNAlig4[169]; P{RFP[DsRed.2xr4] GFP[CFP.2xr4] y[+t7.7]=SuRe.AD2(Bxb1)1}attP2 |
| 606137 | y[1] M{Act5C-Cas9.P.RFP-}ZH-2A w[1118] DNAlig4[169]; P{RFP[DsRed.2xr4] GFP[CFP.3xP3] y[+t7.7]=SuRe.AD2(Bxb1)2}attP40/CyO |
| 606138 | y[1] M{Act5C-Cas9.P.RFP-}ZH-2A w[1118] DNAlig4[169]; P{RFP[DsRed.2xr4] GFP[CFP.3xP3] y[+t7.7]=SuRe.AD2(Bxb1)2}attP2 |
| 606139 | y[1] M{Act5C-Cas9.P.RFP-}ZH-2A w[1118] DNAlig4[169]; PBac{RFP[DsRed.2xr4] GFP[CFP.3xP3] y[+mDint2]=SuRe.AD2(Bxb1)2}VK00027 |
| 606140 | y[1] M{Act5C-Cas9.P.RFP-}ZH-2A w[1118] DNAlig4[169]; P{GFP[EGFP.TpnC41C] RFP[mCh.TpnC41C] y[+t7.7]=SuRe.AD1(TP901-1)1}attP40/CyO |
| 606141 | y[1] M{Act5C-Cas9.P.RFP-}ZH-2A w[1118] DNAlig4[169]; P{GFP[EGFP.TpnC41C] RFP[mCh.TpnC41C] y[+t7.7]=SuRe.AD1(TP901-1)1}attP2 |
| 606142 | y[1] M{Act5C-Cas9.P.RFP-}ZH-2A w[1118] DNAlig4[169]; P{RFP[DsRed.TpnC41C] GFP[CFP.TpnC41C] y[+t7.7]=SuRe.AD2(TP901-1)1}attP40/CyO |
| 606143 | y[1] M{Act5C-Cas9.P.RFP-}ZH-2A w[1118] DNAlig4[169]; P{RFP[DsRed.TpnC41C] GFP[CFP.TpnC41C] y[+t7.7]=SuRe.AD2(TP901-1)1}attP2 |
| 606144 | y[1] M{y[+mDint]=vas-Bxb1.nanos3'UTR.RFP-}ZH-2A w[*]; P{RFP[mCh.2xr4] y[+t7.7] w[+mC]=AD1(Bxb1),MB083C,MB011B}attP2/TM6B, Tb[1] |
| 606145 | y[1] M{y[+mDint]=vas-Bxb1.nanos3'UTR.RFP-}ZH-2A w[*]; P{RFP[mCh.2xr4] y[+t7.7] w[+mC]=AD1(Bxb1),13XLexAop2-mCD8::GFP,R57C10-P-lexA::p65}attP40/CyO |
| 606154 | w[*]; P{y[+t7.7] w[+mC]=R82C10-lexA.x2,13XlexAop-Ace2N-2AA-mNeon2}attP40/CyO; P{y[+t7.7] w[+mC]=R52B07-GAL4.DBD}attP2 PBac{y[+mDint2] w[+mC]=20XUAS-pAce, R52H01-p65.AD}VK00027 |
